# Genome-wide macroevolutionary signatures of key innovations in butterflies colonizing new host plants

**DOI:** 10.1101/2020.07.08.193086

**Authors:** Rémi Allio, Benoit Nabholz, Stefan Wanke, Guillaume Chomicki, Oscar A. Pérez-Escobar, Adam M. Cotton, Anne-Laure Clamens, Gaël J. Kergoat, Felix A.H. Sperling, Fabien L. Condamine

**Author notes:** Correspondence Rémi Allio, Fabien L. Condamine.

## Abstract

The exuberant proliferation of herbivorous insects is attributed to their associations with plants. Despite abundant studies on insect-plant interactions, we do not know whether host-plant shifts have impacted both genomic adaptation and species diversification over geological times. We show that the antagonistic insect-plant interaction between swallowtail butterflies and the highly toxic birthworts began 55 million years ago in Beringia, followed by several major ancient host-plant shifts. This evolutionary framework provides a unique opportunity for repeated tests of genomic signatures of macroevolutionary changes and estimation of diversification rates across their phylogeny. We find that host-plant shifts in butterflies are associated with both genome-wide adaptive molecular evolution (more genes under positive selection) and repeated bursts of speciation rates, contributing to an increase in global diversification through time. Our study links ecological changes, genome-wide adaptations and macroevolutionary consequences, lending support to the importance of ecological interactions as evolutionary drivers over long time periods.

---

Plants and phytophagous insects constitute most of the documented species of terrestrial organisms. To explain their staggering diversity, Ehrlich and Raven^1^ proposed a model in which a continual arms race of attacks by herbivorous insects and new defences by their host plants is linked to species diversification via the creation of new adaptive zones, later termed the ‘escape-and-radiate’ model^2^. Study of insect-plant interactions has progressed tremendously since then through focus on chemistry^3^, phylogenetics^4, 5^, and genomics^6–9^. Divergence of key gene families^7–10^ and high speciation rates^11–13^ have been identified after host-plant shifts, with one example linking duplication of key genes to the ability to feed on new plants and increase diversification^7^. However, a major knowledge gap lies in our understanding of the evolutionary linkages and drivers of host-plant shifts, genome-wide signatures of adaptations, and processes of species diversification^14^.

Here we address this gap with an emblematic group that was instrumental in Ehrlich & Raven’s model - the swallowtail butterflies (Lepidoptera: Papilionidae). First, we created an extensive phylogenetic dataset including 7 genetic markers for 71% of swallowtail species diversity (408 of ∼570 described species, *Methods*). Second, we compiled host-plant preferences for each swallowtail species in the dataset. Their caterpillars feed on diverse flowering-plant families, and a third of swallowtail species are specialized on the flowering plant family Aristolochiaceae (birthworts), which is one of the most toxic plant groups and carcinogenic to many organisms^15, 16^. Phylogenetic estimates of ancestral host-plant preferences indicate that Aristolochiaceae were either the foodplant of ancestral Papilionidae^17^ or were colonized twice^18^, suggesting an ancient and highly conserved association with Aristolochiaceae throughout swallowtail evolution. Using a robust and newly reconstructed time-calibrated phylogeny (Supplementary Figs. 1-3), we have traced the evolutionary history of food-plant use and infer that the family Aristolochiaceae was the ancestral host for Papilionidae (Fig. 1; relative probabilities = 0.915, 0.789, and 0.787 with three models, Supplementary Figs. 4, 5). We further show that the genus *Aristolochia* was the ancestral host-plant, as almost all Aristolochiaceae-associated swallowtails feed on *Aristolochia* (Supplementary Fig. 6). Across the swallowtail phylogeny, we recover only 14 host-plant shifts at the family level (14 nodes out of 407; Supplementary Figs. 4, 5), suggesting strong evolutionary host-plant conservatism.

**Fig. 1.**
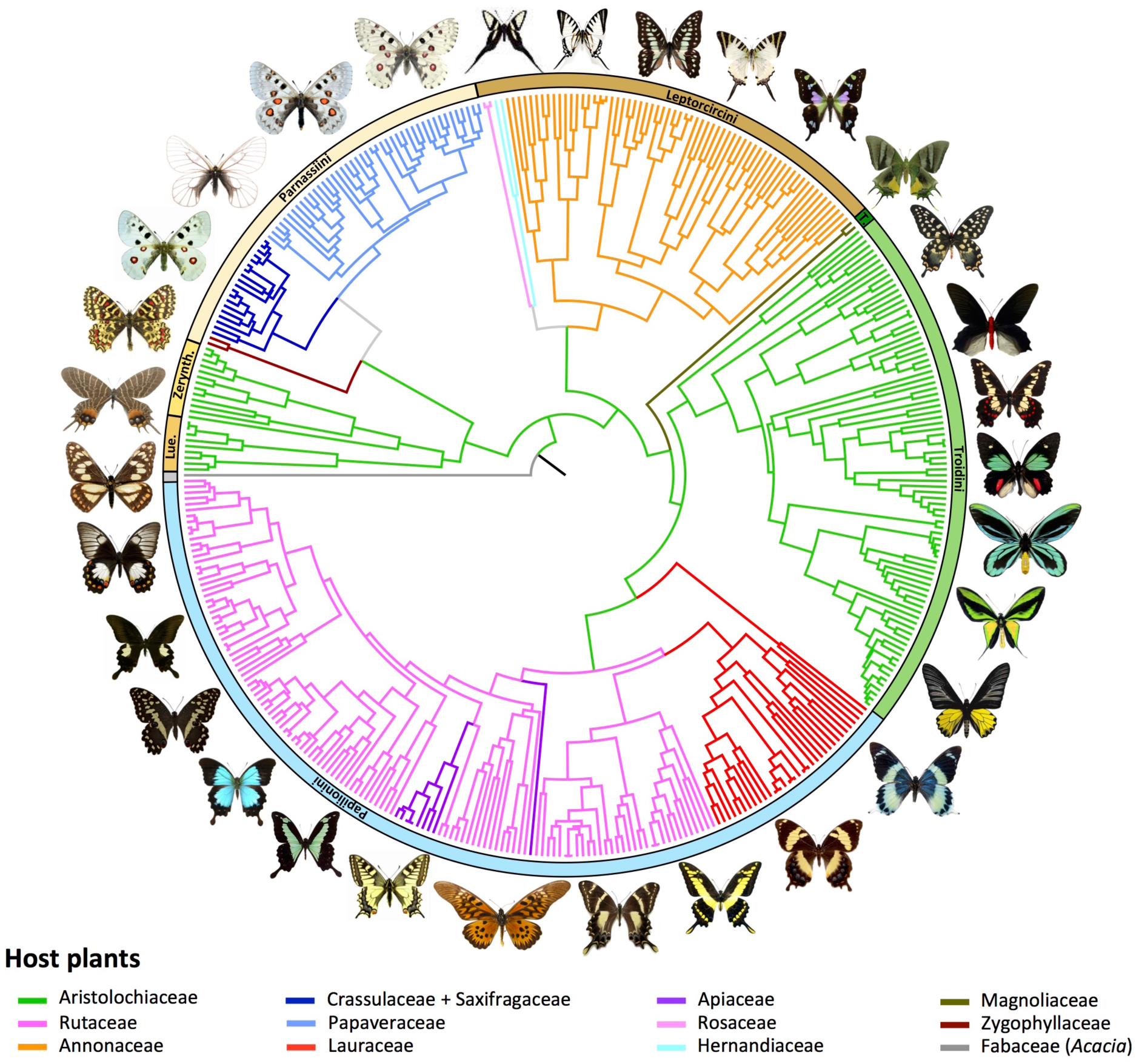
Evolution of host-plant association through time shows strong host-plant conservatism across swallowtail butterflies. Phylogenetic relationships of swallowtail butterflies, with coloured branches mapping the evolution of host-plant association, as inferred by a maximum-likelihood model (Supplementary Figs. 4, 6). Additional analyses with two other maximum-likelihood and Bayesian models inferred the same host-plant associations across the phylogeny (Supplementary Fig. 5). Lue. = Luehdorfiini, Zerynth. = Zerynthiini, and T. = Teinopalpini.

With the ancestor of swallowtails feeding on birthworts, evidence for synchronous temporal and geographical origins further links the genus *Aristolochia* and the family Papilionidae and supports the ‘escape and radiate’ model. Reconstructions of co-phylogenetic history for other insect-plant antagonistic interactions have shown either synchronous diversification^5^ or herbivore diversification lagging behind that of their host plants^4, 19^. We assembled a molecular dataset for ∼45% of the species diversity of Aristolochiaceae (247 of ∼550 described species; *Methods*) and reconstructed their phylogeny (Supplementary Fig. 7). Divergence time estimates indicate highly synchronous radiation by Papilionidae (55.4 million years ago [Ma], 95% credibility intervals: 47.8-71.0 Ma) and *Aristolochia* (55.5 Ma, 95% credibility intervals: 39.2-72.8 Ma) since the early Eocene (Fig. 2; Supplementary Figs. 3, 8, 9). This result is robust to known biases in inferring divergence times, with slightly older ages inferred for both groups when using more conservative priors on clade ages (Supplementary Fig. 9). Such temporal congruence between *Aristolochia* and Papilionidae raises the question of whether both clades had similar geographical origins and dispersal routes. To characterize the macroevolutionary patterns of the *Aristolochia*/Papilionidae arms-race in space, we assembled two datasets of current geographic distributions for all species included in the phylogenies of both Aristolochiaceae and Papilionidae. We reconstructed the historical biogeography of both groups, taking into account palaeogeographical events throughout the Cenozoic (*Methods*). The results show that both Papilionidae and *Aristolochia* were ancestrally co-distributed throughout a region including West Nearctic, East Palearctic, and Central America in the early Eocene, when Asia and North America were connected by the Bering land bridge (Fig. 2, Supplementary Figs. 10, 11). This extraordinary combination of close temporal and spatial congruence provides strong evidence that Papilionidae and *Aristolochia* diversified concurrently through time and space until several swallowtail lineages shifted to new host-plant families in the middle Eocene.

**Fig. 2.**
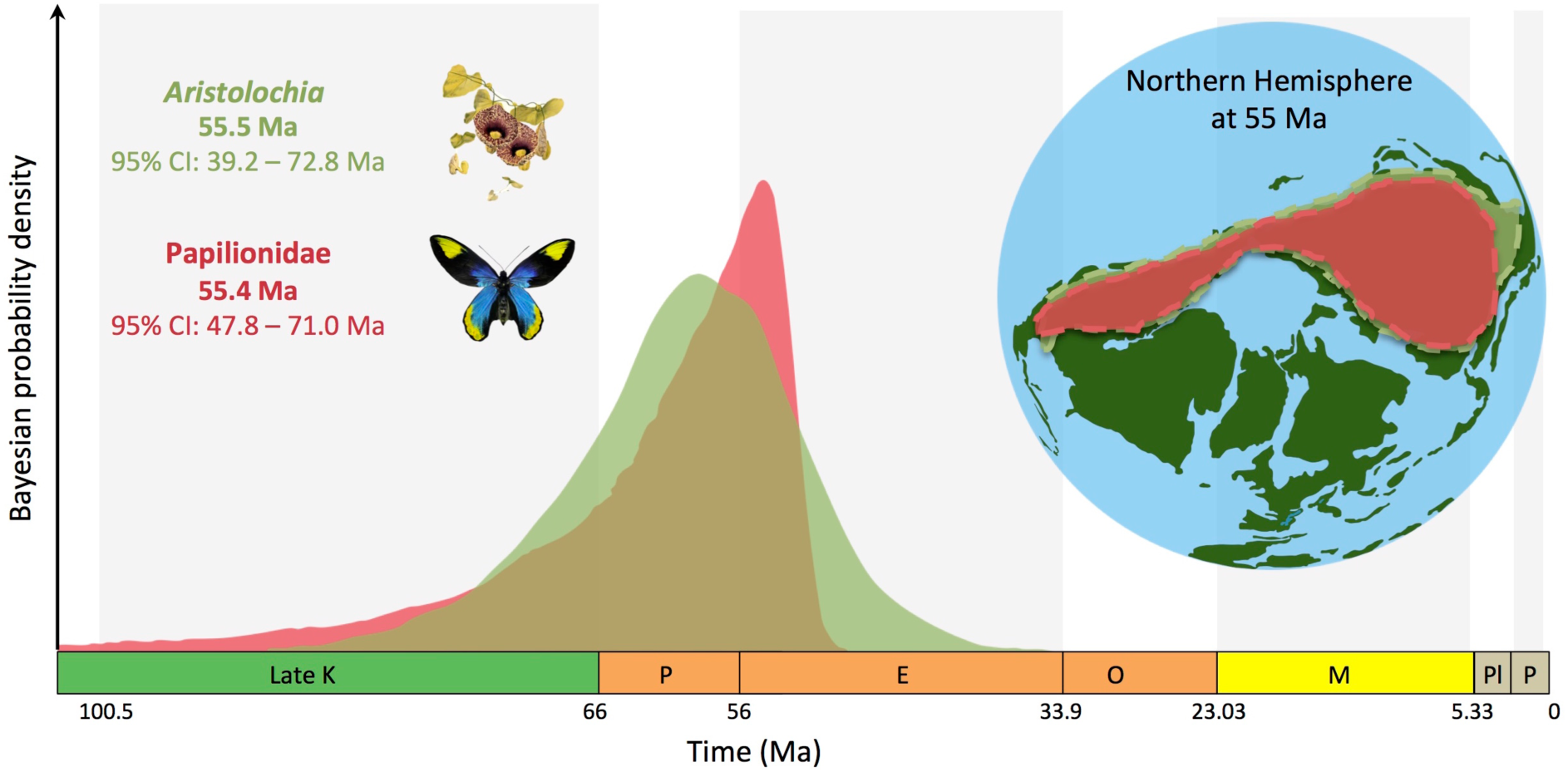
Synchronous temporal and geographic origin for swallowtails and birthworts. Bayesian molecular divergence times with exponential priors estimate an early Eocene origin (∼55 Ma) for both swallowtails and *Aristolochia* (alternatively, analyses with uniform prior estimated an origin around 67 Ma for swallowtails and 64 Ma for *Aristolochia*, Supplementary Figs. 3, 8, 9). Biogeographical maximum-likelihood models infer an ancestral area of origin comprising West Nearctic, East Palearctic and Central America for both swallowtails and birthworts (Supplementary Figs. 10, 11). K = Cretaceous, P = Palaeocene, E = Eocene, O = Oligocene, M = Miocene, Pl = Pliocene, and P = Pleistocene. Ma = million years ago.

Our ancestral state estimates and biogeographic analyses are consistent with a sustained arms race between *Aristolochia* and Papilionidae in the past 55 million years. According to the escape-and-radiate model, a host-plant shift should confer higher rates of species diversification for herbivores through the acquisition of novel resources to radiate into^1, 2^ and/or the lack of competitors (Aristolochiaceae-feeder swallowtails have almost no competitors^20^). We tested the hypothesis that increases of diversification rates occurred in swallowtail lineages that shifted to new host-plants. Applying a suite of birth-death models (*Methods*), we find evidence for (1) upshifts of diversification at host-plant shifts with trait-dependent birth-death models (Fig. 3a; Supplementary Figs. 12, 13, Supplementary Table 1), and (2) host-plant shifts contributing to a global increase through time with time-dependent birth-death models (Fig. 3b; Supplementary Figs. 14-16). Surprisingly, we do not observe the classical slowdown of diversification recovered in most phylogenies, often attributed to ecological limits and niche filling processes^21^. This sustained and increasing diversification during the Cenozoic may be explained by ecological opportunities not decreasing, due to a steady increase in host breadth for Papilionidae with new host-plant families colonized through time (Supplementary Fig. 17). Opening up new niches would allow continuous increase in diversification rates through time in a dynamic biotic environment, lending support to the primary role of ecological interactions in clade diversification over long timescales.

**Fig. 3.**
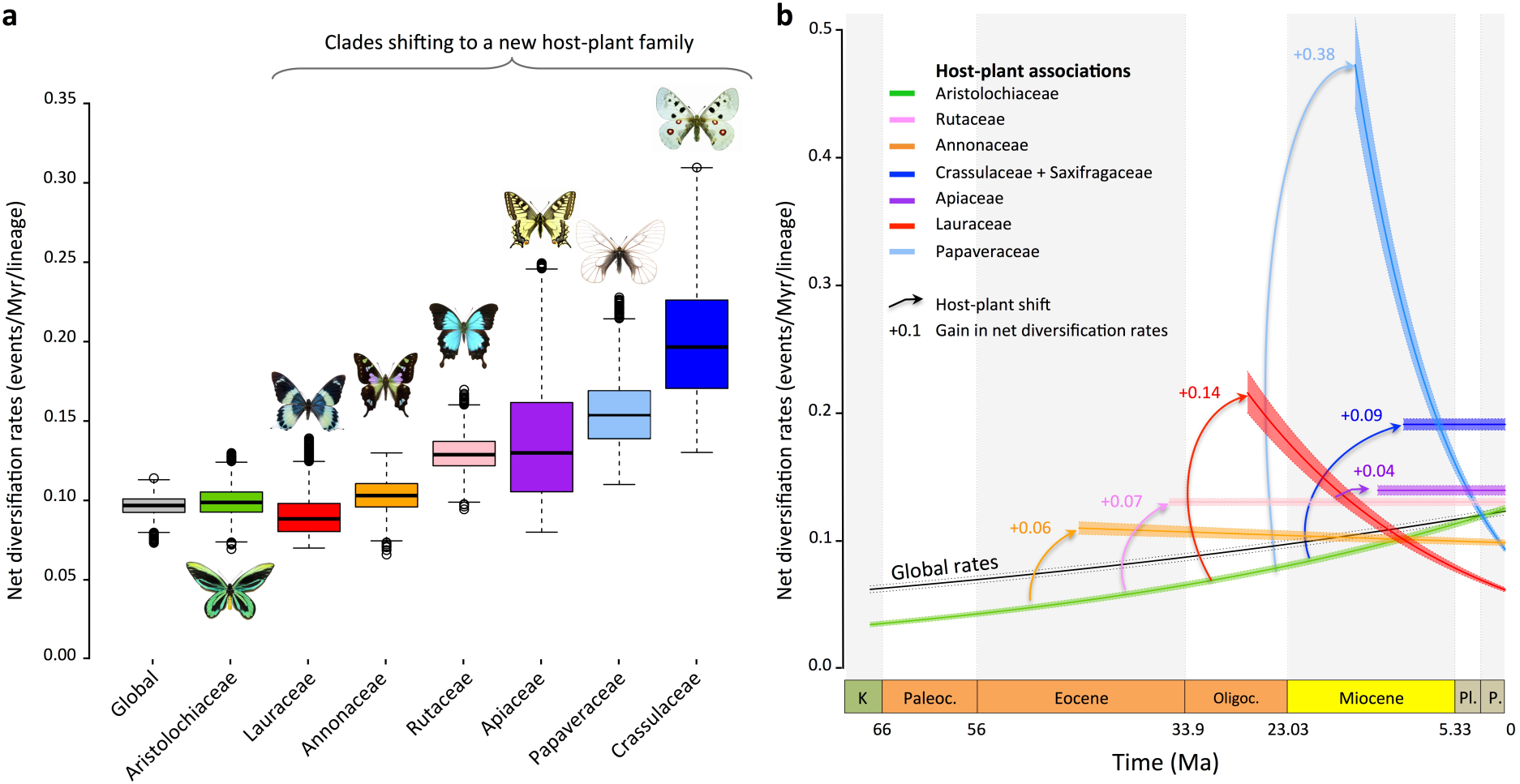
Host-plant shifts lead to repeated bursts in diversification rates and a sustained overall increase in diversification through time. **a**, Diversification tends to be higher for clades shifting to new host plants, as estimated by trait-dependent diversification models. Boxplots represent Bayesian estimates of net diversification rates for clades feeding on particular host plants (see also Supplementary Fig. 12). **b**, A global increase in diversification is recovered with birth-death models estimating time-dependent diversification (see also Supplementary Figs. 14, 15). Taking into account rate heterogeneity by estimating host-plant and clade-specific diversification indicates positive gains of net diversification after shifting to new host plants (see also Supplementary Fig. 13). K = Cretaceous, Paleoc. = Palaeocene, Oligoc. = Oligocene, Pl = Pliocene, P = Pleistocene, Ma = million years ago.

Key innovations are often considered to underlie ecological opportunities and/or evolutionary success^22^, particularly in the case of chemically mediated interactions between butterflies and their host-plants^7^. Studies on Papilionidae have provided strong examples of specific changes in key genes that confer new abilities to feed on toxic plants and allow host-plant shifts^23, 24^. Adaptations of swallowtails to their hosts have particularly been assessed through the study of cytochrome P450 monooxygenases (P450s), which have a major role in detoxifying secondary plant compounds. New P450s appear to arise in swallowtails that colonize new hosts to bypass toxic defences, providing survival and diversification on some but not all plants^9, 23, 25^. This supports the hypothesis that insect-plant interactions contributed to P450-gene family diversification, with P450s being key innovations that explain the evolutionary and ecological success of phytophagous insects^8, 9, 24, 26–28^. However, host-plant shifts not only alter single genes but may also influence unlinked genes^29^. Moreover, host-plant shifts can accompany changes of abiotic environment, which may in turn require further adaptation (new predators and/or competitors). But the macroevolutionary and genomic consequences of the evolutionary dynamics of host-plant shifts have not yet been demonstrated.

Relying on a genomic dataset comprising 45 genomes covering all swallowtail genera^30–33^, we asked whether there are any genomic signatures of positive selection caused by host-plant shifts within swallowtails. We performed a comparative genomic survey of molecular evolution to test whether there is a contrasting pattern of molecular adaptation between swallowtail lineages that shifted to new host plants compared to non-shifting lineages (*Methods*). We selected 14 phylogenetic branches representing a host-plant shift and 14 phylogenetic branches with no change as negative controls^34, 35^ (Fig. 4a). For a fair molecular comparison, each branch selected as a negative control was chosen to be as close as possible to a test branch representing a host-plant shift (i.e. sister groups, Supplementary Fig. 18). Among branches with host-plant shifts, 5 branches also had a shift in climate preference (represented by distributional changes from tropical to temperate conditions). Using a maximum-likelihood method, we estimated the ratio of non-synonymous substitutions (dN) other synonymous substitutions (dS) in all branches where a host-plant shift was identified relative to branches with no host-plant shift^36, 37^ (*Methods*). The dN/dS analyses on branches with host-plant shifts (combined or not with environmental shifts) showed more genome-wide molecular adaptations (i.e. more genes under positive selection, dN/dS > 1) in lineages shifting to a new plant family, although the difference was marginally non-significant (Fig. 4b, *P* = 0.0501 / 0.0345 for the two datasets, respectively, Wilcoxon rank-sum test, see *Methods* for the definition of the datasets). However, dN/dS analyses on branches with environmental shifts indicated a balanced number of genes under positive selection (Fig. 4c, *P* = 0.336 / 0.834 for the two datasets, respectively, Wilcoxon rank-sum test), suggesting a lower impact of environmental shifts than host-plant shifts. We then performed dN/dS analyses for branches with host-plant shifts only (not followed by environmental shifts) and found that swallowtail lineages shifting to a new host-plant family had significantly more genes under positive selection (4.41% / 3.64% of genes under positive selection for the two datasets, respectively) than non-shifting lineages (3.02% / 2.33% of genes under positive selection for the two datasets, respectively, Fig. 4d, *P* = 0.0071 / 0.0152 for the two datasets, respectively, Wilcoxon rank-sum test). We checked individually the gene alignments and performed sensitivity analyses that showed our results are not driven either by an excess of misaligned regions, nor missing data and GC-content variations among species (*Methods*; Supplementary Figs. 19-25). Surprisingly, the dual changes in climate and host-plant preferences did not spur molecular adaptation across swallowtail lineages (*P* = 1 / 0.517 for the two datasets, respectively, Wilcoxon rank-sum test) and even less than host-plant shifts only (*P* = 0.0327 / 0.147 for the two datasets, respectively, Wilcoxon rank-sum test; Fig. 3d). Although these genome-wide comparisons rely on a few branches (5 out of 14 which significantly differ from others, tested with 1000 random comparisons), no plausible hypothesis can explain this result that would require more in-depth work.

**Fig. 4.**
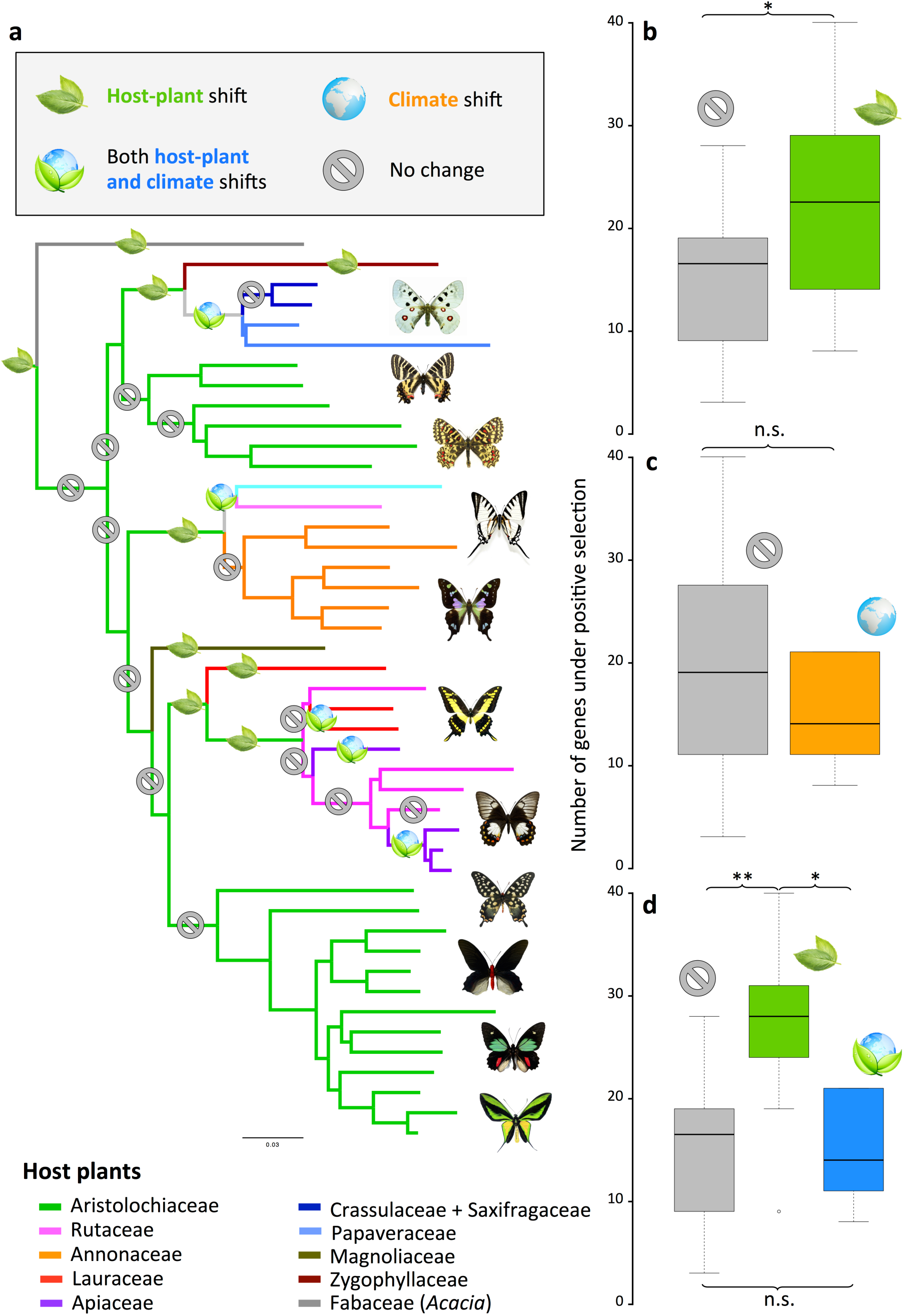
Host-plant shifts promote higher molecular adaptations. **a**, Genus-level phylogenomic tree displaying branches with and without host-plant shifts, on which genome-wide analyses of molecular evolution are performed. **b**, Number of genes under positive selection (dN/dS > 1) for swallowtail lineages shifting to new host-plant families (green) or not (grey). **c**, Number of genes under positive selection for swallowtail lineages undergoing climate shifts (orange) or not (grey). **d**, Number of genes under positive selection for swallowtail lineages shifting to new host plants (green), shifting both host plant and climate (blue) or not (grey). This demonstrates genome-wide signatures of adaptations in swallowtail lineages shifting to new host-plant families. Genes under positive selection did not contain over-or under-represented functional GO categories (Supplementary Table 2). n.s. = not significant (*P* > 0.05), * = *P* ≤ 0.05, ** = *P* ≤ 0.01.

We further studied the functional categories of positively selected genes by using gene ontology (GO) analyses (PANTHER and EggNOG; *Methods*). Applied to the high-quality genomes of *Papilio xuthus*^31^ and *Heliconius melpomene*^38^, we found that ∼70% of the genes are associated with a gene function, which suggests a gap of knowledge in insect gene function database. Among the annotated genes, we found that genes under positive selection along branches with host shifts did not contain over- or under-represented functional GO categories: 252 out of 1213 GO categories represented by genes under positive selection (*P* > 0.05, Fisher’s exact test after false discovery rate correction; Supplementary Table 2). These results support the hypothesis that genome-wide signatures of adaptations are associated with host-plant shifts, and encourage extending the long-held hypothesis that only changes in a single candidate family gene are enough to act as a key innovation for adaptation to new resources^7, 10^. Despite a weak signal, it is striking that host-plant shifts left stronger genome-wide signatures than were associated with changing climate preferences. This result further suggests that the success of phytophagous insects involved deeper adaptation to biotic interactions than for shifts in the abiotic environment.

Establishing linkages between ecological adaptations, genomic changes, and species diversification over geological timescales remains a tremendous challenge^14^ with, for instance, important limitations due to the lack of knowledge in functional gene annotations in insects. However, the successful development of powerful analytical tools in conjunction with the increasing availability of insect genomes and improvements in genomic analyses^39^ allow detecting more genes than the known genes involved in detoxification pathways playing a role in long-term relationships between plants and insects. This opens new research avenues for finding the functionality of genes involved in the adaptation and diversification of phytophagous insects. We hope that our study will help movement in that direction, and that it will provide interesting perspectives for future investigations of other model groups.

Over a half century ago, Ehrlich and Raven^1^ proposed that insect-plant interactions driven by diffuse co-evolution over long evolutionary periods can be a major source of terrestrial biodiversity. Applied to a widely appreciated case in the insect-plant interactions theory, our study reveals that genome-wide adaptive processes and corresponding macroevolutionary consequences are more pervasive than previously recognized in the diversification of herbivorous insects. Close relationships between insects and their larval host plants involve more adaptations than in just the gene families in detoxification pathways that were detected through antagonist interactions^39^, and show genomically wide-ranging co-evolutionary consequences^29, 40^. Hence, genome-wide macroevolutionary consequences of key adaptations in new insect-plant interactions may be a general feature of the co-evolutionary interactions that have generated Earth’s diversity.

## Acknowledgements

This project has received funding from the Marie Curie International Outgoing Fellow under the European Union’s Seventh Framework Programme (project BIOMME, agreement No. 627684), a PICS grant from the CNRS (project PASTA), an “Investissement d’Avenir” grant from the Agence Nationale de la Recherche (project CASMA, CEBA, ref. ANR-10-LABX-25-01), and the European Research Council (ERC) under the European Union’s Horizon 2020 research and innovation programme (project GAIA, agreement No. 851188) to F.L.C.; a Natural Sciences and Engineering Research Council of Canada (NSERC) Discovery Grant (RGPIN-2018-04920) to F.A.H.S.; and a German Research Foundation grant (WA 2461/9-1) to S.W. We are grateful to Sophie Dang, Troy Locke, and Corey Davis at the Molecular Biology Service Unit of the University of Alberta for their help, assistance, and advice on next-generation sequencing. The analyses benefited from the Montpellier Bioinformatics Biodiversity (MBB) platform services. Finally, we are grateful to Seth Bybee, Frédéric Delsuc, Claude dePamphilis, Krushnamegh Kunte, Conrad Labandeira, Harald Letsch, Sören Nylin, Timothy O’Hara, Susanne Renner and Chris Wheat for helpful comments and discussions on earlier drafts of the study.

## Author contributions

F.L.C. and F.A.H.S. designed and conceived the research. R.A. and F.L.C. assembled the phylogenetic data for swallowtail butterflies. S.W., O.A.P.E., G.C., F.L.C and R.A. assembled the phylogenetic data for birthworts. R.A. and F.L.C. analysed the phylogenetic data. R.A. and F.L.C. performed the ancestral states estimations. F.L.C. performed the diversification analyses. A.-L.C. and F.L.C. generated the genomic data. R.A. and B.N. assembled and analysed the genomic data. All authors contributed to the interpretation and discussion of results. R.A. and F.L.C. drafted the paper with substantial input from all authors.

## Methods

### Time-calibrated phylogeny of Papilionidae

We assembled a supermatrix dataset with available data extracted from GenBank as of May 2017 (most of which has been generated by our research group), using five mitochondrial genes (*COI*, *COII*, *ND1*, *ND5* and *rRNA 16S*) and two nuclear markers (*EF-1a* and *Wg*) for 408 Papilionidae species (∼71% of the total species diversity) and 20 outgroup species. We aligned the DNA sequences for each gene using MAFFT 7.110^41^ with default settings (E-INS-i algorithm), and the alignments were checked for codon stops and eventually refined by eye with Mesquite 3.1 (available at: www.mesquiteproject.org). The best-fit partitioning schemes and substitution models for phylogenetic analyses were determined with PartitionFinder 2.1.1^42^ using the *greedy* search algorithm and the Bayesian Information Criterion. All gene alignments were concatenated in a supermatrix, which is available in Figshare (see Data availability).

Phylogenetic relationships were estimated with both maximum likelihood (ML) and Bayesian inference. ML analyses were carried out with IQ-TREE 1.6.8^43^. We set the best-fit partitioning scheme and used ModelFinder to determine the best-fit substitution model for each partition^44^ and then estimated model parameters separately for every partition^45^ such that all partitions shared the same set of branch lengths, but we allowed each partition to have its own evolution rate. We performed 1,000 ultrafast bootstrap replicates to investigate nodal support across the topology, considering values ≥ 95 as strongly supported nodes^46^.

Estimating phylogenetic relationships for such a dataset is computationally intensive with Bayesian inference. The ML tree inferred with IQ-TREE was used as a starting tree for Bayesian inference as implemented in MrBayes 3.2.6^47^. Rather than using a single substitution model per molecular partition, we sampled across the entire substitution-model space^48^ using reversible-jump Markov Chain Monte Carlo (rj-MCMC). Two independent analyses with one cold chain and seven heated chains, each run for 50 million generations, sampled every 5,000 generations. Convergence and performance of Bayesian runs were evaluated using Tracer 1.7.1^49^, the average deviation of split frequencies (ADSF) between runs, the effective sample size (ESS) and the potential scale reduction factor (PSRF) values for each parameter. A 50% majority-rule consensus tree was built after conservatively discarding 25% of sampled trees as burn-in. Node support was evaluated with posterior probability considering values ≥ 0.95 as strong support^50^. All analyses were performed on the CIPRES Science Gateway computer cluster^51^, using BEAGLE^52^.

Dating inferences were performed using Bayesian relaxed-clock methods accounting for rate variation across lineages^53^. MCMC analyses implemented in BEAST 1.8.4^54^ were employed to approximate the posterior distribution of rates and divergences times and infer their credibility intervals. Estimation of divergence times relied on constraining clade ages through fossil calibrations. Swallowtail fossils are scarce, but five can unambiguously be attributed to the family. The oldest fossil occurrences of Papilionidae are the fossils †*Praepapilio colorado* and †*Praepapilio gracilis*^55^, both from the Green River Formation (Colorado, USA). The Green River Formation encompasses a 5 million-years period between ∼48.5 and 53.5 Ma, which falls within the Ypresian (47.8-56 Ma) in the early Eocene^56^. These fossils can be phylogenetically placed at the crown of the family as they share synapomorphies with all extant subfamilies^57, 58^, and have proven to be reliable calibration points for the crown group^12, 17, 33^. Two other fossils belong to Parnassiinae, whose systematic position was assessed using phylogenetic analyses based on both morphological and molecular data in a total-evidence approach^12^. The first is †*Thaites ruminiana*^59^, a compression fossil from limestone in the Niveau du gypse d’Aix Formation of France (Bouches-du-Rhône, Aix-en-Provence, France) within the Chattian (23.03–28.1 Ma) of the late Oligocene^60, 61^. †*Thaites* is sister to Parnassiini, and occasionally sister to Luehdorfiini + Zerynthiini^12^. Thus we constrained the crown age of Parnassiinae with a uniform distribution bounded by a minimum age of 23.03 Ma. The second is †*Doritites bosniaskii*^62^, an exoskeleton and compression fossil from Italy (Tuscany) from the Messinian (5.33–7.25 Ma, late Miocene)^61^. †*Doritites* is sister to *Archon* (Luehdorfiini^12^), in agreement with Carpenter^63^. The crown of Luehdorfiini was thus constrained for divergence time estimation using a uniform distribution bounded with 5.33 Ma. Absolute ages of geological formations were taken from the latest update of the geological time scale.

We used a conservative approach to applying calibration priors with the selected fossil constraints by setting uniform priors bounded with a minimum age equal to the youngest age of the geological formation where each fossil was found. All uniform calibration priors were set with an upper bound equal to the estimated age of angiosperms (150 Ma^64^), which is more than three times older than the oldest Papilionidae fossil. This upper age is intentionally set as ancient to allow exploration of potentially old ages for the clade. Since the fossil record of butterflies is incomplete and biased^65^, caution is needed in using these fossil calibrations (effect shown in burying beetles^66^).

After enforcing the fossil calibrations, we set the following settings and priors: a partitioned dataset (after the best-fitting PartitionFinder scheme) was analysed using the uncorrelated lognormal distribution clock model, with the mean set to a uniform prior between 0 and 1, and an exponential prior (lambda = 0.333) for the standard deviation. The branching process prior was set to a birth–death^67^ process, using the following uniform priors: the birth–death mean growth rate ranged between 0 and 10 with a starting value at 0.1, and the birth–death relative death rate ranged between 0 and 1 (starting value = 0.5). We performed four independent BEAST analyses for 100 million generations, sampled every 10,000th, resulting in 10,000 samples in the posterior distribution of which the first 2500 samples were discarded as burn-in. All analyses were performed on the CIPRES Science Gateway computer cluster^51^, using BEAGLE^52^. Convergence and performance of each MCMC run were evaluated using Tracer 1.7.1^49^ and the ESS for each parameter. We combined the four runs using LogCombiner 1.8.4^54^. A maximum-clade credibility (MCC) tree was reconstructed, with median ages and 95% credibility intervals (CI). The BEAST files generated for this study are available in Figshare (see Data availability).

### Estimating ancestral host-plant association

We inferred the temporal evolution of host-plant association up to the ancestral host plant(s) at the root of Papilionidae using three approaches: the ML implementation of the Markov k-state (Mk) model^68^, the ML Dispersal-Extinction-Cladogenesis (DEC) model^69^, and the Bayesian approach in BayesTraits^70^. These approaches require a time-calibrated tree and a matrix of character states (current host-plant preference) for each species in the tree. An extensive bibliographic survey was conducted to obtain primary larval host-plants at the family level^1, 71–74^. The host associations of species were categorized using the following twelve character states: (1) Annonaceae, (2) Apiaceae, (3) Aristolochiaceae, (4) Crassulaceae or Saxifragaceae (core Saxifragales), (5) Fabaceae, (6) Hernandiaceae, (7) Lauraceae; (8) Magnoliaceae, (9) Papaveraceae, (10) Rosaceae, (11) Rutaceae, and (12) Zygophyllaceae. The host-plant matrix of Papilionidae is available in Figshare (see Data availability).

Ancestral states for host-plant association were first reconstructed using the Mk model (one rate for all transitions between states) allowing any host shift to be equally probable. The Mk model does not allow multiple states for a species. The few species that use multiple host families were thus scored with the most frequent host association. The Mk model was performed with Mesquite 3.1 (available at: www.mesquiteproject.org). To estimate the support of any one character state over another, the most likely state was selected according to a decision threshold, such that if the log likelihoods between two states differ by two log-likelihood units, the one with lower likelihood is rejected^68^.

The DEC model was also used to reconstruct ancestral host-plant states^69, 75^. As the Mk model, we assumed that host-plant shifts occurred at equivalent probabilities between plant families and through time, which may not be true given that the host-plant families of Papilionidae did not originate at the same time (e.g. Aristolochiaceae originated around 108.07 Ma [95% credibility intervals: 81.01-132.66 Ma]^76^, and Annonaceae originated about 98.94 Ma [95% credibility intervals: 84.78-113.70 Ma]^76^). We used the estimated molecular ages of the different host-plant groups to constrain our inferences of ancestral host plants *a posteriori*. We preferred such an approach compared to a more constrained one in which the DEC model is informed with a matrix of host-plant appearances based on their estimated ages by implementing matrices of presence/absence of the character states through time (equivalent to the time-stratified palaeogeographic model, see below for inference of biogeographical history).

Finally, the Bayesian approach implemented in BayesTraits 3.0.1^70^ was performed to provide a cross-validation of ML analyses. This approach automatically detects shifts in rates of evolution for multistate data using rj-MCMC. Numbers of parameters and priors were set by default. We ran the rj-MCMC for 10 million generations and sampled states and parameters every 1,000 generations (burn-in of 10,000 generations). We specifically estimated ancestral states at 21 nodes as well as at the root of Papilionidae. For this analysis, we used a set of 100 trees randomly taken from the dating analysis to probe the robustness of our ancestral state estimation across topological uncertainty.

The results of these inferences determined the host-plant family(ies) that was (were) the most likely ancestral host(s) at the origin of Papilionidae, indicating *(i)* which plant phylogeny to reconstruct for studying the macroevolution of the arms race, and *(ii)* the evolution of ancestral host-plant association along the phylogeny to identify the tree branches where shifts occurred and test for genome-wide changes.

The Mk and BayesTraits models always inferred with high support (relative probability = 0.915 and 0.789, respectively) that Aristolochiaceae is the ancestral host plant at the crown of Papilionidae. With the unconstrained DEC model, we found that the ancestral host-plant preference for Papilionidae was always composed of Aristolochiaceae, but also included another family (either Fabaceae, Hernandiaceae or Zygophyllaceae, which are only fed upon by *Baronia*, *Lamproptera* and *Hypermnestra*, respectively). As the sister lineage to all other Papilionidae, *Baronia* is the only species that feeds on Fabaceae. More precisely, only one species of Fabaceae is consumed: *Vachellia cochliacantha* (formerly *Acacia cochliacantha*; recent changes in *Acacia* taxonomy^77^). However, *Vachellia* diverged from its sister clade in the Eocene, approximately 50 Ma, and diversified in the Miocene between 13 and 17 Ma^78^, which substantially postdate the origin of Papilionidae. Therefore this result suggests that Aristolochiaceae family represents the most likely candidate as the ancestral host-plant of Papilionidae. Hernandiaceae are consumed by *Lamproptera* (occasionally by *Papilio homerus*, *Graphium codrus*, *G. doson* and *G. empedovana*^73^). More precisely, the host plants of *Lamproptera* belong to the genus *Illigera*. This plant genus diverged from its sister genus 48 Ma^76^ and started diversifying 27 Ma^79^. The derived phylogenetic position of *Lamproptera* and the age of its use as a host plant make it very unlikely that Hernandiaceae could constitute the ancestral host plant for Papilionidae. Similarly, the family Zygophyllaceae is consumed by *Hypermnestra*, most specifically it feeds on the genus *Zygophyllum* in Central Asia. The genus *Zygophyllum* is not monophyletic, but Asian *Zygophyllum* appeared 19.6 Ma^80^. Applying the same rationale, we are able to discard Zygophyllaceae as a candidate ancestral host plant for Papilionidae. To further refine our ancestral host-plant estimates, we built a presence-absence matrix of plant families based on clade origins estimated in molecular dating studies. Thereby, the age of the different plants can be used to constrain the inference of ancestral host plants. Under such a constrained model, Aristolochiaceae is always recovered as the most likely ancestral host-plant for Papilionidae. It is also interesting that almost all Aristolochiaceae feeders have *Aristolochia* as host plants, and tests to determine which genus of Aristolochiaceae was originally consumed by Papilionidae showed that it was *Aristolochia*.

### Time-calibrated phylogeny of the ancestral host: the Aristolochiaceae

Estimation of ancestral host-plant relationships revealed that the family Aristolochiaceae was the ancestral host for Papilionidae. We refer to Aristolochiaceae in its traditional circumscription including the genera *Asarum*, *Saruma*, *Thottea* and *Aristolochia.* The Angiosperm Phylogeny Group^81^ proposes that Aristolochiaceae also includes the holoparasitic genera *Hydnora* and *Prosopanche* (Hydnoraceae), as well as the monotypic family Lactoridaceae from the Juan Fernandez Islands of Chile (*Lactoris fernandeziana*). The conclusion of APG^81^ is based on an online survey^82^ rather than on primary data and this is why we disagree with their argumentation as well as the resulting conclusion of APG given available resilient primary molecular phylogenomic data. However, arguments based on morphology and anatomy^83–86^, genetics^87–92^, molecular divergence time^76, 92^, and conservation considerations (Tod Stuessy, pers. comm. with S.W., July 2019) favour splitting them into four families: Aristolochiaceae (*Aristolochia* and *Thottea*), Asaraceae (*Asarum* and *Saruma*), Hydnoraceae (*Hydnora* and *Prosopanche*), and Lactoridaceae (*Lactoris*), collectively called the perianth-bearing Piperales. Therefore we extracted and assembled a supermatrix dataset with available data from GenBank for the perianth-bearing Piperales and its sister lineage, the perianth-less Piperales including Saururaceae and Piperaceae (as of May 2017, most of which has been generated by our research group). We obtained four chloroplast genes (*matK*, *rbcl*, *trnL*, *trnL-trnF*) and one nuclear marker (*ITS*) for 247 species of perianth-bearing Piperales (∼45% of the total species diversity^93^) and six outgroups from perianth-less Piperales. We could not include the two genera *Hydnora* and *Prosopanche* (Hydnoraceae) because available genetic data do not overlap those of perianth-bearing Piperales^87, 91, 94, 95^. We applied the same analytical procedure that we did for Papilionidae. DNA sequences for each gene were aligned using MAFFT 7.110^41^ with default settings (E-INS-i algorithm and Q-INS-I to take into account secondary structure). Resulting alignments were checked for codon stops and eventually refined by eye with Mesquite 3.1 (available at: www.mesquiteproject.org). The best-fit partitioning schemes and substitution models for phylogenetic analyses were determined with PartitionFinder 2.1.1^42^. All gene alignments were concatenated into a supermatrix; the final dataset is available in Figshare (see Data availability).

Phylogenetic relationships were estimated with Bayesian inference as implemented in MrBayes 3.2.6^47^. Rather than using a single substitution model per molecular partition, we sampled across the entire substitution-model space^48^ using rj-MCMC. Two independent analyses with one cold chain and seven heated chains, each were run for 50 million generations, sampled every 5,000 generations. Convergence and performance of Bayesian runs were evaluated using Tracer 1.7.1^49^ and the ESS, ADSF and PSRF criteria. Once convergence was achieved, a 50% majority-rule consensus tree was built after discarding 25% of the sampled trees as burn-in.

Bayesian relaxed-clock methods were used that accounted for rate variation across lineages^53^. MCMC analyses implemented in BEAST 1.8.4^54^ were employed to approximate the posterior distribution of rates and divergences times and infer their credibility intervals. Estimation of divergence times relied on constraining clade ages through fossil calibrations. Three unambiguous fossils from perianth-bearing Piperales (Aristolochiaceae *sensu lato*), and one corresponding to the family Saururaceae were used. First, we relied on the fossil record of the monotypic family Lactoridaceae (*Lactoris fernandeziana*)^87, 92^, a shrub endemic to cloud forest of the Juan Fernández Islands archipelago of Chile. The oldest pollen fossil for the group is †*Lactoripollenites africanus*^96, 97^ from the Turonian/Campanian (72.1-89.8 Ma) of the Orange Basin in South Africa. This fossil confers a minimum age of 72.1 Ma for the stem node of *Lactoris fernandeziana*. Second, the oldest and only pollen record of the Aristolochiaceae was recently described from Late Cretaceous sediments of Siberia: †*Aristolochiacidites viluiensis*^98^ from the Timerdyakh Formation of the latest Campanian to earliest Maastrichtian (66-72.1 Ma) in the Vilui Basin (Russia). Because inaperturate pollen grains in combination with this unique exine configuration and fitting size can be observed in extant members of Aristolochiaceae, this fossil provides a minimum age of 66 Ma for the family. The third fossil belongs to the genus *Aristolochia* and described as †*Aristolochia austriaca*^99^ from the Pannonian (late Miocene) in the Hollabrunn-Mistelbach Formation (Austria). Based on a thorough morphological leaf comparison, this fossil is assigned to a species group including *Aristolochia baetica* and *Aristolochia rotunda*, which then confers a minimum age of 7.25 Ma for the clade. Finally, we used the fossil †*Saururus tuckerae*^100^ from the Princeton Chert of Princeton in British Columbia (Canada), which is part of the Princeton Group, Allenby Formation dated with stable isotopes to the middle Eocene^101^. This fossil has been phylogenetically placed as sister to extant *Saururus* species^101^, hence providing a minimum age of 44.3 Ma for the stem node of *Saururus*. Absolute ages of geological formations were taken from the latest update of the geological time scale.

We set the following settings and priors: a partitioned dataset (after the best-fitting PartitionFinder scheme) was analysed using the uncorrelated lognormal distribution clock model, with the mean set to a uniform prior between 0 and 1, and an exponential prior (lambda = 0.333) for the standard deviation. The branching process prior was set to a birth–death ^67^ process, using the following uniform priors: the birth–death mean growth rate ranged between 0 and 10 with a starting value at 0.1, and the birth–death relative death rate ranged between 0 and 1 (starting value = 0.5). We performed four independent BEAST analyses for 100 million generations, sampled every 10,000th, resulting in 10,000 samples in the posterior distribution of which the first 2500 samples were discarded as burn-in. All analyses were performed on the CIPRES Science Gateway computer cluster^51^, using BEAGLE^52^. Convergence and performance of each MCMC run were evaluated using Tracer 1.7.1^49^ and the ESS for each parameter. We combined the four runs using LogCombiner 1.8.4^54^. The MCC tree was reconstructed with median age and 95% CI. The BEAST files generated for this study are available in Figshare (see Data availability).

### Dual biogeographic history of Papilionidae and Aristolochiaceae

We estimated the ancestral area of origin and geographic range evolution for both clades using the ML approach of DEC model^69^ as implemented in the C++ version^102, 103^ that is available at: https://github.com/champost/DECX. To infer the biogeographic history of a clade, DEC requires a time-calibrated tree, the current distribution of each species for a set of geographic areas, and a time-stratified geographic model that is represented by connectivity matrices for specified time intervals spanning the entire evolutionary history of the group.

The geographic distribution for each species in Papilionidae^72–74^ and Aristolochiaceae was categorized as present or absent in each of the following areas: (1) West Nearctic [WN], (2) East Nearctic [EN], (3) Central America [CA], (4) South America [SA], (5) West Palearctic [WP], (6) East Palearctic [EP], (7) Madagascar [MD], (8) Indonesia and Wallacea [WA], (9) India [IN], (10) Africa [AF], and (11) Australasia [AU]. The resulting matrices of species distribution for the two groups are available in Figshare (see Data availability).

A time-stratified geographic model was built using connectivity matrices that take into account paleogeographic changes through time, with time slices indicating the possibility or not for a species to access a new area^103^. Based on palaeogeographical reconstructions^104–106^, we created a connectivity matrix for each geological epoch that represented a period bounded by major changes in tectonic and climatic conditions thought to have affected the distribution of organisms. The following geological epochs were selected: *(i)* 0 to 5.33 Ma (Pliocene to present), *(ii)* 5.33 to 23.03 Ma (Miocene), *(iii)* 23.03 to 33.9 Ma (Oligocene), *(iv)* 33.9 to 56 Ma (Eocene), and *(v)* 56 Ma to the origin of the clade (Palaeocene to Late Cretaceous). For each of these five time intervals, we specified constraints on area connectivity by coding 0 if any two areas are not connected or 1 if they are connected in a given time interval. We assumed a conservative dispersal matrix with equal dispersal rates between areas through time^107^.

### Impact of host-plant shifts on swallowtail diversification

We tested the effect of host-plant association on diversification by estimating speciation and extinction rates with five methods to cross-test hypotheses and corroborate results. Analyses were performed on 100 dated trees randomly sampled from the Bayesian dating analyses to take into account the uncertainty in age estimates. We used the following approaches: *(i)* ML-based trait-dependent diversification^108, 109^; *(ii)* ML-based time-dependent diversification^110^; *(iii)* Bayesian analysis of macroevolutionary mixture^111^; *(iv)* Bayesian branch-specific diversification rates^112^; and *(v)* Bayesian episodic birth-death model^113^. It is worth mentioning that each method differs at several points in their estimation of speciation and extinction rates. For instance, trait-dependent birth-death models estimate constant speciation and extinction rates ^109^, whereas time-dependent birth-death models estimate clade-specific speciation and extinction rates and their variation through time^110, 112^. Therefore, we expect some differences in the values of estimated diversification rates that are inherent to each approach. Our diversification analyses should be seen as complementary to the inferred diversification trend rather than corroborating the values and magnitude of speciation and extinction rates.

Firstly, we computed the probability of obtaining a clade as large as size *n*, given the crown age of origin, the overall net diversification rate of the family, and an extinction rate as a fraction of speciation rate following the approach in Condamine et al.^17^ relying on the method of moments^114^. We used the R-package *LASER* 2.3^115^ to estimate the net diversification rates of Papilionidae and six clades shifting to new host plants with the *bd.ms* function (providing crown age and total species diversity). Then we used the *crown.limits* function to estimate the mean expected clade size for each clade shifting to new host plants given clades’ crown age and overall net diversification rates, and we finally computed the probability to observe such clade size using the *crown.p* function. All rate estimates were calculated with three ε values (ε=0/0.5/0.9), knowing that the extinction rate in swallowtails is usually low^17^ (supported by the results of this study).

First, we relied on the state-dependent speciation and extinction (SSE) model, in which speciation and extinction rates are associated with phenotypic evolution of a trait along a phylogeny^108^. In particular, we used the Multiple State Speciation Extinction model (MuSSE^109^) implemented in the R-package *diversitree* 0.9–10^116^, which allows multiple character states to be studied. Larval host-plant data were taken from previous works^1, 12, 17, 72– 74, 117^. The following 10 host-plant character states and corresponding ratios of sampled species in the tree of all known species for each character (sampling fractions) were used: 1 = Aristolochiaceae (110/152), 2 = Annonaceae (69/138), 3 = Lauraceae (33/39), 4 = Apiaceae (9/10), 5 = Rutaceae (119/163), 6 = Crassulaceae (19/19), 7 = Papaveraceae (44/44), 8 = Fabaceae (1/1), 9 = Zygophyllaceae (2/2), and 10 = Magnoliaceae (2/2). Data at a lower taxonomic level than plant family were not used because of the large number of multiple associations exhibited by genera that could alter the phylogenetic signal. We assigned a single state to each species by selecting the foodplant with the maximum number of collections for each species. We did not employ multiple states per species, which represents a lesser problem because *(i)* few swallowtail species feed on multiple plant families, *(ii)* current shared-state models can only model two states, and *(iii)* the addition of multi-plant states to the MuSSE analysis would have greatly increased the number of parameters. We performed both ML and Bayesian MCMC analyses (10,000 steps) performed using an exponential (1/(2 × net diversification rate)) prior with starting parameter values obtained from the best-fitting ML model and resulting speciation, extinction and transition rates. After a burnin of 500 steps, we estimated posterior density distribution for speciation, extinction and transition rates. There have been concerns about the power of SSE models to infer diversification dynamics from a distribution of species traits^118–120^, hence other birth-death models were used to corroborate the results obtained with SSE models.

To provide an independent assessment of the relationship between diversification rates and host specificity, we used the ML approach of Morlon et al.^110^ implemented in the R-package *RPANDA* 1.3^121^. This is a birth–death method in which speciation and/or extinction rates may change continuously through time. This method has the advantage of not assuming constant extinction rate over time (unlike BAMM^111^), and allows clades to have declining diversity since extinction can exceed speciation, meaning that diversification rates can be negative^110^. For each clade that shifted to a new host family, we designed and fitted six diversification models: *(i)* a Yule model, where speciation is constant and extinction is null; *(ii)* a constant birth-death model, where speciation and extinction rates are constant; *(iii)* a variable speciation rate model without extinction; *(iv)* a variable speciation rate model with constant extinction; *(v)* a rate-constant speciation and variable extinction rate model; and *(vi)* a model in which both speciation and extinction rates vary. Models were compared by computing the ML estimate of each model and the resulting Akaike information criterion corrected by sample size (AICc) We then plotted rates through time with the best fit model for each clade, and the rates for the family as a whole for comparison purpose. We also performed models that allow diversification rates to vary among clades across the whole phylogeny. BAMM 2.5^111, 122^ was used to explore for differential diversification dynamic regimes among clades differing in their host-plant feeding. BAMM can automatically detect rate shifts and sample distinct evolutionary dynamics that explain the diversification dynamics of a clade without *a priori* hypotheses on how many and where these shifts might occur. Evolutionary dynamics can involve time-variable diversification rates; in BAMM, speciation is allowed to vary exponentially through time while extinction is maintained constant: subclades in a tree may diversify faster (or slower) than others. This Bayesian approach can be useful in detecting shifts of diversification potentially associated with key innovations^123^. BAMM analyses were run with four MCMC for 10 million generations, sampling every 10,000^th^ and with three different values (1, 5 and 10) of the compound Poisson prior (CPP) to ensure the posterior is independent of the prior^124^. We accounted for non-random incomplete taxon sampling using the implemented analytical correction; we set a sampling fraction per genus based on the known species diversity of each genus. Mixing and convergence among runs (ESS ≥ 200 after 15% burn-in) were assessed with the R-package *BAMMtools* 2.1^125^ to estimate *(i)* the mean global rates of diversification through time, *(ii)* the estimated number of rate shifts evaluating alternative diversification models comparing priors and posterior probabilities, and *(iii)* the clade-specific rates through time when a distinct macroevolutionary regime is identified.

BAMM has been criticized for incorrectly modelling rate-shifts on extinct lineages, that is, unobserved (extinct or unsampled) lineages inherit the ancestral diversification process and cannot experience subsequent diversification-rate shifts^124, 126^. To solve this, we used a novel Bayesian approach implemented in RevBayes 1.0.10^127^ that models rate shifts consistently on extinct lineages by using the SSE framework ^112, 124^. Although there is no information of rate shifts for unobserved/extinct lineages in a phylogeny including extant species only, these types of events must be accounted for in computing the likelihood. The number of rate categories is fixed in the analysis but RevBayes allows any number to be specified, thus allowing direct comparison of different macroevolutionary regimes.

Finally, we evaluated the impact of abrupt changes in diversification using the Bayesian episodic birth-death model of CoMET^113^ implemented in the R-package *TESS* 2.1^128^. These models allow detection of discrete changes in speciation and extinction rates concurrently affecting all lineages in a tree, and estimate changes in diversification rates at discrete points in time, but can also infer mass extinction events (sampling events in which the extant diversity is reduced by a fraction^129^). Speciation and extinction rates can change at those points but remain constant within time intervals. In addition, TESS uses independent CPPs to simultaneously detect mass extinction events and discrete changes in speciation and extinction rates, while TreePar estimates the magnitude and timing of speciation and extinction changes independently to the occurrence of mass extinctions (i.e. the three parameters cannot be estimated simultaneously due to parameter identifiability issues^129^). We performed two independent analyses allowing and disallowing mass extinction events. Bayes factor comparisons were used to assess model fit between models with varying number and time of changes in speciation/extinction rates and mass extinctions.

### Detecting genome-wide adaptations during host-plant shifts

We analysed genomic sequence data in swallowtails that have independently shifted to new ecological (biological) traits. Similar approaches have been conducted on mammals^130, 131^ and birds^132^, but have been rarely implemented on arthropod groups and, to our knowledge, this is the first time over such a long geological time scale. Here we estimated swallowtail molecular evolution with whole genome data and compared selection regimes on protein-coding genes along independent branches with or without host-plant shift and/or environmental shift.

For these analyses, we studied 45 whole genomes^33^ covering all 32 genera of the family Papilionidae: 41 of which were previously generated by our research group added to four genomes already available^30–32^. In summary, raw reads (Sequence Read Archive: SRR8954507-SRR8954549) were cleaned using Trimmomatic 0.33^133^, and assembled into contigs and scaffolds with SOAPdenovo-63mer 2.04^134^ to obtain whole genome assemblies (30x average read depth^33^). All coding DNA sequences (CDS) were retrieved from the high-quality annotated genome of *Papilio xuthus*^31^. To annotate the sequences of all our genomes, a BLAST search using all available CDS of *Papilio xuthus* was performed at the amino-acid level (using tblastn). For each species the recovered genes were aligned one by one with *Papilio xuthus* using TranslatorX^135^. This method performs alignment at the amino-acid level and preserves the open reading frame. All sites showing intraspecific variation were set to N, to conservatively avoid false informative sites. Any contamination was removed using CroCo 0.1^136^ and orthologous proteins were identified with OrthoFinder 2.2.0^137^. Finally, CDS alignments were strongly cleaned from misaligned sequences (gene by gene) using HMMCleaner 1.8^138^. A last cleaning step was performed using trimAl 1.2rev59^139^, which is designed to trim alignments for large-scale phylogenomic analyses. The resulting dataset comprised 6,621 genes in at least four sampled species (median of 32% of missing data), which was used to reconstruct a robust phylogenomic tree of Papilionidae^33^ (Supplementary Fig. 18).

We used this genomic dataset of 45 for all consisting on all genera in which the resulting genus-level swallowtail phylogenomic tree^33^ accurately represents the evolutionary associations with host plants as estimated using the ancestral-state analyses applied to the species-level phylogeny^17^ (Fig. 1, Supplementary Figs. 4, 5). We thus transferred the inference of ancestral host-plant shifts on the phylogenomic tree and selected the branches representing a host-plant shift and branches with a shift of climate preference (in general from tropical to temperate conditions; Supplementary Fig. 10). We also selected branches with no change as negative controls^34^. To test the impact of these different changes on the genomes, two datasets were created, *Dataset 1* and *2*. *Dataset 1* consists of 1,533 genes selected from the 6,621-gene dataset for each focal branch using three criteria: (1) the dataset is composed only of orthologous protein-coding genes (OrthoFinder 2.2^137^), (2) the species needed to accurately define the branch were available (i.e. crown node of the clade), and (3) for each branch, one species per tribe was available, and therefore include a different number of genes per branch. *Dataset 2* comprises 520 genes necessary to define all focal branches leading to less selected genes but the same genes for all branches. As a result, 14 branches are selected to measure the impact of a host-plant shift and 14 branches are selected as controls (Supplementary Fig. 18). Within these 28 branches, some branches represent environmental shifts (from tropical to temperate climate). The genomic dataset is available in Figshare (see Data availability).

We studied the ratio (ω) of nonsynonymous/synonymous substitution rate (dN/dS) to find genes under positive selection^37, 140^. The dN/dS ratio is traditionally used to estimate selective pressure from protein-coding sequences. If host-plant shifts have no effect on the selection of a given gene, we expect a dN/dS = 1 and the selective regime is considered neutral. However, if host-plant shifts result in positive selection on coding genes, the ratio increases such that dN/dS > 1. Finally, it is possible that host-plant shifts lead to purifying selection, thus reducing the number of non-synonymous substitutions and resulting in dN/dS < 1. Here we focused on the adaptation of Papilionidae to host plant shifts, i.e. outgroups are not studied. We tested if branches representing inferred host-plant shifts along the phylogeny of swallowtails have more genes with dN/dS > 1, representing adaptation, than branches representing host-plant conservatism. The *branch-site* models allow ω to vary both among sites in the protein and across branches on the tree and aim to detect positive selection affecting a few sites along particular lineages. The approach described by Zhang et al.^141^ is chosen to determine genome-wide selection regimes as performed with two maximum-likelihood models: (1) a null model assuming two site classes, one with dN/dS < 1 and one with dN/dS = 1; and (2) an alternative model adding a third site class with dN/dS > 1. The fit for including positive selection is tested using a likelihood ratio test comparing the null model with the alternative model with one degree of freedom^37, 142^. If the alternative model is better suited to host-shift branches, it is more likely the gene was under positive selection during the host-plant shifts. For each gene, dN/dS is estimated with both the null and alternative models using CodeML implemented in PAML 4^143^. To test the robustness of the estimations, we used a false discovery rate test to control false positives^144^. Finally, we reported the number of genes under positive selection on the total gene number for each focal branch. The number of genes under positive selection was compared between branches representing host-plant shifts, environmental shifts, both plant and environmental shifts or no shifts using the non-parametric Wilcoxon signed-rank test^145^.

### Sensitivity analyses

We performed several control analyses to ensure that the signal of more genes under positive selection in host-plant shifts branches is not artefactual. Specifically, we focused on missing data and GC content variation among genes known to bias dN/dS estimations. Missing data are prone to introducing misaligned regions that could create false positives in branch-site likelihood method for detecting positive selection^146–148^. Variations in GC content are known to impact the estimation of dN/dS mainly through the process of GC-biased gene conversion (gBGC^149–151^).

The number of missing data (‘N’ and ‘-’) sites and GC content at the third codon position (GC3) were computed using a home-made C++ program created with BIO++ library^152^. Mean GC content and missing data was calculated per gene and for each branch. For a given branch, mean GC3 and missing data were computed for the species of a clade for which the branch is the root. All statistics and graphical representations were performed using the R-packages *tidyverse*^153^ and *cowplot*^154^. We found that genes under positive selection (PS_genes_, *n*_Dataset1_ = 142, *n*_Dataset2_ = 407) have significantly more missing data and GC3 than genes not under positive selection (NS_genes_, *n*_Dataset1_ = 378, *n*_Dataset2_ = 1126; *P* = 0.001 / 0.02 for the two datasets, respectively, Mann-Whitney test; Supplementary Fig. 19). This result confirms that branch-site likelihood methods for detecting positive selection are sensitive to missing data, probably because of misaligned sites^146, 147^, and that GC content that may be influenced by gBGC^149^.

Missing data was, however, heterogeneously distributed among species, ranging from less than 1% in *Papilio xuthus* to 45% in *Hypermnestra helios* (Supplementary Fig. 20). The difference in missing data between branches with (*n* = 14, mean missing _Dataset1_ = 13.4%, mean missing_Dataset2_ = 14.1%) or without host-plant shifts (*n* = 14, mean missing_Dataset1_ = 12.8%, mean missing_Dataset2_ = 12.7%) is not significant (*P* = 0.83 / 1.00 for the two datasets, respectively, Mann-Whitney test; Supplementary Fig. 21). Additionally, there is no correlation between the number of genes under positive selection and the amount of missing data (*P* = 0.33 / 0.20 for the two datasets, respectively, Spearman’s correlation test; Supplementary Fig. 22). For GC3, we also found variation between species ranging from 37% in *Parnassius smintheus* to 44% in *Papilio antimachus* (Supplementary Fig. 23). Similarly to missing data, we found no significant difference between plant-shift and no plant-shift branches (*P* = 0.63 / 0.63 for the two datasets, Mann-Whitney test; Supplementary Fig. 24) and there is no correlation between the number of genes under positive selection and GC3 (*P* = 0.20 / 0.1362 for the two datasets, respectively, Spearman’s correlation test; Supplementary Fig. 25).

Despite the known fact that false positives can increase with the amount of missing data, our control analyses indicate that variations in missing data and GC content do not drive the signal that more genes are under positive selection in branches that have undergone a host-plant shift. Additionally to these controls, we checked by eyes all the gene alignments at the amino-acid level for genes under positive selection in branches with and without host-plant shifts using SeaView 4^155^. Misaligned regions, which could lead to biased dN/dS ratios^156^, were not significantly more detected for genes under positive selection in branches with host-plant shifts. In some cases we found ourselves in complicated situations to discriminate between false and true positive selected genes.

Overall, given the our alignment checks and sensitivity analyses, we do not see any reason for biased dN/dS ratios in genes along branches with or without host-plant shifts. False positive and false negative genes can be present in the two categories of branches but, in any cases, the general pattern observed is likely to remain conserved.

### Gene ontology

To annotate proteins of our alignment, we used the two different approaches implemented in PANTHER 14^157^ (available at: http://pantherdb.org/) and EggNOG 5.0^158, 159^ (available at: http://eggnog5.embl.de/#/app/home). We used the HMM Scoring tool to assign PANTHER family (library version 14.1^157^) to the protein of *Papilio xuthus* (assembly Pxut_1.0); similar results were obtained using another high-quality annotated genome (from *Heliconius melpomene*) as reference (assembly ASM31383v2). We performed the statistical overrepresentation test implemented on the PANTHER online website, relying on the GO categories in the PANTHER GO-Slim annotation dataset including Molecular function, Biological process, and Cellular component. Firstly, we tested if positively selected genes have over-or under-represented functional GO categories as compared to the whole set of genes (option “PANTHER Generic Mapping”). Secondly, we tested if positively selected genes involving a host-plant shift along the 14 branches have over- or under-represented functional categories. These statistical comparisons were performed with the Fisher’s exact test using the false discovery rate correction to control for false positives. Independently, we used the eggNOG-mapper v2^158^ (https://github.com/eggnogdb/eggnog-mapper) and the associated Lepidoptera database (LepNOG, including the genomes of *Bombyx mori*, *Danaus plexippus* and *Heliconius melpomene*^159^) to annotate the proteins of our dataset. EggNOG uses precomputed orthologous groups and phylogenies from the database to transfer functional information from fine-grained orthologs only. We used the diamond method as recommended^158^. Finally, we reported the known functions of proteins that were only positively selected when there was a host-plant shift in the phylogeny.

### Data availability

All data, including supermatrix datasets (for phylogenetic analyses), phylogenetic trees, host-plant preferences, species geographic distributions, gene alignments (for dN/dS analyses) and bioinformatic scripts, that are necessary for repeating the analyses described here have been made available through the Figshare digital data repository (https://figshare.com/s/1ce98308a3c012514857).

## Supplementary Figures

**Supplementary Figure 1.** Phylogenetic relationships of 408 swallowtail butterfly species (Papilionidae). Left phylogeny is inferred with the maximum-likelihood approach implemented with IQ-TREE, and right phylogeny is inferred with the Bayesian approach implemented with MrBayes. Both phylogenies show similar relationships except for the placement of the genus *Teinopalpus*, found as sister to Papilionini + Troidini with IQ-TREE and sister to *Meandrusa* (Papilionini) with MrBayes. Node support is indicated by ultrafast bootstrap and posterior probabilities on the maximum-likelihood and Bayesian phylogenies, respectively, with values of 95% and 0.95 considered as indicative of strong node support.

**Supplementary Figure 2.** Node support (ultrafast bootstrap) of the maximum-likelihood phylogeny. The histogram shows the distribution of node support for all Papilionidae, and indicates a high overall tree resolution with ∼80% of nodes having ultrafast bootstrap values ≥ 95%.

**Supplementary Figure 3.** Bayesian estimates of divergence times for swallowtail butterflies. The first inference was performed with exponential priors on fossil calibrations, while the second inference was carried out with uniform priors. The analysis based on exponential priors estimated a crown age for the family at 55.4 Ma (95% CI: 47.8-71.0 Ma), while the analysis based on uniform priors estimated the origin at 67.2 Ma (95% CI: 47.8-112 Ma).

**Supplementary Figure 4.** Estimation of ancestral host-plant preferences for the two molecular dated trees with the Dispersal-Extinction-Cladogenesis (DEC) model. The results show that the family Aristolochiaceae is recovered as the ancestral feeding habit of the Papilionidae. K = Cretaceous, Pl = Pliocene, P = Pleistocene.

**Supplementary Figure 5.** Estimation of ancestral host-plant preferences with the maximum-likelihood model of Markov 1-parameter (Mk) and the Bayesian approach of BayesTraits. The results are represented by pie charts indicating the relative probability for each state inferred at a given node. The results consistently show that (1) the family Aristolochiaceae is recovered as the ancestral feeding habit of the Papilionidae, and (2) the host-plant shifts are recovered at the same nodes, except at the root of Papilionini and at the root of *Iphiclides* + *Lamproptera* (due to the fact the the Mk model can include only 10 states).

**Supplementary Figure 6.** Estimation of ancestral host-plant preferences for the Aristolochiaceae feeders with the Dispersal-Extinction-Cladogenesis (DEC) model. The results show that the genus *Aristolochia* is the primary Aristolochiaceae host plant while being also recovered as the ancestral feeding habit of the Papilionidae.

**Supplementary Figure 7.** Phylogenetic relationships within the Aristolochiaceae (perianth-bearing Piperales) for 247 species. The phylogeny is inferred with the Bayesian approach of MrBayes. Node support is indicated by posterior probabilities, with values ≥ 0.95 considered as strong node support.

**Supplementary Figure 8.** Bayesian estimates of divergence times for Aristolochiaceae. The first inference was performed with exponential priors on fossil calibrations and 150 Ma as maximum age. The second inference was performed with exponential priors on fossil calibrations and 221 Ma as maximum age. The third inference was performed with uniform priors on fossil calibrations and 150 Ma as maximum age. The fourth inference was performed with uniform priors on fossil calibrations and 221 Ma as maximum age. The origin of the genus *Aristolochia* is estimated at 55.5 Ma (95% CI: 39.2-72.8 Ma) in the first analysis, at 58.8 Ma (95% CI: 42.5-76.2 Ma) in the second analysis, at 60.7 Ma (95% CI: 43.9-80.5 Ma) in the third analysis, and at 64.8 Ma (95% CI: 47.3-83.1 Ma) in the fourth analysis.

**Supplementary Figure 9.** Median node ages and 95% credibility intervals (CI) for the two dating analyses of Papilionidae and the four dating analyses of Aristolochiaceae. The 95% CI overlap substantially between the two groups regardless of the dating analysis. J = Jurassic, Pl = Pliocene, P = Pleistocene.

**Supplementary Figure 10.** Estimation of the historical biogeography for the two molecular dated trees of Papilionidae with the Dispersal-Extinction-Cladogenesis (DEC) model. For each tree, two DEC analyses were performed: one with time-stratified palaeogeographic constraints, and one without such constraints. The swallowtail butterflies originated in a northern region centred around the Bering land bridge. K = Cretaceous, Pl = Pliocene, P = Pleistocene.

**Supplementary Figure 11.** Estimation of the historical biogeography for the four molecular dated trees of Aristolochiaceae with the Dispersal-Extinction-Cladogenesis (DEC) model. For each tree, two DEC analyses were performed: one with time-stratified palaeogeographic constraints, and one without such constraints. The genus *Aristolochia* originated in a northern region centred around the Bering land bridge. J = Jurassic, K = Cretaceous, Pl = Pliocene, P = Pleistocene.

**Supplementary Figure 12.** Trait-dependent diversification of Papilionidae linked to their host plant. **a**, Bayesian inferences made with the full MuSSE model showed that speciation rates vary according to the host-plant trait. **b**, Boxplots showing the increase of diversification rates following host-plant shifts from the ancestral state (Aristolochiaceae). Only the species-poor swallowtail lineages feeding on Fabaceae, Zygophyllaceae and Magnoliaceae show decrease of diversification rates.

**Supplementary Figure 13.** Time-dependent diversification of Papilionidae after shifting to new host plants. Diversification is inferred with the RPANDA models, and the best-fit model is plotted showing rates through time for each clade. A model with increasing diversification over time best fits the Aristolochiaceae feeders. A model with a slowdown of diversification through time explained the diversification of Annonaceae feeders, Lauraceae feeders, and Papaveraceae feeders. A model with constant rates through time best fits the diversification of Apiaceae feeders, Crassulaceae feeders, and Rutaceae feeders. K = Cretaceous, Pl = Pliocene, P = Pleistocene.

**Supplementary Figure 14.** Bayesian analysis of clade-specific and time-dependent diversification of Papilionidae obtained with BAMM. a, Phylorate plot showing that global diversification rates increase through time in Papilionidae with no significant rate shifts detected by BAMM (the inset plot indicates the posterior probability for the estimated number of shifts). b, Rate-through-time plots for selected swallowtail lineages feeding on distinct host-plant families. The results also show an overall diversification increase through time for each group of swallowtails. P = Palaeocene, E = Eocene, O = Oligocene, M = Miocene.

**Supplementary Figure 15.** Bayesian analysis of branch-specific and time-dependent diversification of Papilionidae obtained with RevBayes. The median rates of diversification are plotted along each branch of the phylogeny, which shows a global increase of diversification rates through time in Papilionidae. Contrary to BAMM, this approach detected shifts in diversification rates in particular within the genera *Parnassius* and *Papilio* that have both shifted to new host-plant families. P = Palaeocene, E = Eocene, O = Oligocene, M = Miocene.

**Supplementary Figure 16.** Bayesian analysis of episodic diversification of Papilionidae obtained with CoMET. The four plots represent speciation, extinction, net diversification, and relative extinction rates through time for the whole family. The result indicates a global increase of diversification rates over time, notably starting ∼40 Ma. P = Palaeocene, E = Eocene, O = Oligocene, M = Miocene.

**Supplementary Figure 17.** Number of host plants consumed through time by Papilionidae. Using the estimation of ancestral host-plant preferences (Supplementary Fig. 4), we plotted the time at which a new host-plant family was colonised. This result shows that the swallowtail butterflies have a steady increase in the number of host families consumed over time. This ecological diversification can be paralleled with the global increase in diversification rates estimated by birth-death models (Supplementary Figs. 13-16). K = Cretaceous, Pl = Pliocene, P = Pleistocene.

**Supplementary Figure 18.** Genus-level phylogenomic tree of Papilionidae showing the 14 selected branches with host-plant shifts and the 14 selected branches without host-plant shifts (control branches). The selection of these branches is based on the estimation of ancestral state models using the species-level phylogenies and current host-plant preferences (Supplementary Figs. 4, 5).

**Supplementary Figure 19.** Violin plots of the percentage of missing data (“N” or “-”) and proportion of GC at third codon position (GC3) in alignment were positive selection have been detected (“Yes”) and positive selection have not been detected (“No”). Panels a and b are dataset 1 with 520 genes, and panels c and d are dataset 2 with 1533 genes.

**Supplementary Figure 20.** The percentage of missing data (“N” or “-”) per genes across species computed for dataset 1 and dataset 2.

**Supplementary Figure 21.** The percentage of missing data (“N” or “-”) per branch for the branches with (“Yes”, *n* = 14) and without (“No”, *n* = 14) host-plant shift. For a given branch, the percentage of missing data is the mean value of the species of a clade for which the branch is the root.

**Supplementary Figure 22.** Relationship between the percentage of missing data (“N” or “-”) and the number of positively selected genes per branch. For a given branch, the percentage of missing data is the mean value of the species of a clade for which the branch is the root.

**Supplementary Figure 23.** The percentage of GC at third codon position (GC3) per gene across species computed for dataset 1 and dataset 2.

**Supplementary Figure 24.** The percentage of GC at third codon position (GC3) per branch for the branches with (“Yes”, *n* = 14) and without (“No”, *n* = 14) host-plant shift. For a given branch, the percentage of GC3 is the mean value of the species of a clade for which the branch is the root.

**Supplementary Figure 25.** Relationship between the percentage of GC at third codon position (GC3) and the number of positively selected genes per branch. For a given branch, the percentage of GC3 is the mean value of the species of a clade for which the branch is the root.

**Supplementary Table 1.** Results from analyses of diversification rates performed with LASER. For clades shifting to new host plants, net diversification rates are estimated based on their crown age and extant species diversity using the method of moments. Net diversification rates for shifting clades are higher than the global rates of the family, suggesting that shifting to a new host plant confer higher rates of species diversification. Estimates of expected clade size based on the global diversification rates and crown age of shifting clades show that four clades diversified significantly faster than background diversification rates of non-shifting clades.

**Supplementary Table 2.** Information on orthogroups of dataset 2 (1,533 genes). The columns 2 to 6 indicate whether the genes are under positive selection and along which branch (column ‘Branch ID’ see Supplementary Fig. 18 for the annotated tree with branch numbers). The column ‘*Papilio xuthus* seq ID’ is the GenBank accession number for the corresponding sequences in *Papilio xuthus*. The column ‘PANTHER family:subfamily accession’ is family and subfamily accessions, and the column ‘PANTHER family name’ list the names for gene families based on PANTHER classification (see http://pantherdb.org/ for more information). Finally, ‘HMM e-value score’ is the Hidden Markov model e-value score, as reported by HMMER (Eddy 2011) performed through the online PANTHER scoring tool ftp://ftp.pantherdb.org/hmm_scoring/current_release. Following PANTHER recommendation, we have not considered e-values above 10^-11^ as significant.

