## Supplementary Figures for "Genome-wide macroevolutionary signatures of key innovations in butterflies colonizing new host plants"

**Supplementary figure 1.** Phylogenetic relationships of 408 swallowtail butterfly species (Papilionidae). Left phylogeny is inferred with the maximum-likelihood approach implemented with IQ-TREE, and right phylogeny is inferred with the Bayesian approach implemented with MrBayes. Both phylogenies show similar relationships except for the placement of the genus *Teinopalpus*, found as sister to Papilionini + Troidini with IQ-TREE and sister to *Meandrusa* (Papilionini) with MrBayes. Node support is indicated by ultrafast bootstrap and posterior probabilities on the maximum-likelihood and Bayesian phylogenies, respectively, with values of 95% and 0.95 considered as indicative of strong node support.

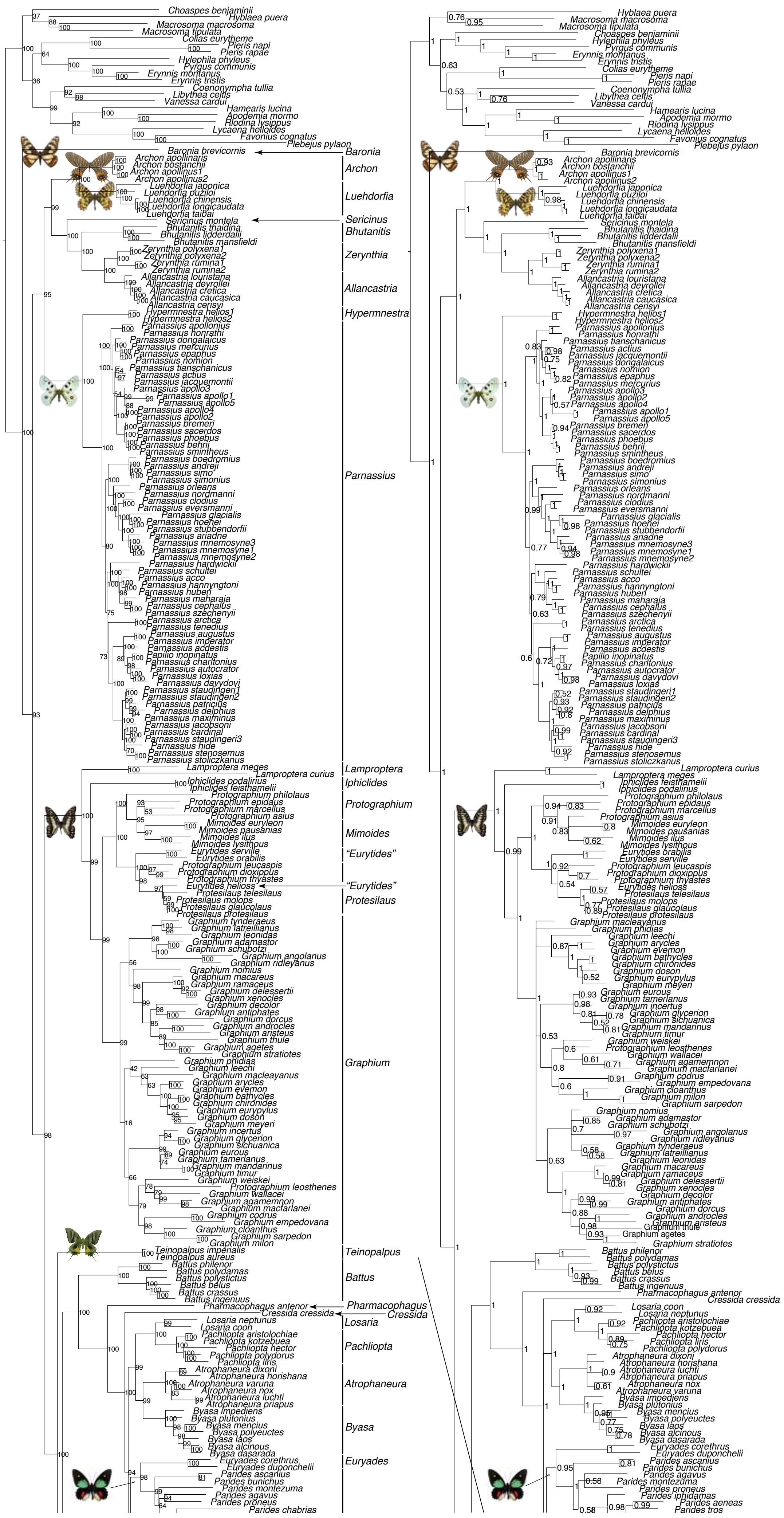

Baronia
Archon
Luehdorfia
Sericinus
Bhutanitis
Zerynthia
Allancastria
Hypermnestra
Parnassius
Lamproptera
Iphiclides
Protographium
Mimoides
"Eurytides"
"Eurytides"
Protesilaus
Graphium
Teinopalpus
Battus
Pharmacophagus
Cressida
Losaria
Pachliopta
Atrophaneura
Byasa
Euryades

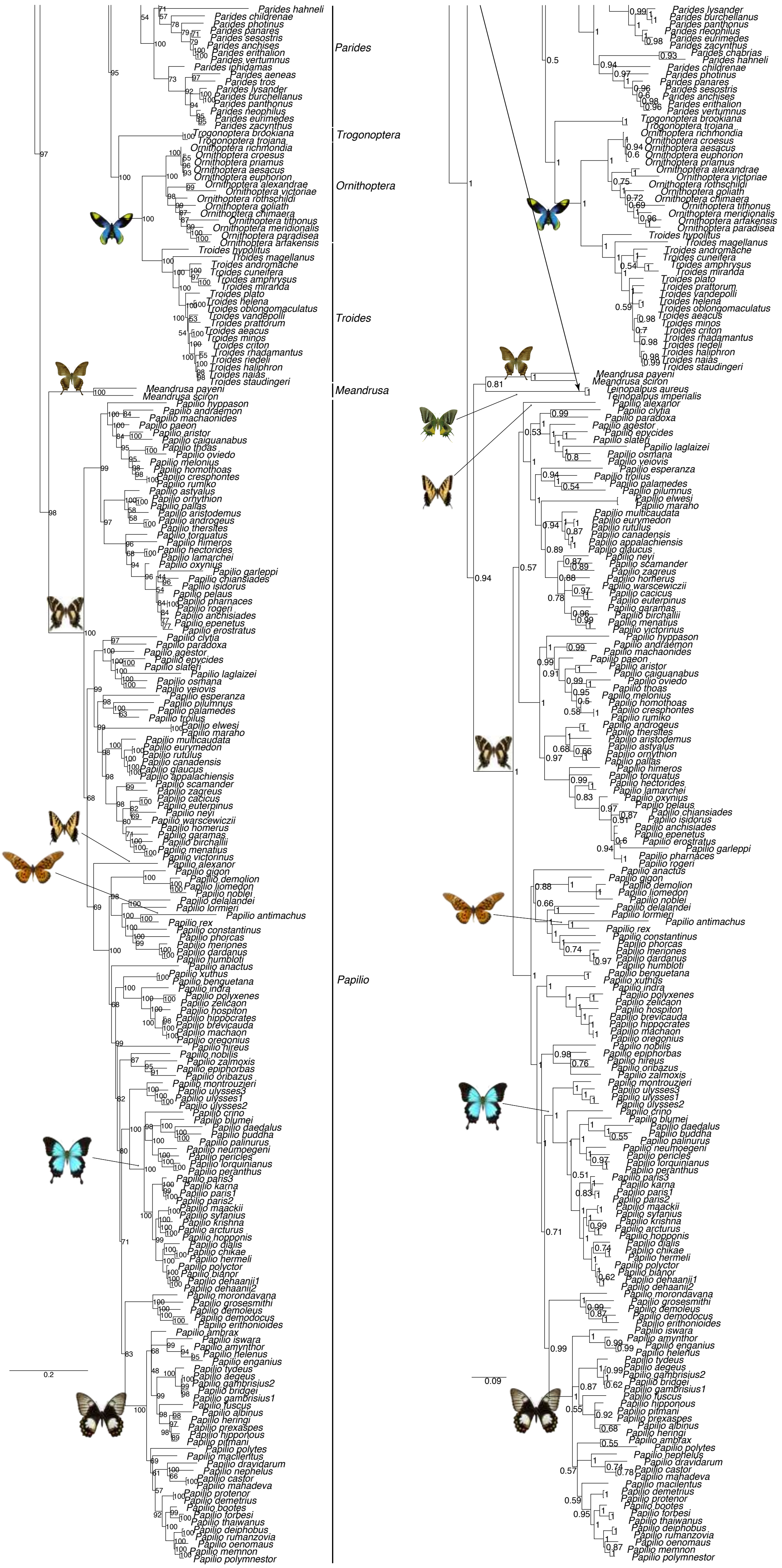
Parides
Trogonoptera
Ornithoptera
Troides
Meandrusa
Papilio
0.2
0.09

**Supplementary figure 2.** Node support (ultrafast bootstrap) of the maximum-likelihood phylogeny. The histogram shows the distribution of node support for all Papilionidae, and indicates a high overall tree resolution with ~80% of nodes having ultrafast bootstrap values ≥ 95%.

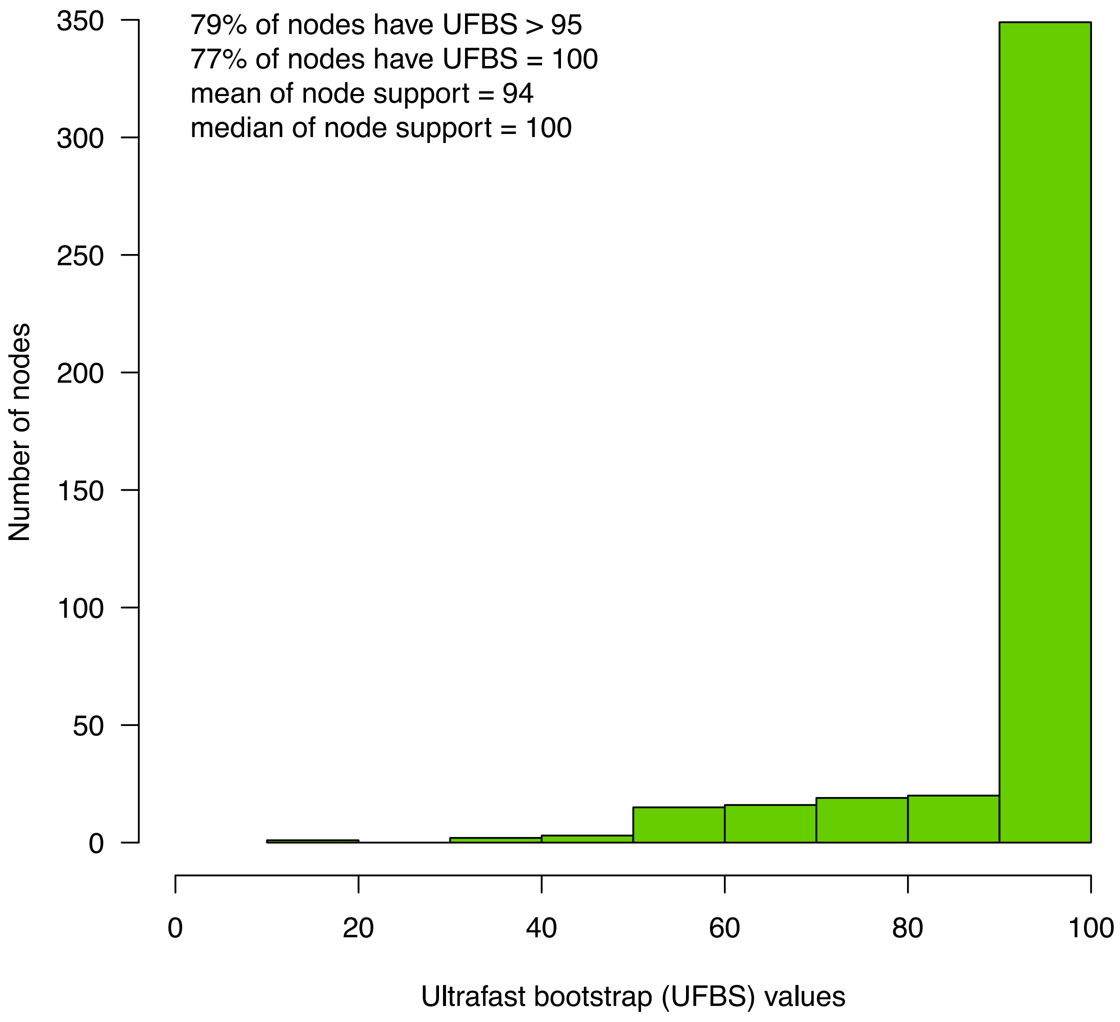

**Supplementary figure 3.** Bayesian estimates of divergence times for swallowtail butterflies. The first inference was performed with exponential priors on fossil calibrations, while the second inference was carried out with uniform priors. The analysis based on exponential priors estimated a crown age for the family at 55.4 Ma (95% CI: 47.8-71.0 Ma), while the analysis based on uniform priors estimated the origin at 67.2 Ma (95% CI: 47.8-112 Ma).

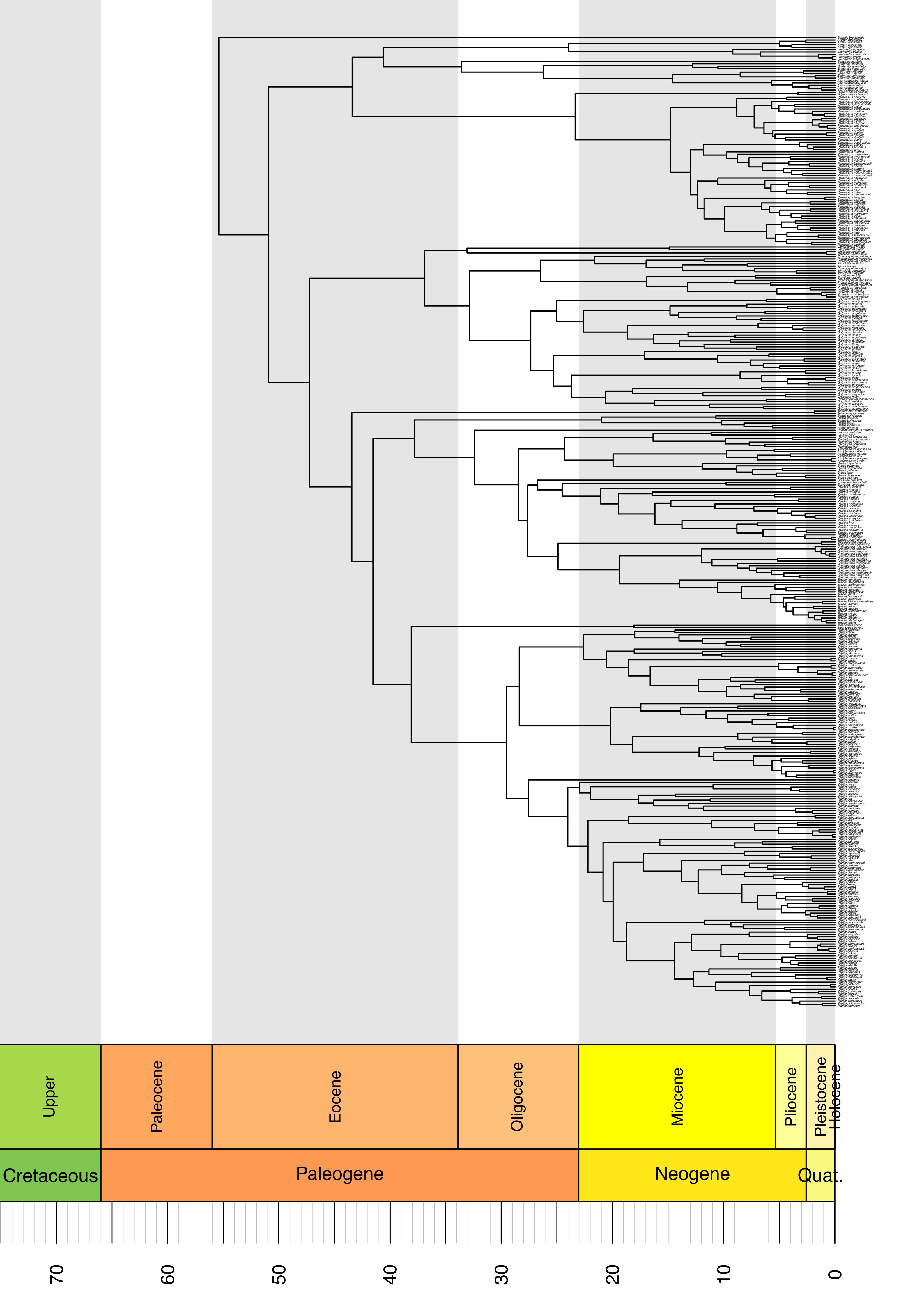
Upper
Paleocene
Eocene
Oligocene
Miocene
Pliocene
Pleistocene
Holocene
Cretaceous
Paleogene
Neogene
Quat.
70
60
50
40
30
20
10
0

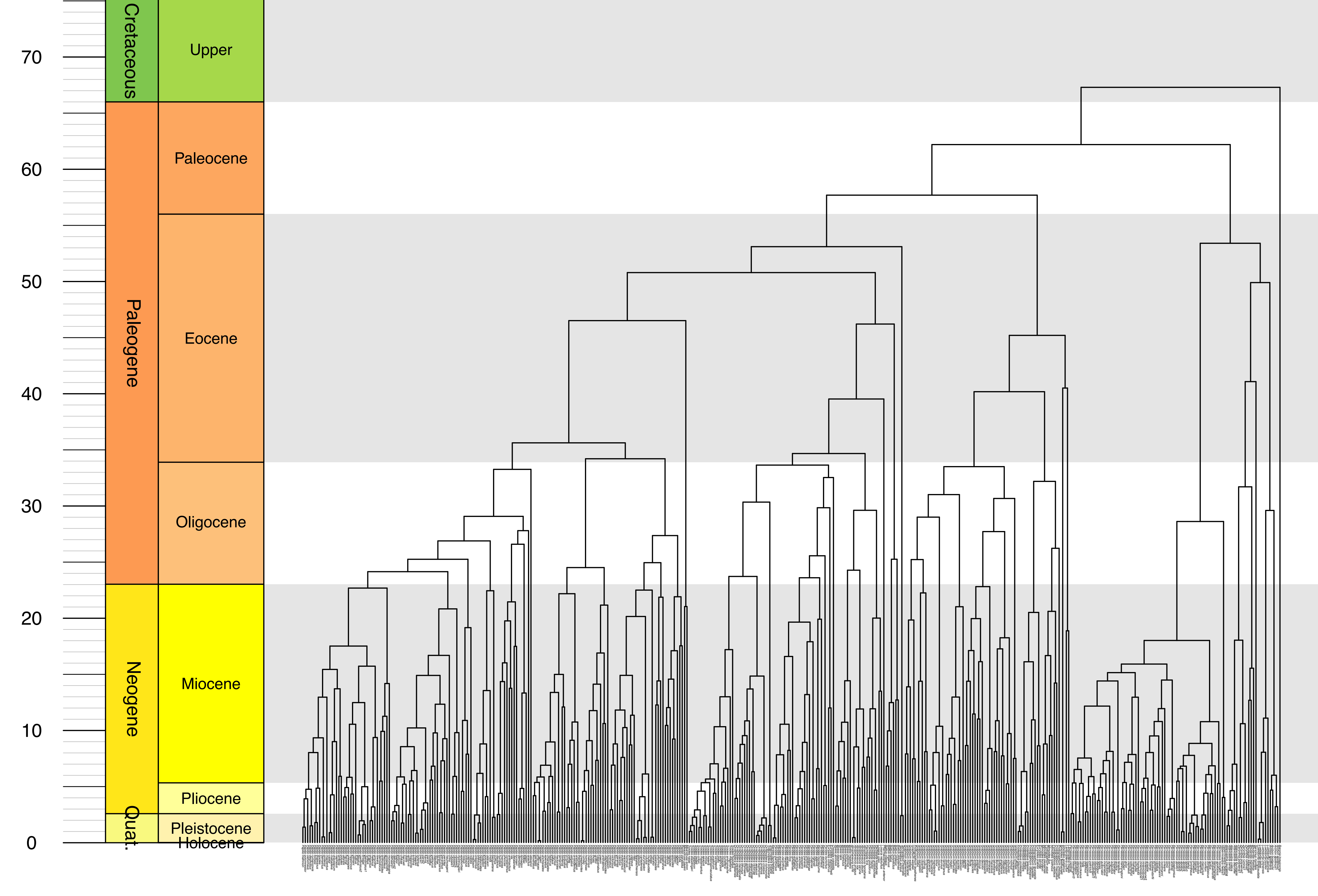

70
60
50
40
30
20
10
0
Cretaceous
Paleogene
Neogene
Quat.
Upper
Paleocene
Eocene
Oligocene
Miocene
Pliocene
Pleistocene
Holocene

**Supplementary figure 4.** Estimation of ancestral host-plant preferences for the two molecular dated trees with the Dispersal-Extinction-Cladogenesis (DEC) model. The results show that the family Aristolochiaceae is recovered as the ancestral feeding habit of the Papilionidae. K = Cretaceous, Pl = Pliocene, P = Pleistocene.

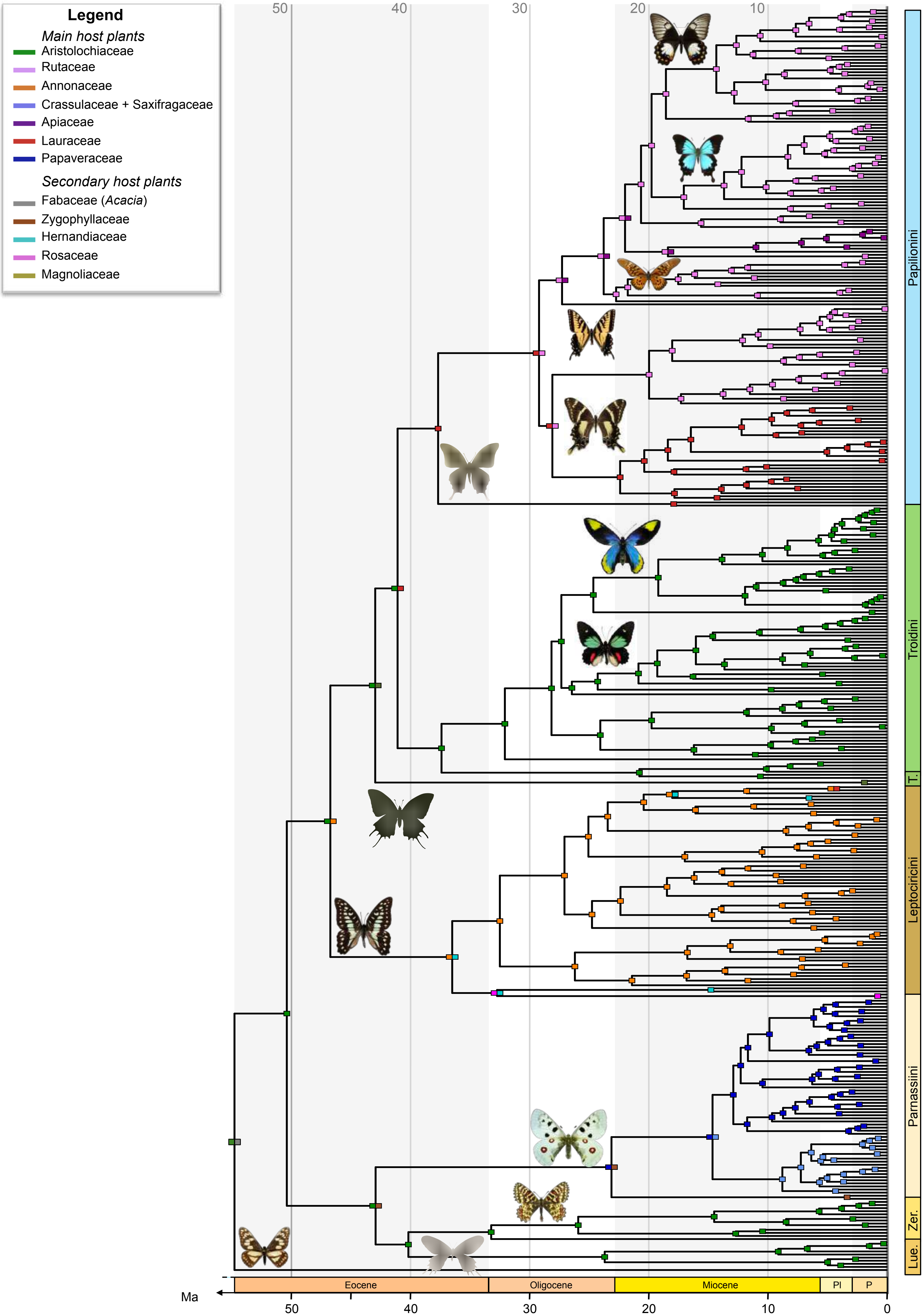

Legend
Main host plants
Aristolochiaceae
Rutaceae
Annonaceae
Crassulaceae + Saxifragaceae
Apiaceae
Lauraceae
Papaveraceae
Secondary host plants
Fabaceae (Acacia)
Zygophyllaceae
Hernandiaceae
Rosaceae
Magnoliaceae
50
40
30
20
10
Papilionini
Troidini
T.
Leptocircini
Parnassiini
Zer.
Lue.
Eocene
Oligocene
Miocene
Pl
P
Ma
50
40
30
20
10
0

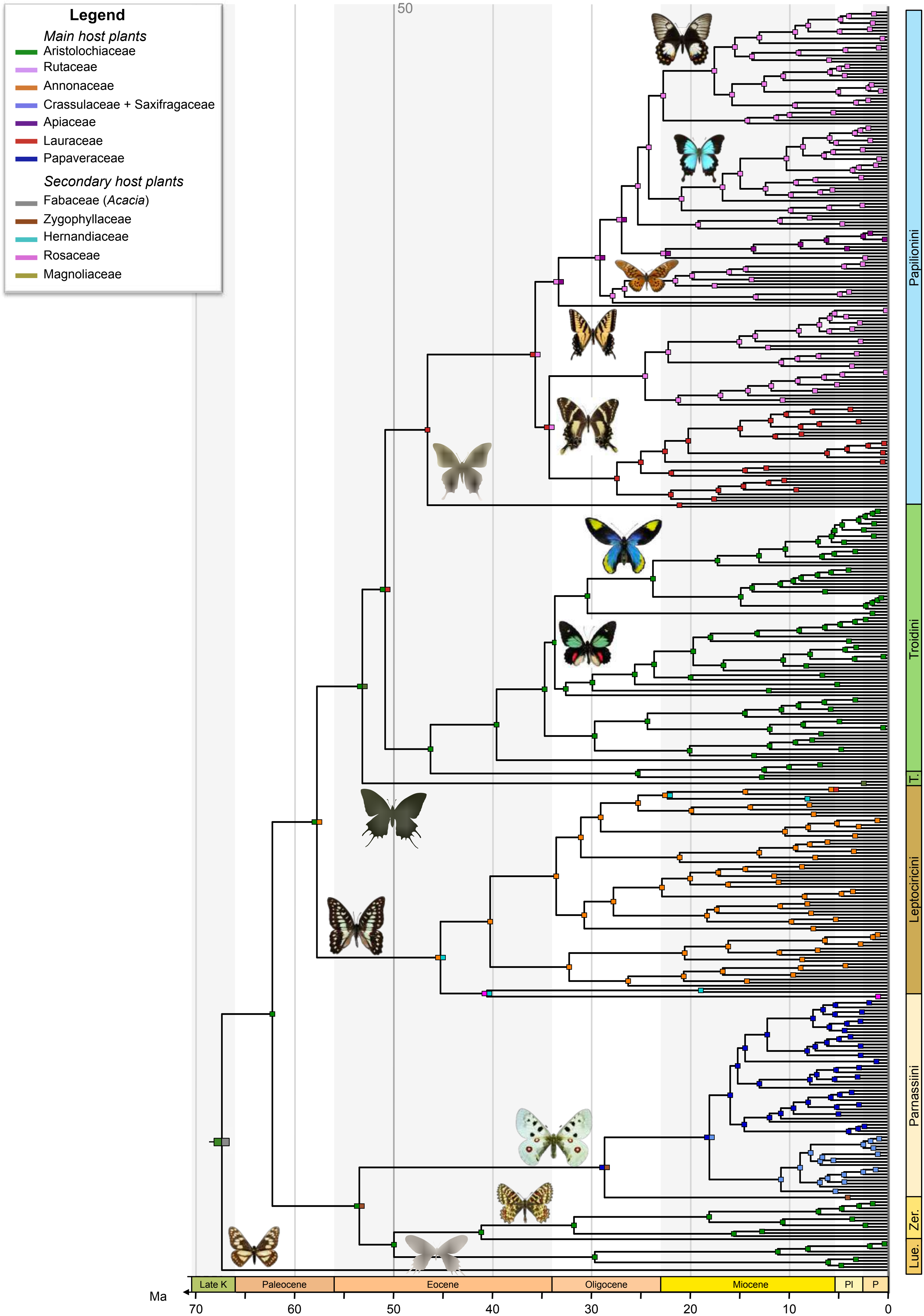
Legend
Main host plants
Aristolochiaceae
Rutaceae
Annonaceae
Crassulaceae + Saxifragaceae
Apiaceae
Lauraceae
Papaveraceae
Secondary host plants
Fabaceae (Acacia)
Zygophyllaceae
Hernandiaceae
Rosaceae
Magnoliaceae
Papilionini
Troidini
T.
Leptocircini
Parnassiini
Zer.
Lue.
Late K
Paleocene
Eocene
Oligocene
Miocene
Pl
P
Ma
70
60
50
40
30
20
10
0

**Supplementary figure 5.** Estimation of ancestral host-plant preferences with the maximum-likelihood model of Markov 1-parameter (Mk) and the Bayesian approach of BayesTraits. The results are represented by pie charts indicating the relative probability for each state inferred at a given node. The results consistently show that (1) the family Aristolochiaceae is recovered as the ancestral feeding habit of the Papilionidae, and (2) the host-plant shifts are recovered at the same nodes, except at the root of Papilionini and at the root of *Iphiclides* + *Lamproptera* (due to the fact the the Mk model can include only 10 states).

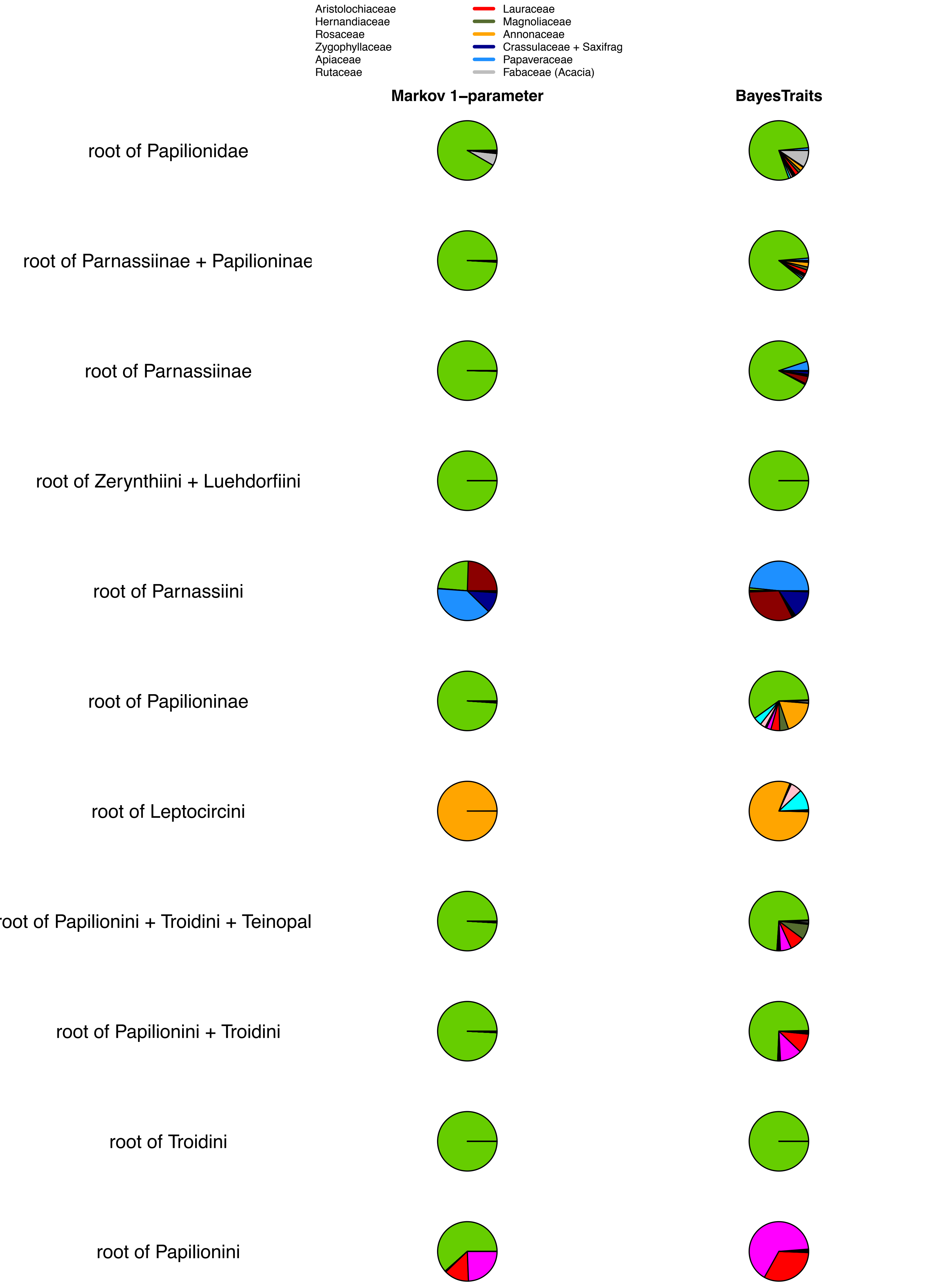
Aristolochiaceae
Hernandiaceae
Rosaceae
Zygophyllaceae
Apiaceae
Rutaceae
Lauraceae
Magnoliaceae
Annonaceae
Crassulaceae + Saxifrag
Papaveraceae
Fabaceae (Acacia)
Markov 1–parameter
BayesTraits
root of Papilionidae
root of Parnassiinae + Papilioninae
root of Parnassiinae
root of Zerynthiini + Luehdorfiini
root of Parnassiini
root of Papilioninae
root of Leptocircini
root of Papilionini + Troidini + Teinopal
root of Papilionini + Troidini
root of Troidini
root of Papilionini

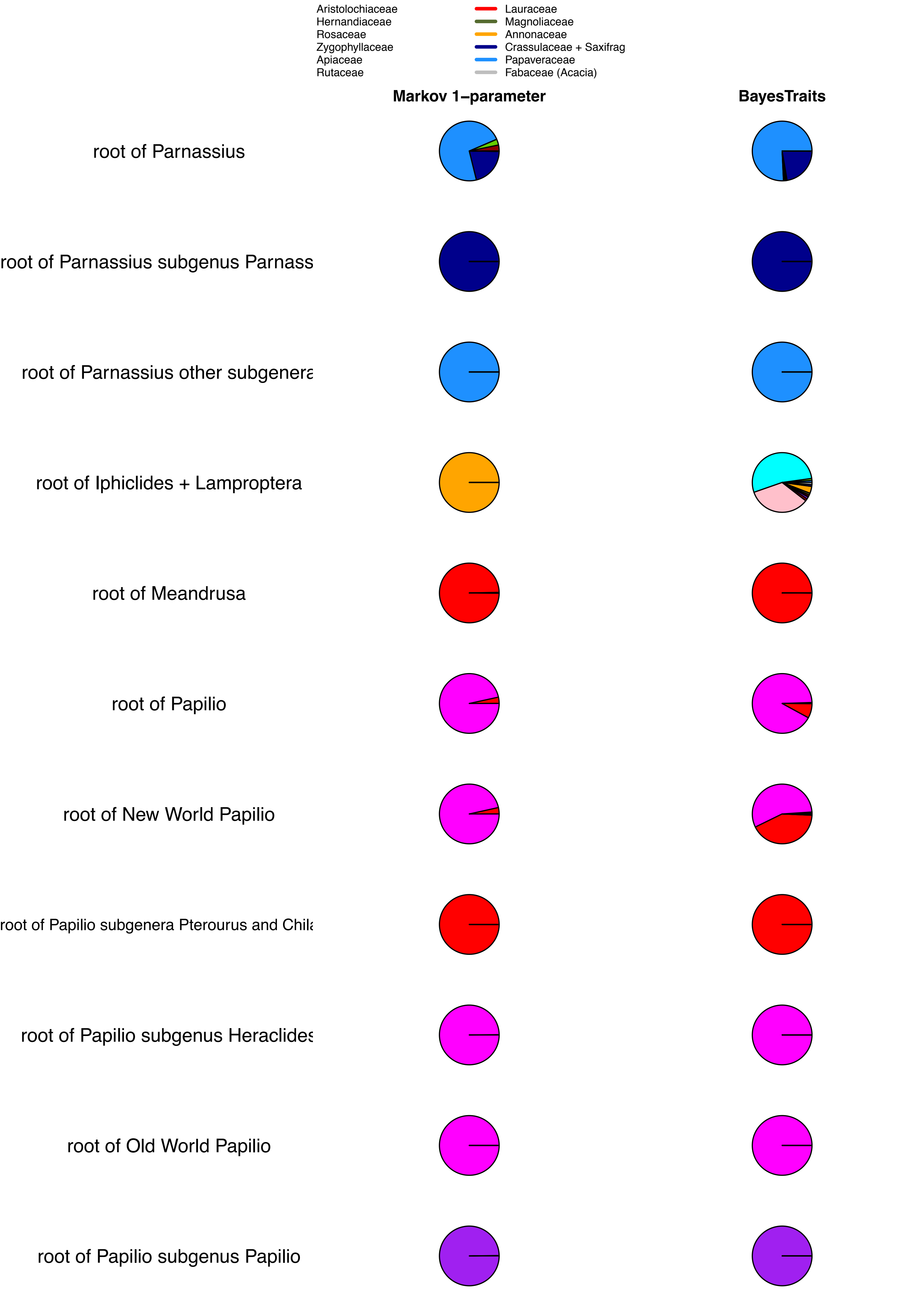
Aristolochiaceae
Hernandiaceae
Rosaceae
Zygophyllaceae
Apiaceae
Rutaceae
Lauraceae
Magnoliaceae
Annonaceae
Crassulaceae + Saxifrag
Papaveraceae
Fabaceae (Acacia)
Markov 1–parameter
BayesTraits
root of Parnassius
root of Parnassius subgenus Parnass
root of Parnassius other subgenera
root of Iphiclides + Lamproptera
root of Meandrusa
root of Papilio
root of New World Papilio
root of Papilio subgenera Pterourus and Chila
root of Papilio subgenus Heraclides
root of Old World Papilio
root of Papilio subgenus Papilio

**Supplementary figure 6.** Estimation of ancestral host-plant preferences for the Aristolochiaceae feeders with the Dispersal-Extinction-Cladogenesis (DEC) model. The results show that the genus *Aristolochia* is the primary Aristolochiaceae host plant while being also recovered as the ancestral feeding habit of the Papilionidae.

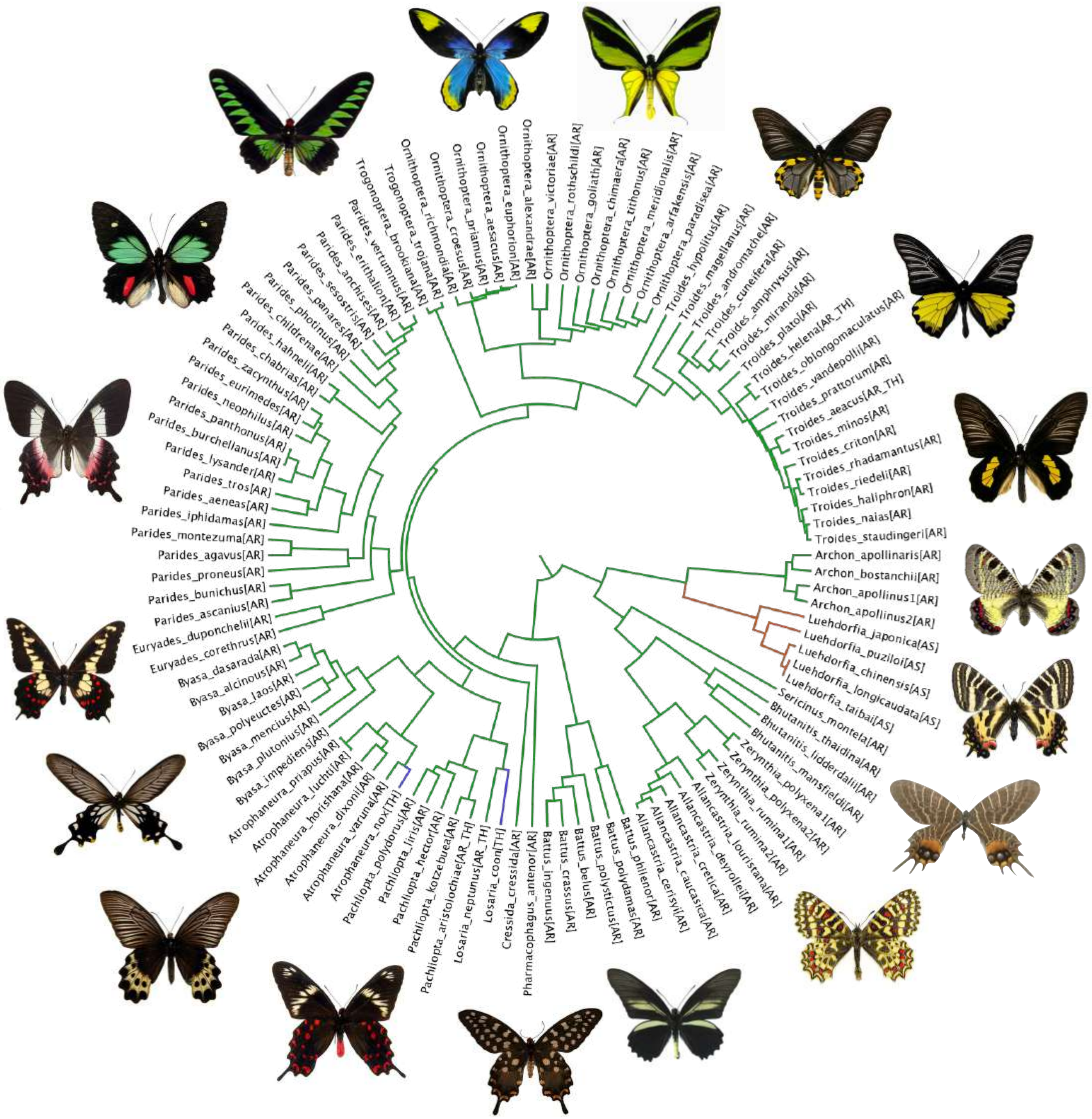


**Legend**

| | | |
|---|---|---|
| (green) | *Aristolochia* | [AR] |
| (orange) | *Asarum* | [AS] |
| (blue) | *Thottea* [TH] | |

**Supplementary figure 7.** Phylogenetic relationships within the Aristolochiaceae (perianth-bearing Piperales) for 247 species. The phylogeny is inferred with the Bayesian approach of MrBayes. Node support is indicated by posterior probabilities, with values $\geq$ 0.95 considered as strong node support.

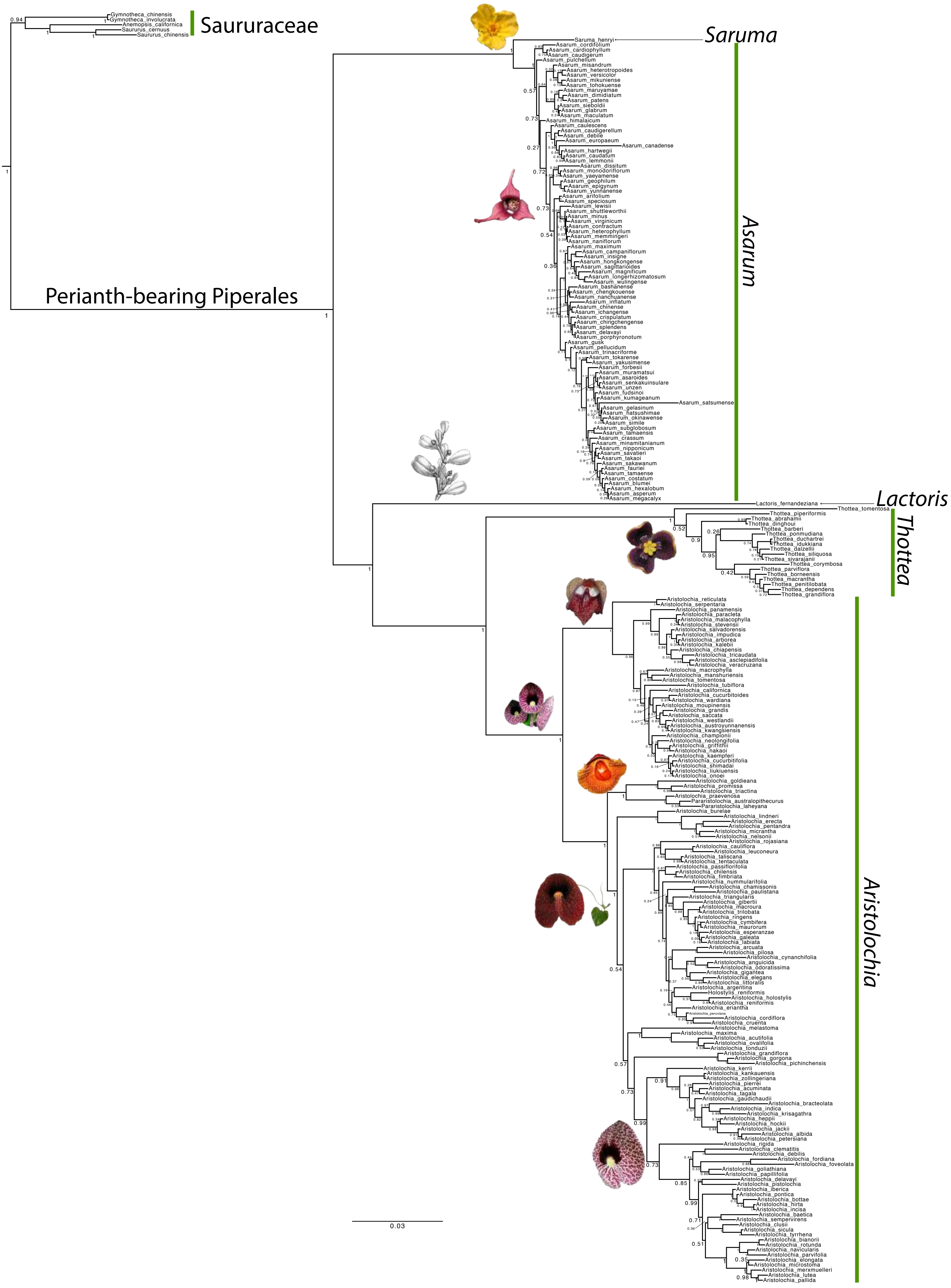

Saururaceae
Gymnotheca_chinensis
Gymnotheca_involucrata
Anemopsis_californica
Saururus_cernuus
Saururus_chinensis
Perianth-bearing Piperales
Saruma
Saruma_henryi
Asarum
Asarum_cordifolium
Asarum_cardiophyllum
Asarum_caudigerum
Asarum_pulchellum
Asarum_misandrum
Asarum_heterotropoides
Asarum_versicolor
Asarum_mikuniense
Asarum_tohokuense
Asarum_maruyamae
Asarum_dimidiatum
Asarum_patens
Asarum_sieboldii
Asarum_glabrum
Asarum_maculatum
Asarum_himalaicum
Asarum_caulescens
Asarum_caudigerellum
Asarum_debile
Asarum_europaeum
Asarum_canadense
Asarum_hartwegii
Asarum_caudatum
Asarum_lemmonii
Asarum_dissitum
Asarum_monodoriflorum
Asarum_yaeyamense
Asarum_geophilum
Asarum_epigynum
Asarum_yunnanense
Asarum_arifolium
Asarum_speciosum
Asarum_lewisii
Asarum_shuttleworthii
Asarum_minus
Asarum_virginicum
Asarum_contractum
Asarum_heterophyllum
Asarum_memmingeri
Asarum_naniflorum
Asarum_maximum
Asarum_campaniflorum
Asarum_insigne
Asarum_hongkongense
Asarum_sagittarioides
Asarum_magnificum
Asarum_longerhizomatosum
Asarum_wulingense
Asarum_bashanense
Asarum_chengkouense
Asarum_nanchuanense
Asarum_inflatum
Asarum_chinense
Asarum_ichangense
Asarum_crispulatum
Asarum_chingchengense
Asarum_splendens
Asarum_delavayi
Asarum_porphyronotum
Asarum_gusk
Asarum_pellucidum
Asarum_trinacriforme
Asarum_tokarense
Asarum_yakusimense
Asarum_forbesii
Asarum_muramatsui
Asarum_asaroides
Asarum_senkakuinsulare
Asarum_unzen
Asarum_fudsinoi
Asarum_kumageanum
Asarum_satsumense
Asarum_gelasinum
Asarum_hatsushimae
Asarum_okinawense
Asarum_simile
Asarum_subglobosum
Asarum_tamaensis
Asarum_crassum
Asarum_minamitanianum
Asarum_nipponicum
Asarum_savatieri
Asarum_takaoi
Asarum_sakawanum
Asarum_fauriei
Asarum_tamaense
Asarum_costatum
Asarum_blumei
Asarum_hexalobum
Asarum_asperum
Asarum_megacalyx
Lactoris
Lactoris_fernandeziana
Thottea
Thottea_tomentosa
Thottea_piperiformis
Thottea_abrahamii
Thottea_dinghoui
Thottea_barberi
Thottea_ponmudiana
Thottea_duchartrei
Thottea_idukkiana
Thottea_dalzellii
Thottea_siliquosa
Thottea_sivarajanii
Thottea_corymbosa
Thottea_parviflora
Thottea_borneensis
Thottea_macrantha
Thottea_penitilobata
Thottea_dependens
Thottea_grandiflora
Aristolochia
Aristolochia_reticulata
Aristolochia_serpentaria
Aristolochia_panamensis
Aristolochia_paracleta
Aristolochia_malacophylla
Aristolochia_stevensii
Aristolochia_salvadorensis
Aristolochia_impudica
Aristolochia_arborea
Aristolochia_kalebii
Aristolochia_chiapensis
Aristolochia_tricaudata
Aristolochia_asclepiadifolia
Aristolochia_veracruzana
Aristolochia_macrophylla
Aristolochia_manshuriensis
Aristolochia_tomentosa
Aristolochia_tubiflora
Aristolochia_californica
Aristolochia_cucurbitoides
Aristolochia_wardiana
Aristolochia_moupinensis
Aristolochia_grandis
Aristolochia_saccata
Aristolochia_westlandii
Aristolochia_austroyunnanensis
Aristolochia_kwangsiensis
Aristolochia_championii
Aristolochia_neolongifolia
Aristolochia_griffithii
Aristolochia_nakaoi
Aristolochia_kaempferi
Aristolochia_cucurbitifolia
Aristolochia_shimadai
Aristolochia_liukiuensis
Aristolochia_onoei
Aristolochia_goldieana
Aristolochia_promissa
Aristolochia_triactina
Aristolochia_praevenosa
Pararistolochia_australopithecurus
Pararistolochia_laheyana
Aristolochia_burelae
Aristolochia_lindneri
Aristolochia_erecta
Aristolochia_pentandra
Aristolochia_micrantha
Aristolochia_nelsonii
Aristolochia_rojasiana
Aristolochia_cauliflora
Aristolochia_leuconeura
Aristolochia_taliscana
Aristolochia_tentaculata
Aristolochia_passiflorifolia
Aristolochia_chilensis
Aristolochia_fimbriata
Aristolochia_nummularifolia
Aristolochia_chamissonis
Aristolochia_paulistana
Aristolochia_triangularis
Aristolochia_gibertii
Aristolochia_macroura
Aristolochia_trilobata
Aristolochia_ringens
Aristolochia_cymbifera
Aristolochia_maurorum
Aristolochia_esperanzae
Aristolochia_galeata
Aristolochia_labiata
Aristolochia_arcuata
Aristolochia_pilosa
Aristolochia_cynanchifolia
Aristolochia_anguicida
Aristolochia_odoratissima
Aristolochia_gigantea
Aristolochia_elegans
Aristolochia_littoralis
Aristolochia_argentina
Holostylis_reniformis
Aristolochia_holostylis
Aristolochia_reniformis
Aristolochia_eriantha
Aristolochia_peruviana
Aristolochia_cordiflora
Aristolochia_cruenta
Aristolochia_melastoma
Aristolochia_maxima
Aristolochia_acutifolia
Aristolochia_ovalifolia
Aristolochia_tonduzii
Aristolochia_grandiflora
Aristolochia_gorgona
Aristolochia_pichinchensis
Aristolochia_kerrii
Aristolochia_kankauensis
Aristolochia_zollingeriana
Aristolochia_pierrei
Aristolochia_acuminata
Aristolochia_tagala
Aristolochia_gaudichaudii
Aristolochia_bracteolata
Aristolochia_indica
Aristolochia_krisagathra
Aristolochia_heppii
Aristolochia_hockii
Aristolochia_jackii
Aristolochia_albida
Aristolochia_petersiana
Aristolochia_rigida
Aristolochia_clematitis
Aristolochia_debilis
Aristolochia_fordiana
Aristolochia_foveolata
Aristolochia_goliathiana
Aristolochia_papillifolia
Aristolochia_delavayi
Aristolochia_pistolochia
Aristolochia_iberica
Aristolochia_pontica
Aristolochia_bottae
Aristolochia_hirta
Aristolochia_incisa
Aristolochia_baetica
Aristolochia_sempervirens
Aristolochia_clusii
Aristolochia_sicula
Aristolochia_tyrrhena
Aristolochia_bianorii
Aristolochia_rotunda
Aristolochia_navicularis
Aristolochia_parvifolia
Aristolochia_elongata
Aristolochia_microstoma
Aristolochia_merxmuelleri
Aristolochia_lutea
Aristolochia_pallida
0.03

**Supplementary figure 8.** Bayesian estimates of divergence times for Aristolochiaceae. The first inference was performed with exponential priors on fossil calibrations and 150 Ma as maximum age. The second inference was performed with exponential priors on fossil calibrations and 221 Ma as maximum age. The third inference was performed with uniform priors on fossil calibrations and 150 Ma as maximum age. The fourth inference was performed with uniform priors on fossil calibrations and 221 Ma as maximum age. The origin of the genus *Aristolochia* is estimated at 55.5 Ma (95% CI: 39.2-72.8 Ma) in the first analysis, at 58.8 Ma (95% CI: 42.5-76.2 Ma) in the second analysis, at 60.7 Ma (95% CI: 43.9-80.5 Ma) in the third analysis, and at 64.8 Ma (95% CI: 47.3-83.1 Ma) in the fourth analysis.

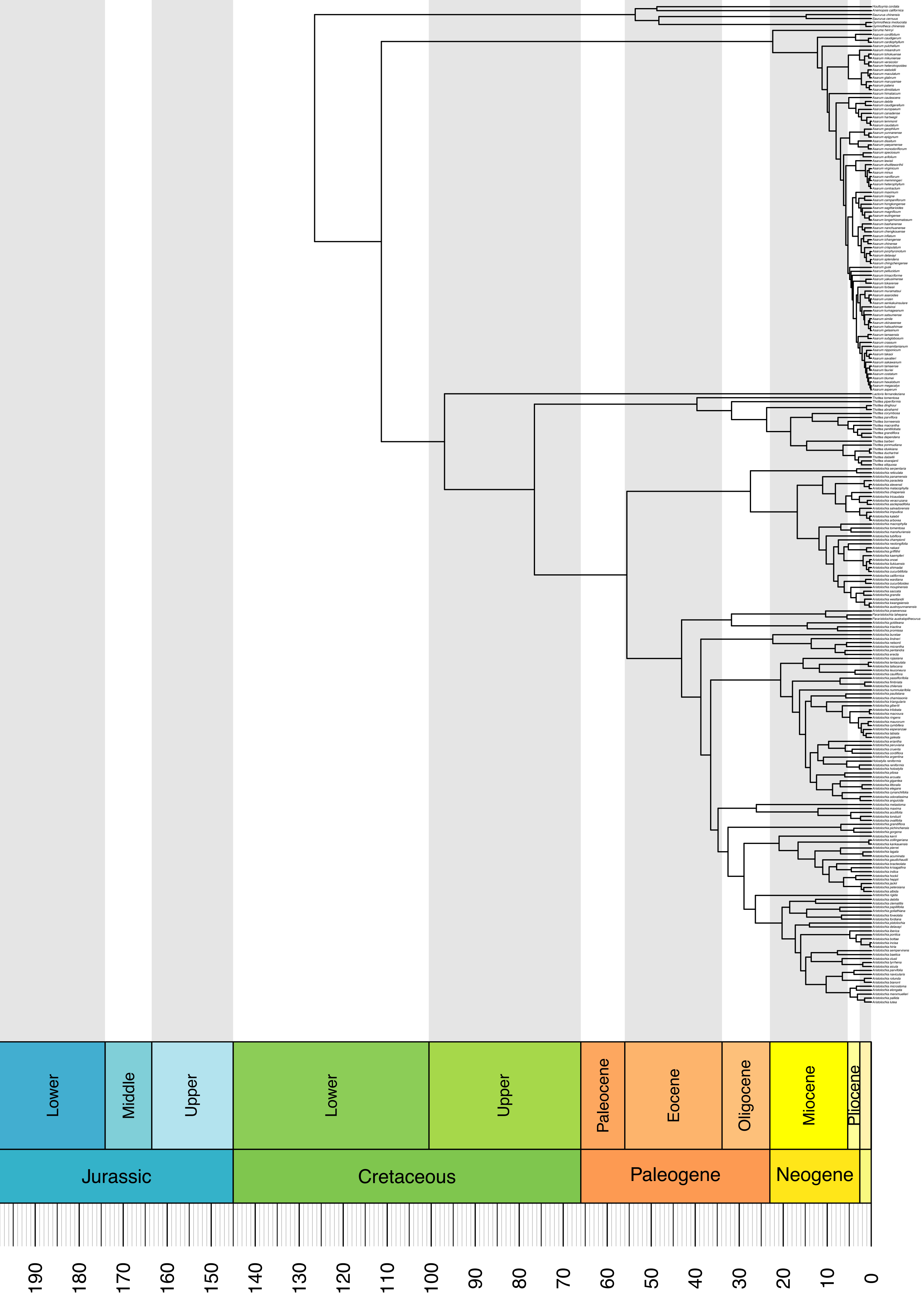
Lower
Middle
Upper
Lower
Upper
Paleocene
Eocene
Oligocene
Miocene
Pliocene
Jurassic
Cretaceous
Paleogene
Neogene
190
180
170
160
150
140
130
120
110
100
90
80
70
60
50
40
30
20
10
0

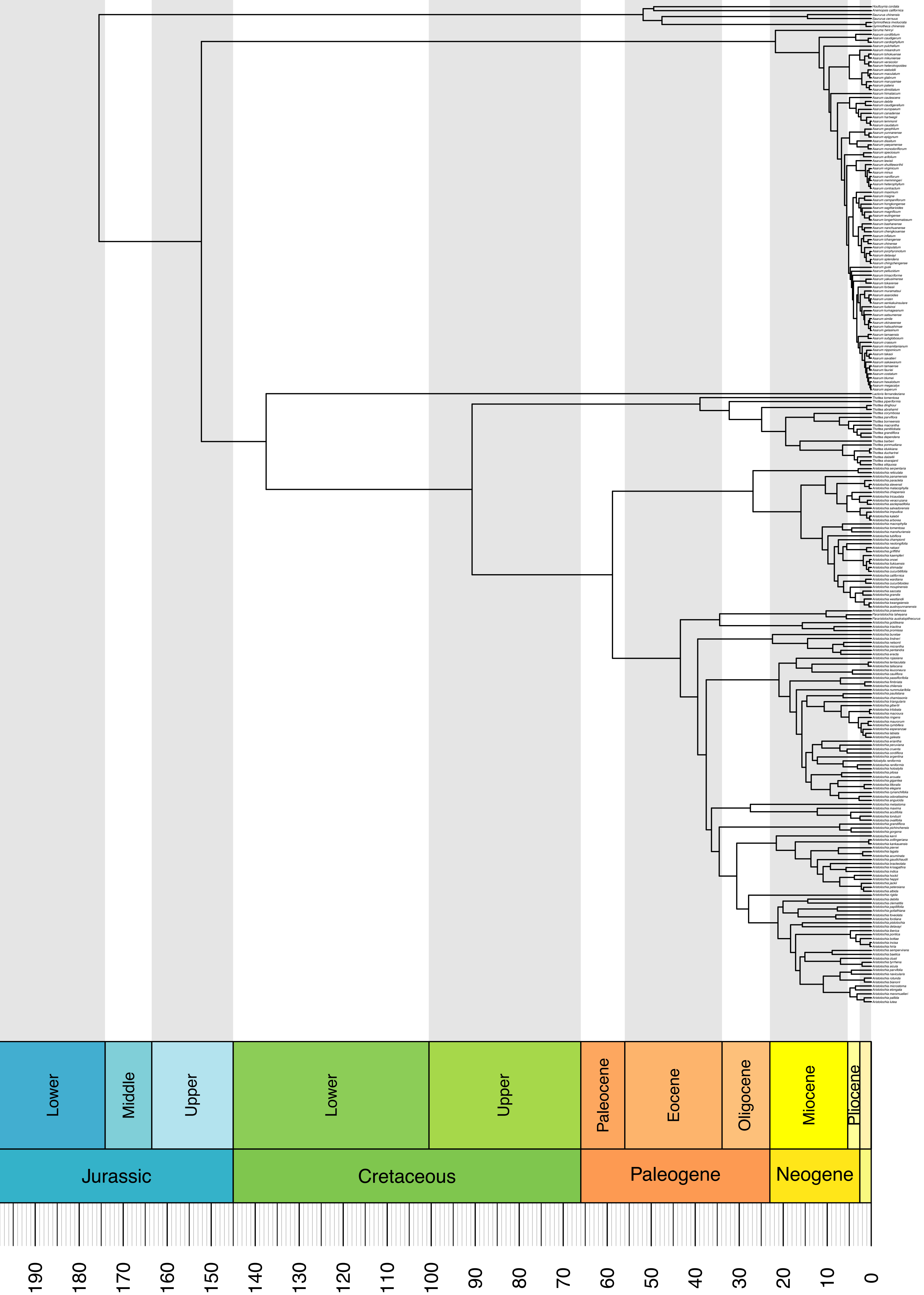

Lower
Middle
Upper
Lower
Upper
Paleocene
Eocene
Oligocene
Miocene
Pliocene
Jurassic
Cretaceous
Paleogene
Neogene
190
180
170
160
150
140
130
120
110
100
90
80
70
60
50
40
30
20
10
0

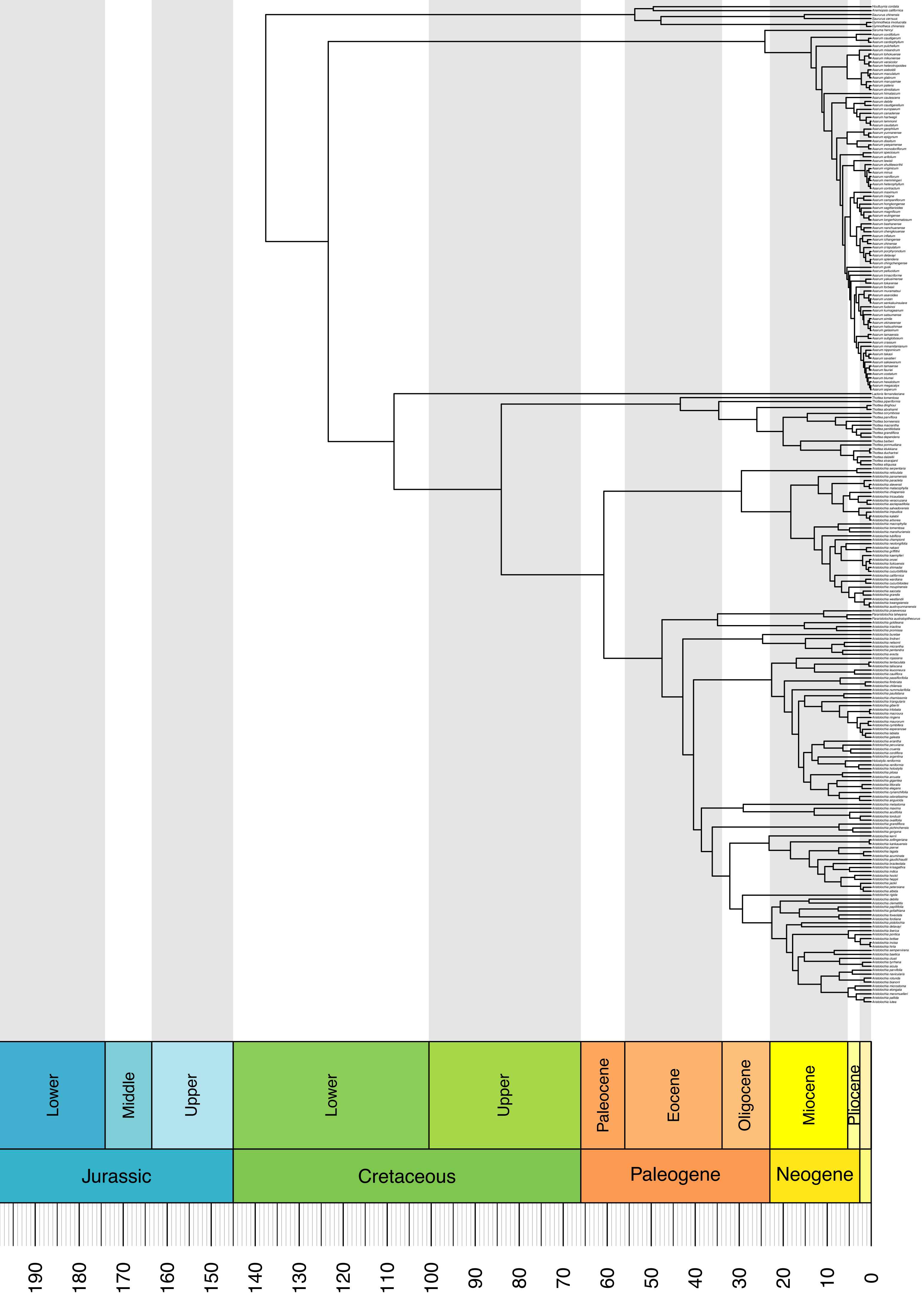
Lower
Middle
Upper
Lower
Upper
Paleocene
Eocene
Oligocene
Miocene
Pliocene
Jurassic
Cretaceous
Paleogene
Neogene
190
180
170
160
150
140
130
120
110
100
90
80
70
60
50
40
30
20
10
0

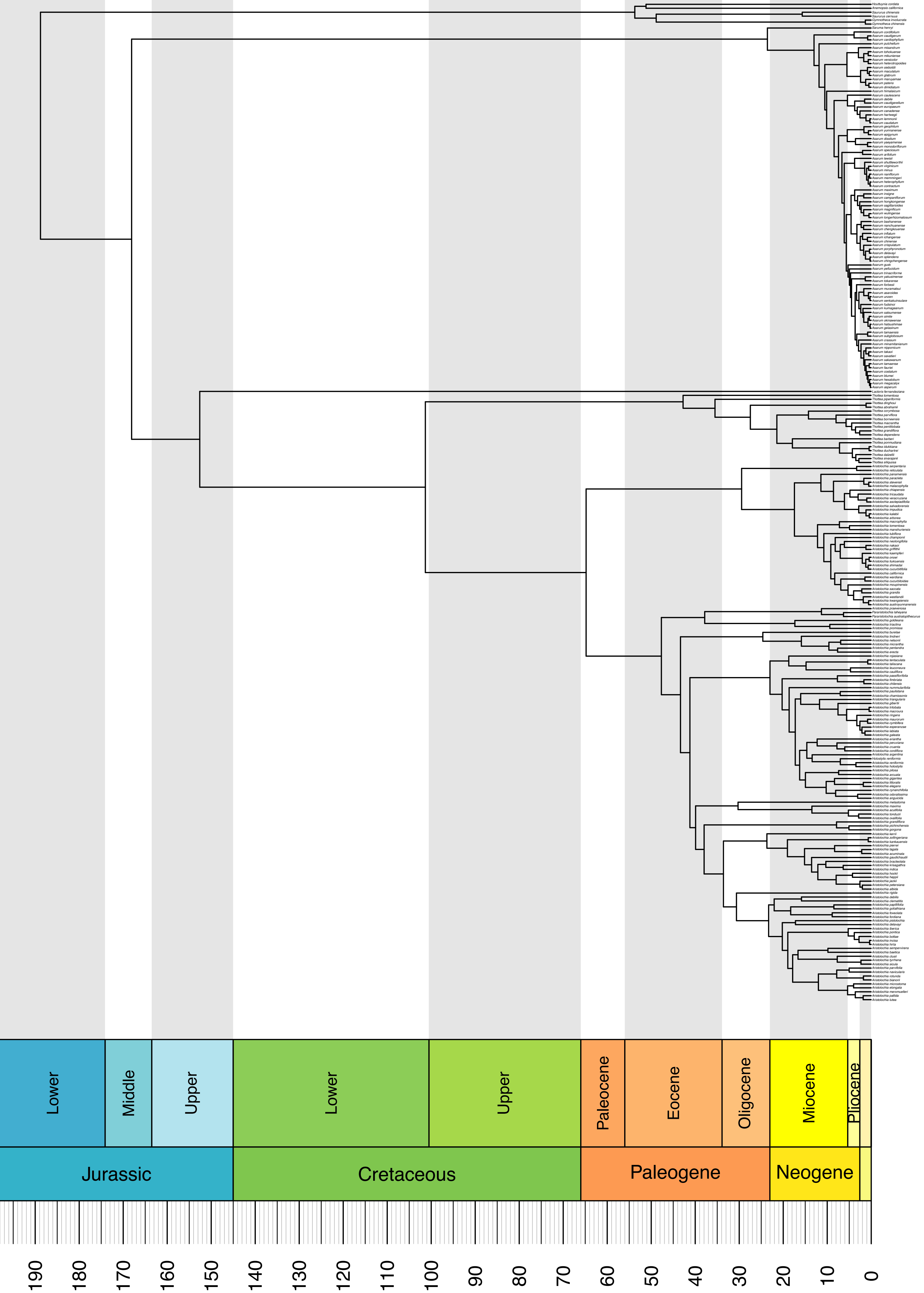

Lower
Middle
Upper
Lower
Upper
Paleocene
Eocene
Oligocene
Miocene
Pliocene
Jurassic
Cretaceous
Paleogene
Neogene
190
180
170
160
150
140
130
120
110
100
90
80
70
60
50
40
30
20
10
0

**Supplementary figure 9.** Median node ages and 95% credibility intervals (CI) for the two dating analyses of Papilionidae and the four dating analyses of Aristolochiaceae. The 95% CI overlap substantially between the two groups regardless of the dating analysis. J = Jurassic, Pl = Pliocene, P = Pleistocene.

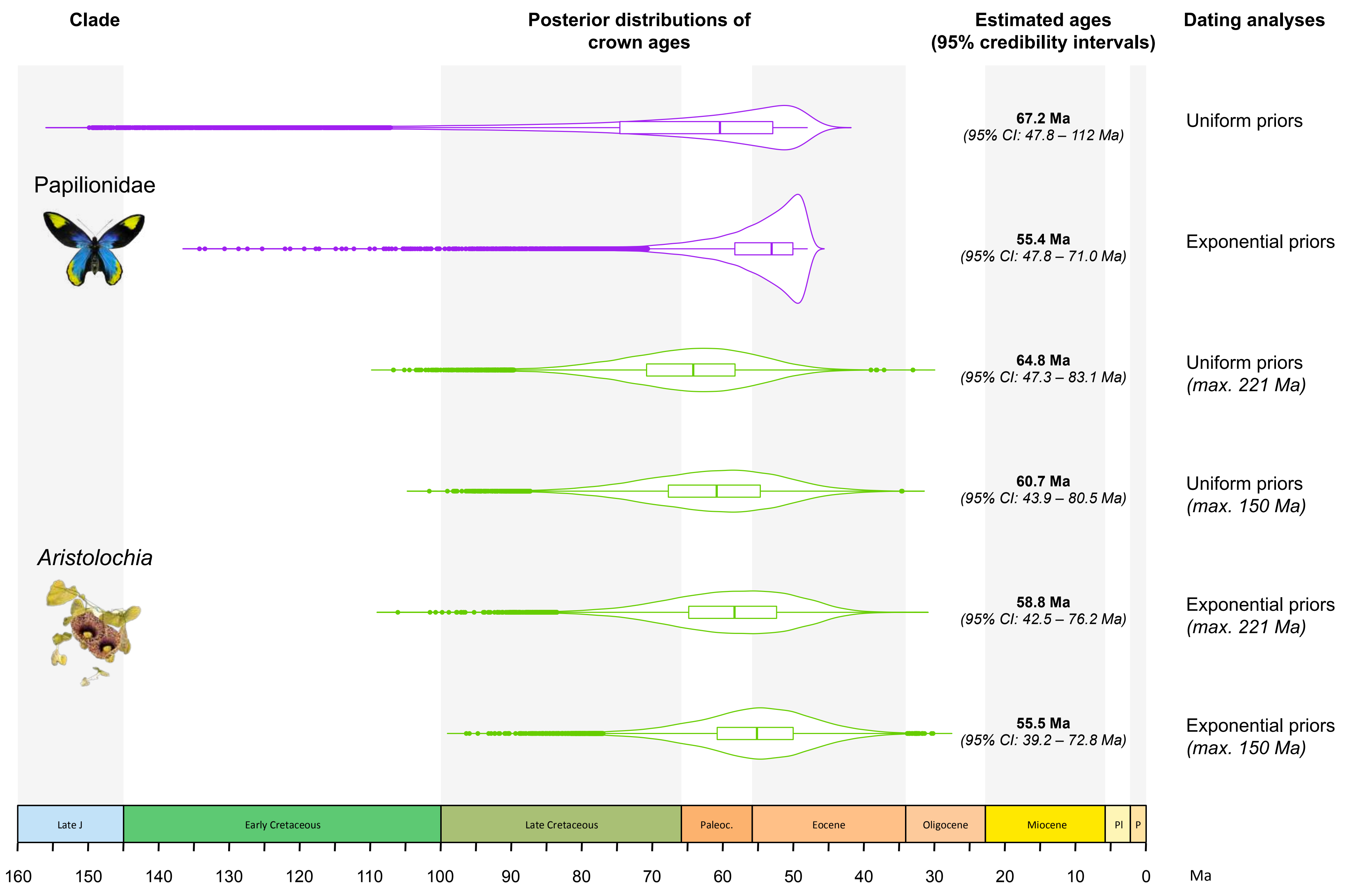
Clade
Posterior distributions of crown ages
Estimated ages (95% credibility intervals)
Dating analyses
Papilionidae
Aristolochia
67.2 Ma
(95% CI: 47.8 – 112 Ma)
Uniform priors
55.4 Ma
(95% CI: 47.8 – 71.0 Ma)
Exponential priors
64.8 Ma
(95% CI: 47.3 – 83.1 Ma)
Uniform priors
(max. 221 Ma)
60.7 Ma
(95% CI: 43.9 – 80.5 Ma)
Uniform priors
(max. 150 Ma)
58.8 Ma
(95% CI: 42.5 – 76.2 Ma)
Exponential priors
(max. 221 Ma)
55.5 Ma
(95% CI: 39.2 – 72.8 Ma)
Exponential priors
(max. 150 Ma)
Late J
Early Cretaceous
Late Cretaceous
Paleoc.
Eocene
Oligocene
Miocene
Pl
P
160
150
140
130
120
110
100
90
80
70
60
50
40
30
20
10
0
Ma

**Supplementary figure 10.** Estimation of the historical biogeography for the two molecular dated trees of Papilionidae with the Dispersal-Extinction-Cladogenesis (DEC) model. For each tree, two DEC analyses were performed: one with time-stratified palaeogeographic constraints, and one without such constraints. The swallowtail butterflies originated in a northern region centred around the Bering land bridge. K = Cretaceous, Pl = Pliocene, P = Pleistocene.

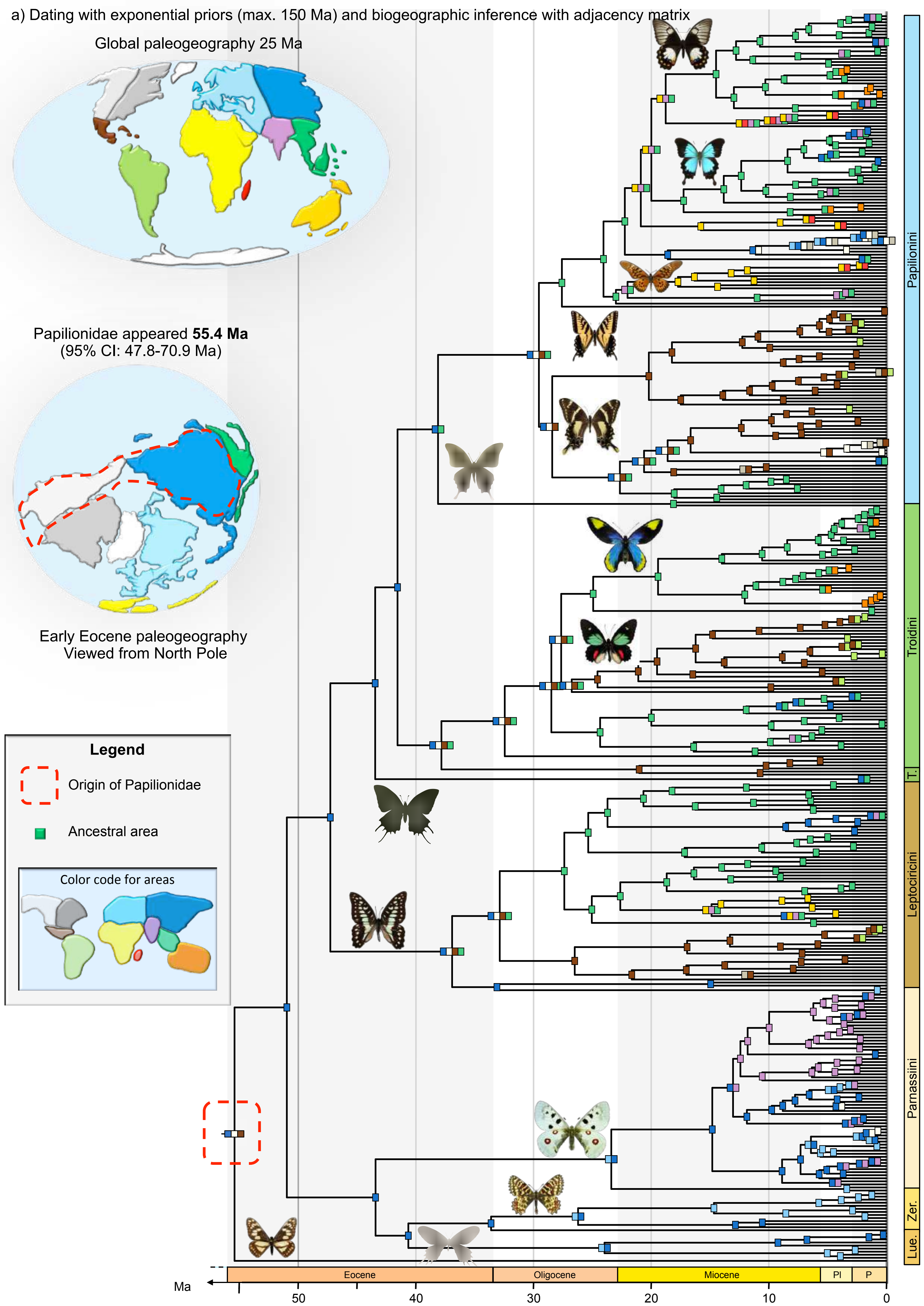

a) Dating with exponential priors (max. 150 Ma) and biogeographic inference with adjacency matrix
Global paleogeography 25 Ma
Papilionidae appeared **55.4 Ma**
(95% CI: 47.8-70.9 Ma)
Early Eocene paleogeography
Viewed from North Pole
Legend
Origin of Papilionidae
Ancestral area
Color code for areas
Papilionini
Troidini
T.
Leptocircini
Parnassiini
Zer.
Lue.
Eocene
Oligocene
Miocene
Pl
P
Ma
50
40
30
20
10
0

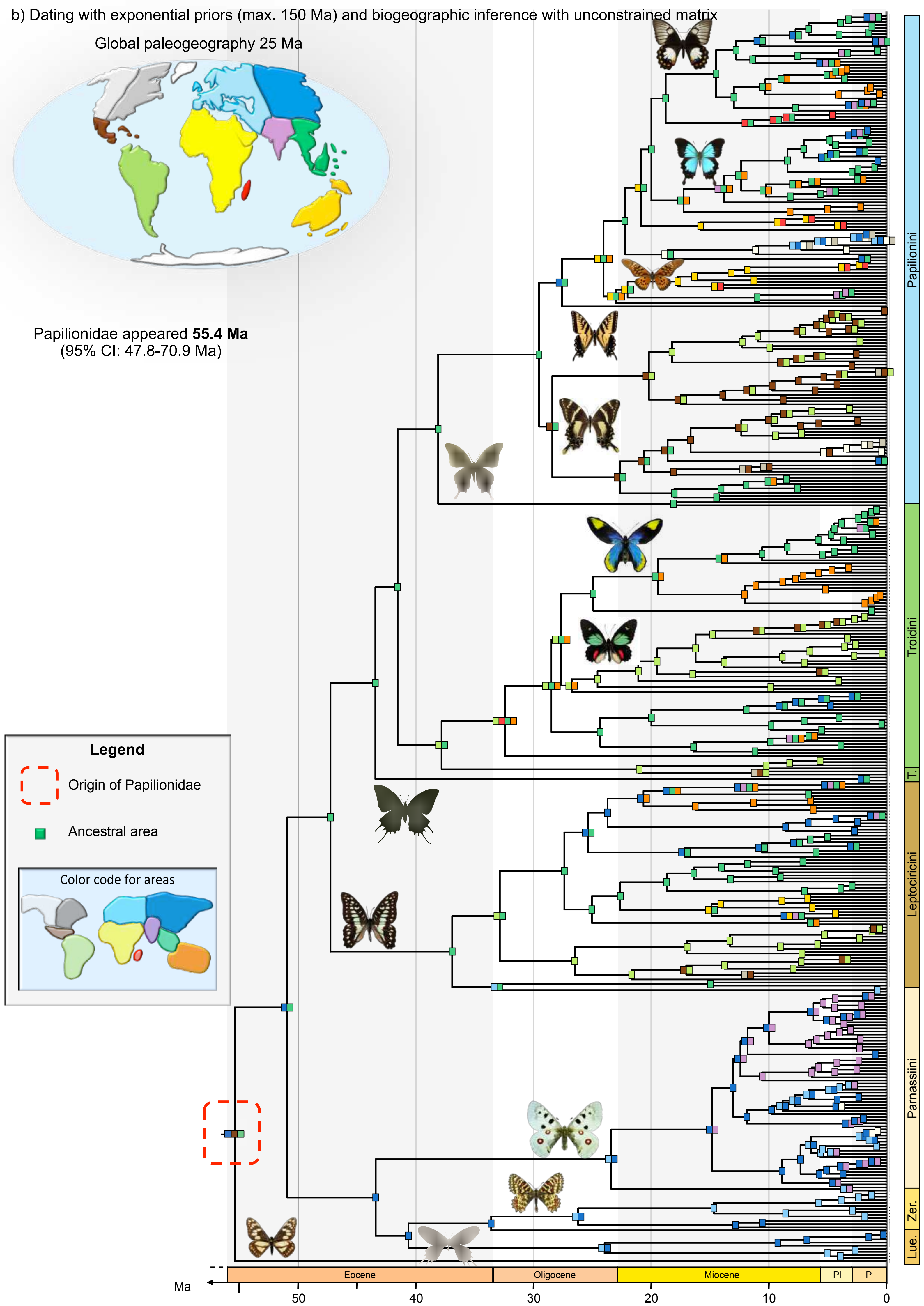
b) Dating with exponential priors (max. 150 Ma) and biogeographic inference with unconstrained matrix
Global paleogeography 25 Ma
Papilionidae appeared 55.4 Ma
(95% CI: 47.8-70.9 Ma)
Legend
Origin of Papilionidae
Ancestral area
Color code for areas
Papilionini
Troidini
T.
Leptocircini
Parnassiini
Zer.
Lue.
Eocene
Oligocene
Miocene
Pl
P
Ma
50
40
30
20
10
0

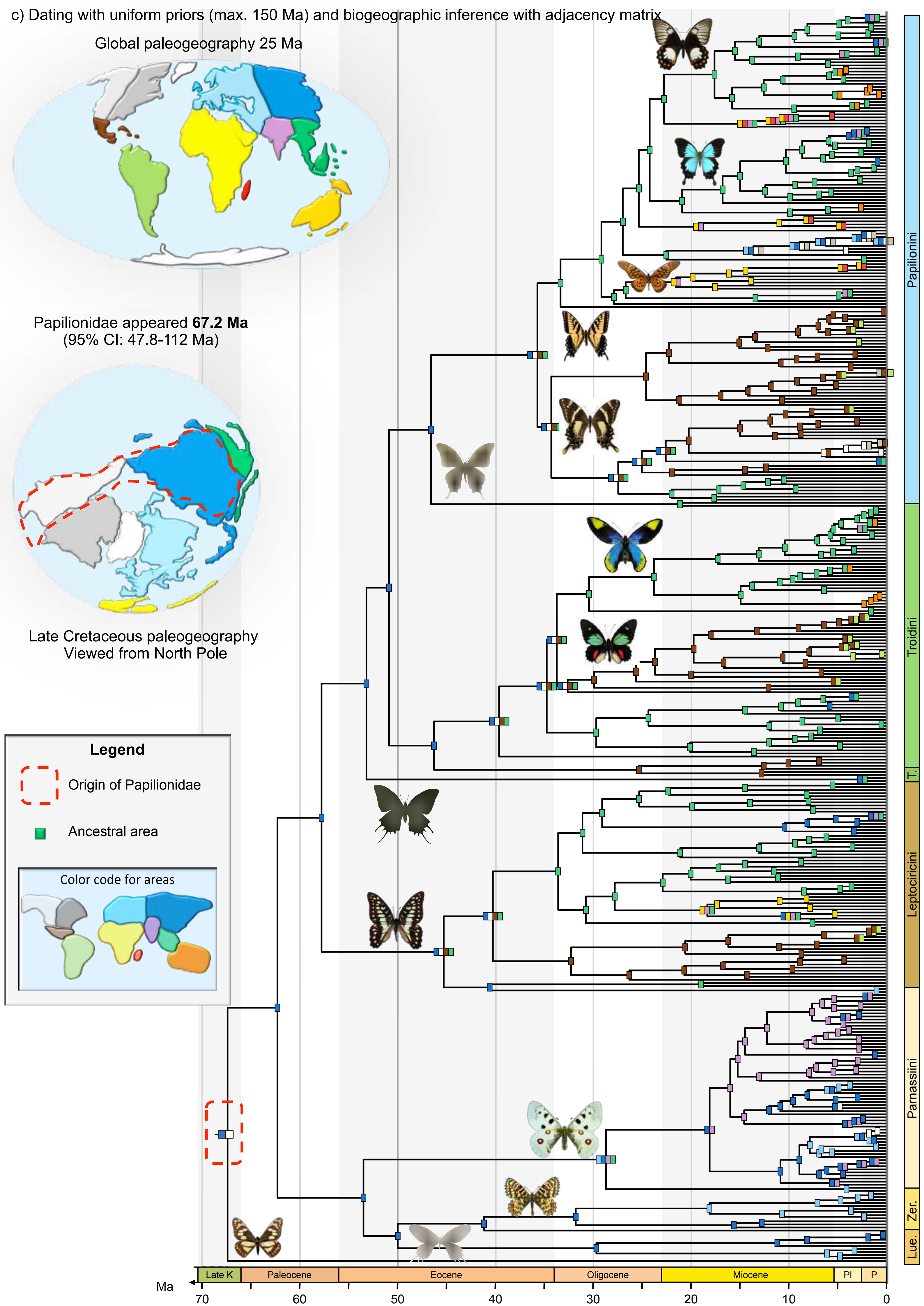

c) Dating with uniform priors (max. 150 Ma) and biogeographic inference with adjacency matrix
Global paleogeography 25 Ma
Papilionidae appeared 67.2 Ma
(95% CI: 47.8-112 Ma)
Late Cretaceous paleogeography
Viewed from North Pole
Legend
Origin of Papilionidae
Ancestral area
Color code for areas
Papilionini
Troidini
T.
Leptocircini
Parnassiini
Zer.
Lue.
Late K
Paleocene
Eocene
Oligocene
Miocene
Pl
P
Ma
70
60
50
40
30
20
10
0

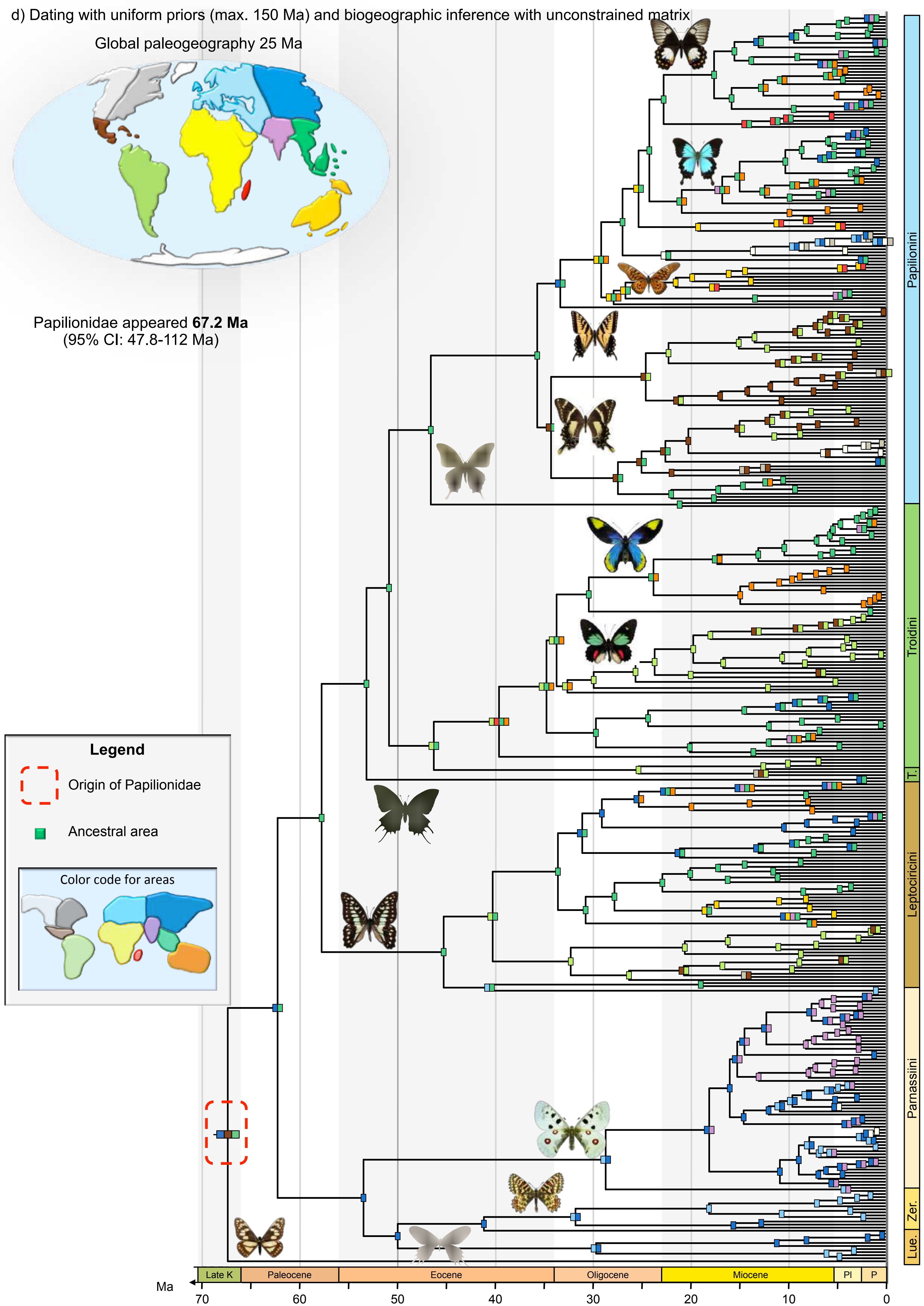

d) Dating with uniform priors (max. 150 Ma) and biogeographic inference with unconstrained matrix
Global paleogeography 25 Ma
Papilionidae appeared **67.2 Ma**
(95% CI: 47.8-112 Ma)
Legend
Origin of Papilionidae
Ancestral area
Color code for areas
Papilionini
Troidini
T.
Leptocircini
Parnassiini
Zer.
Lue.
Late K
Paleocene
Eocene
Oligocene
Miocene
Pl
P
Ma
70
60
50
40
30
20
10
0

**Supplementary figure 11.** Estimation of the historical biogeography for the four molecular dated trees of Aristolochiaceae with the Dispersal-Extinction-Cladogenesis (DEC) model. For each tree, two DEC analyses were performed: one with time-stratified palaeogeographic constraints, and one without such constraints. The genus *Aristolochia* originated in a northern region centred around the Bering land bridge. J = Jurassic, K = Cretaceous, Pl = Pliocene, P = Pleistocene.

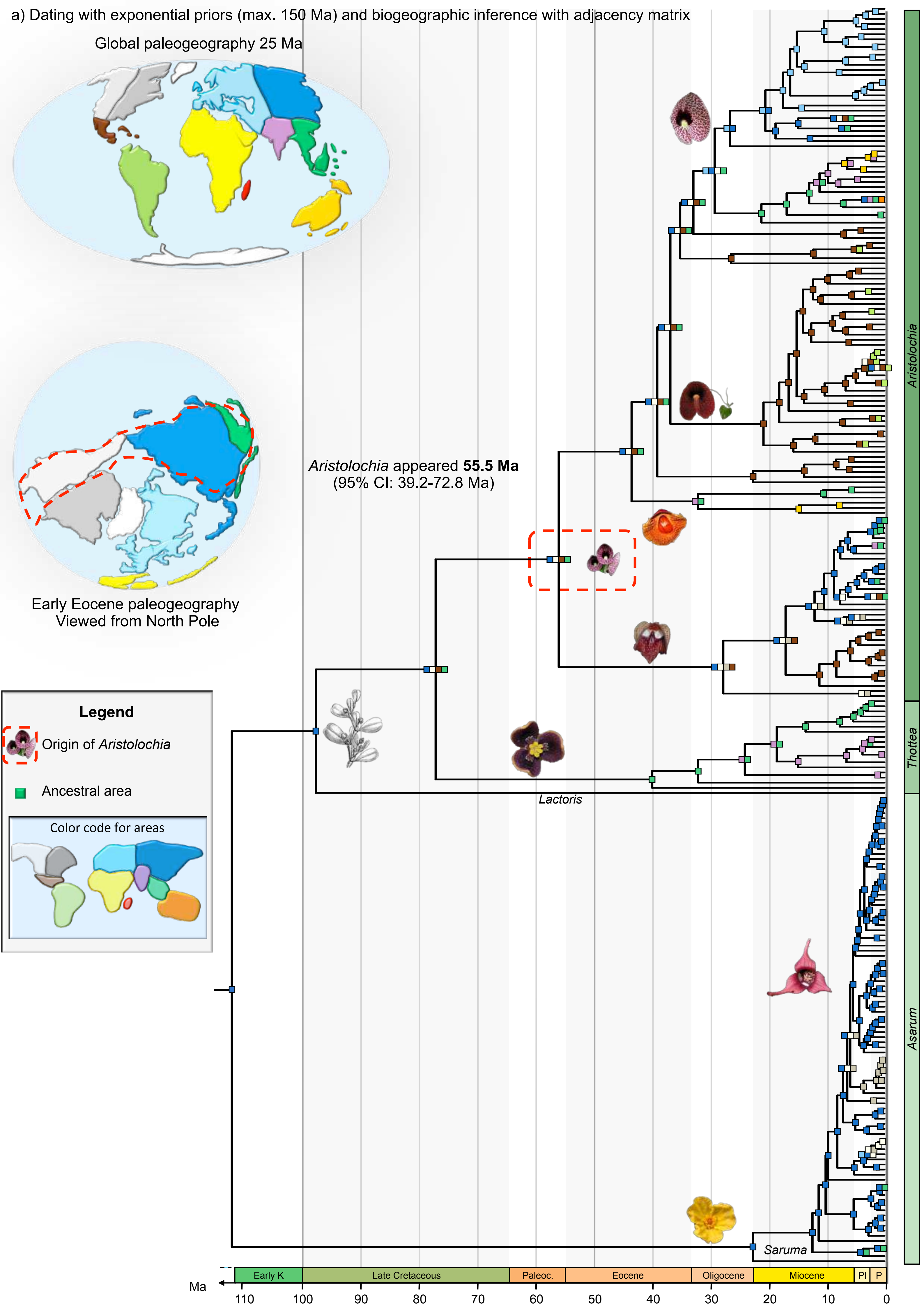
a) Dating with exponential priors (max. 150 Ma) and biogeographic inference with adjacency matrix
Global paleogeography 25 Ma
Early Eocene paleogeography
Viewed from North Pole
Aristolochia appeared 55.5 Ma
(95% CI: 39.2-72.8 Ma)
Legend
Origin of Aristolochia
Ancestral area
Color code for areas
Lactoris
Saruma
Aristolochia
Thottea
Asarum
Early K
Late Cretaceous
Paleoc.
Eocene
Oligocene
Miocene
Pl
P
Ma
110
100
90
80
70
60
50
40
30
20
10
0

b) Dating with exponential priors (max. 150 Ma) and biogeographic inference with unconstrained matrix

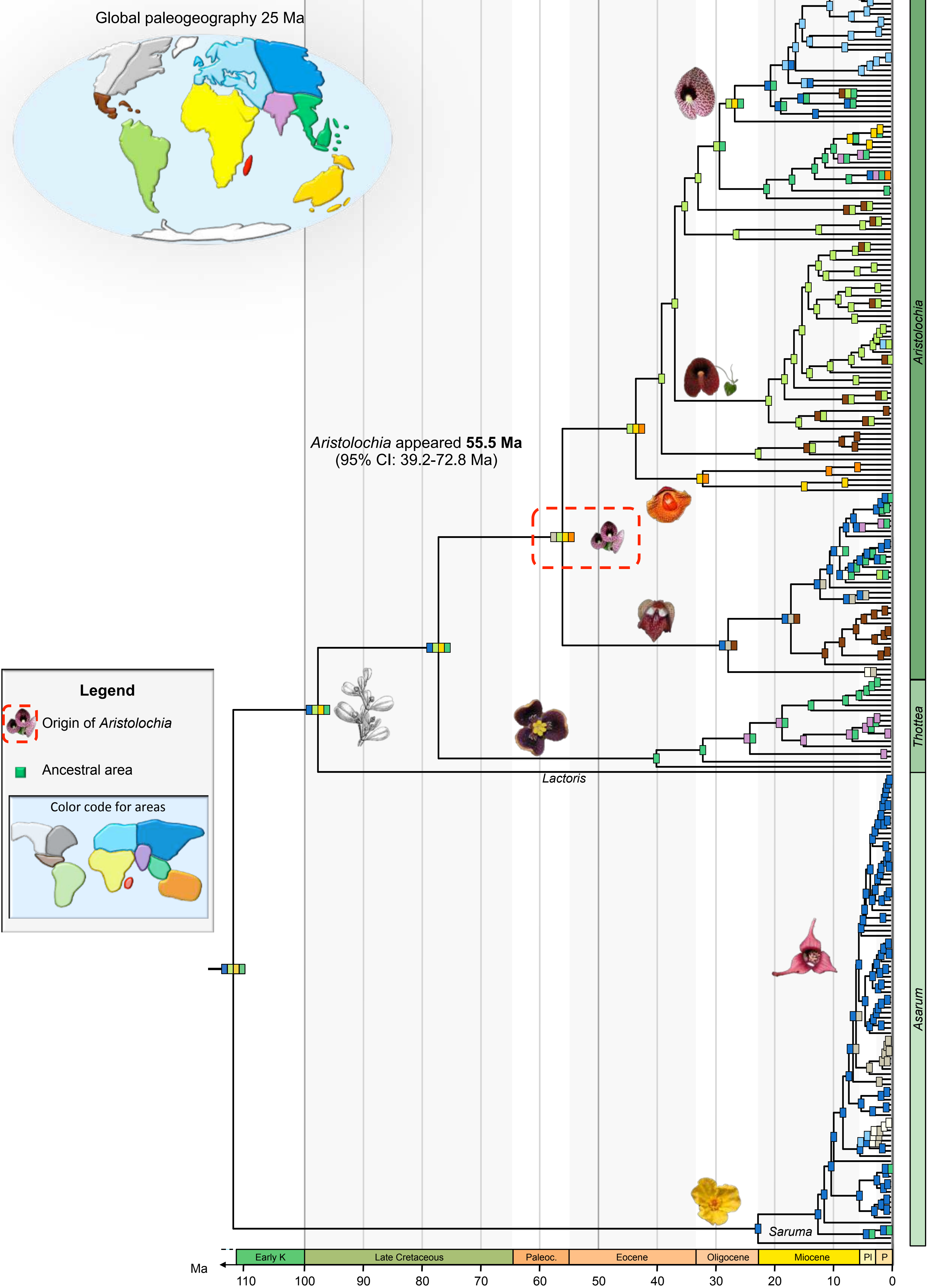

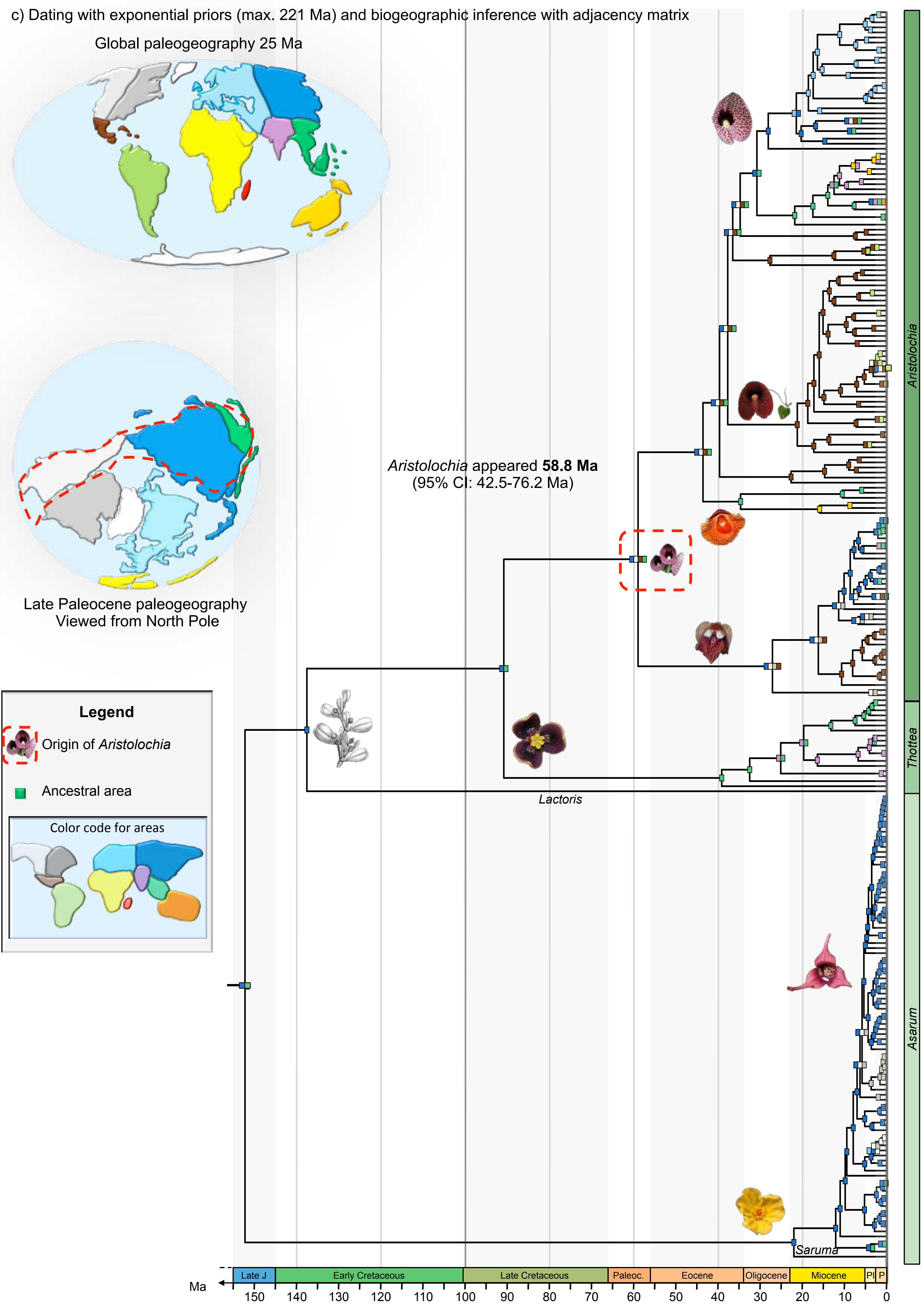

c) Dating with exponential priors (max. 221 Ma) and biogeographic inference with adjacency matrix
Global paleogeography 25 Ma
Late Paleocene paleogeography
Viewed from North Pole
Aristolochia appeared 58.8 Ma
(95% CI: 42.5-76.2 Ma)
Legend
Origin of Aristolochia
Ancestral area
Color code for areas
Lactoris
Saruma
Aristolochia
Thottea
Asarum
Late J
Early Cretaceous
Late Cretaceous
Paleoc.
Eocene
Oligocene
Miocene
Pl
P
Ma
150
140
130
120
110
100
90
80
70
60
50
40
30
20
10
0

d) Dating with exponential priors (max. 221 Ma) and biogeographic inference with unconstrained matrix

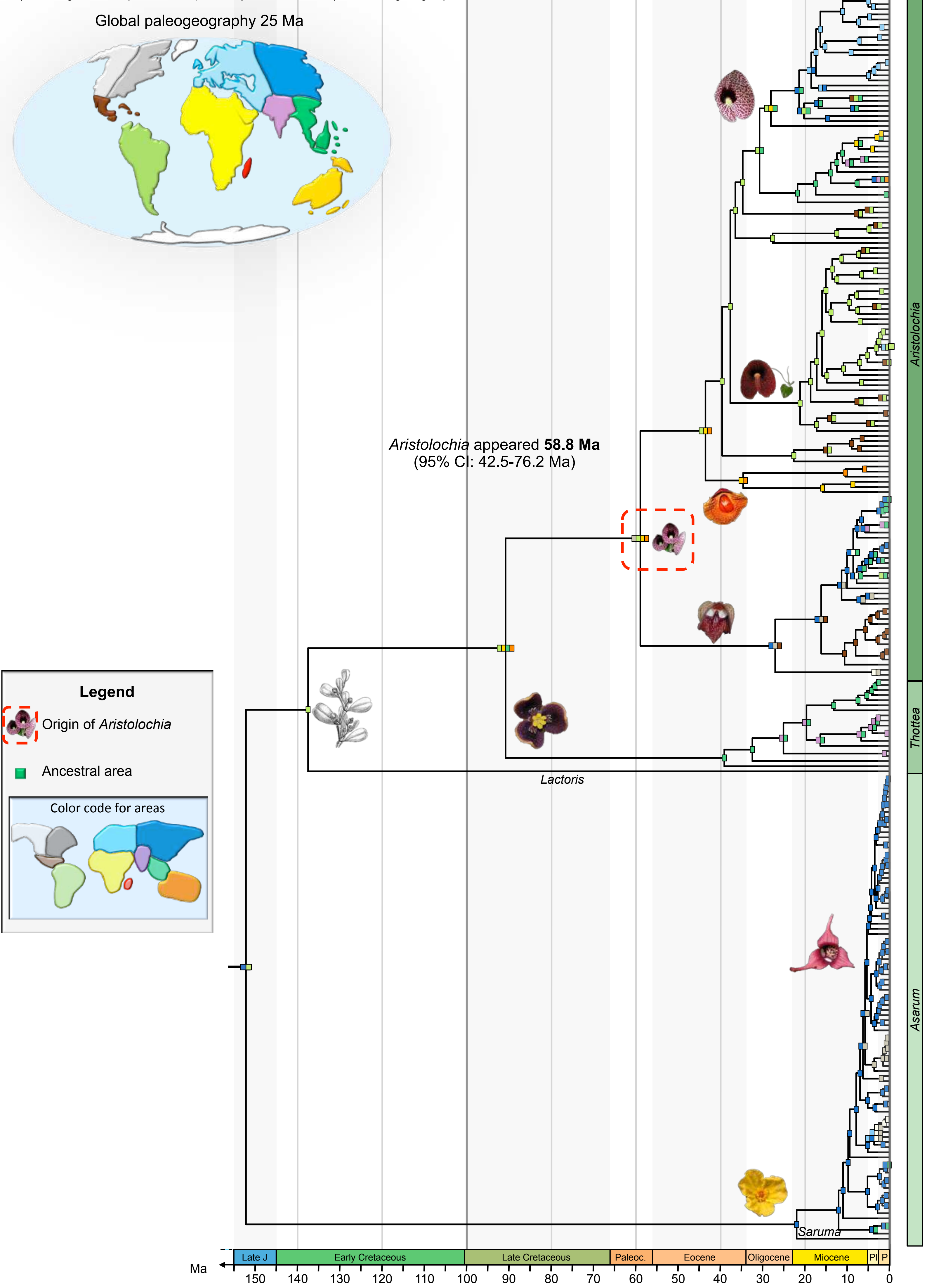

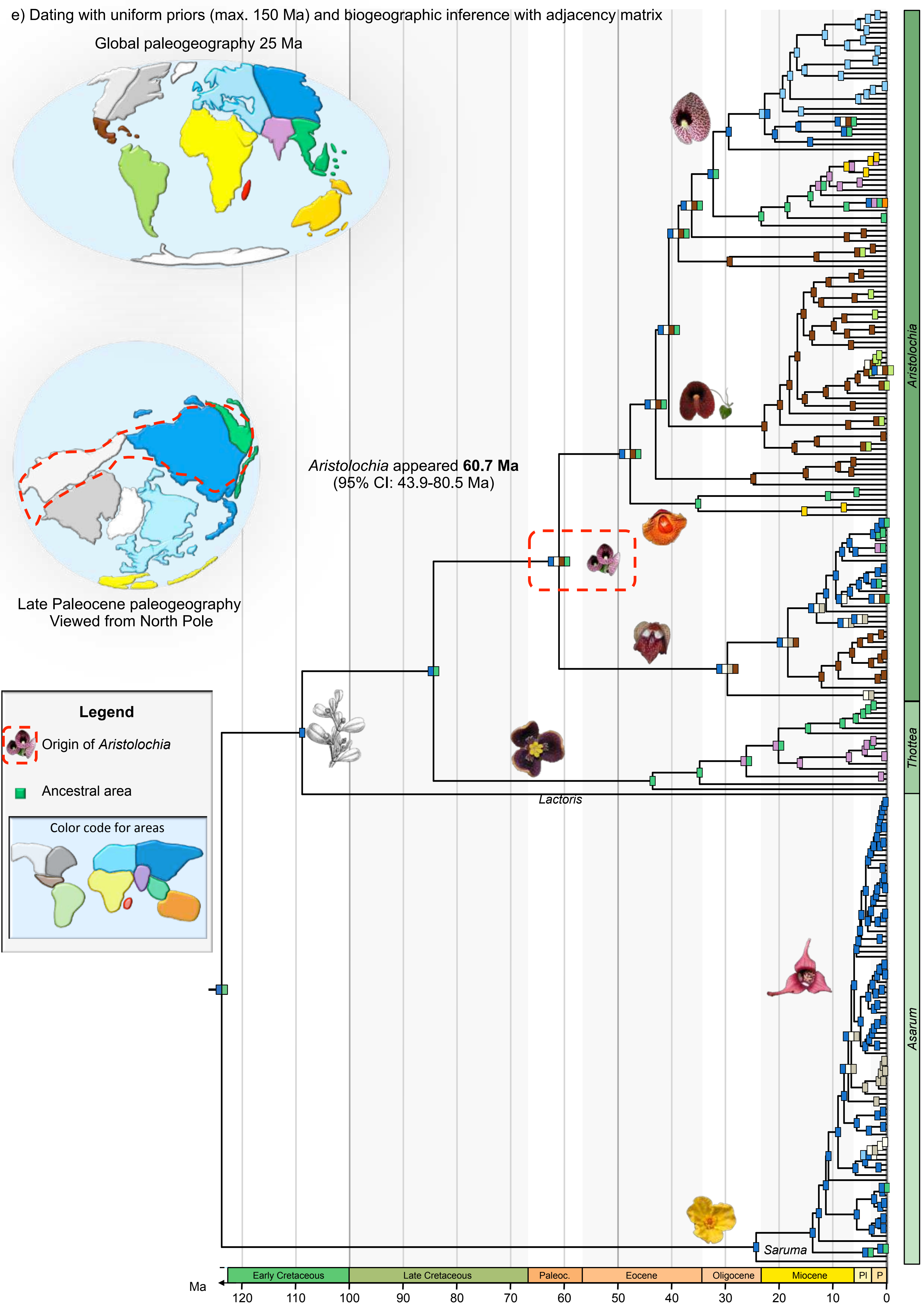

e) Dating with uniform priors (max. 150 Ma) and biogeographic inference with adjacency matrix
Global paleogeography 25 Ma
Late Paleocene paleogeography
Viewed from North Pole
Aristolochia appeared 60.7 Ma
(95% CI: 43.9-80.5 Ma)
Legend
Origin of Aristolochia
Ancestral area
Color code for areas
Lactoris
Saruma
Aristolochia
Thottea
Asarum
Early Cretaceous
Late Cretaceous
Paleoc.
Eocene
Oligocene
Miocene
Pl
P
Ma
120
110
100
90
80
70
60
50
40
30
20
10
0

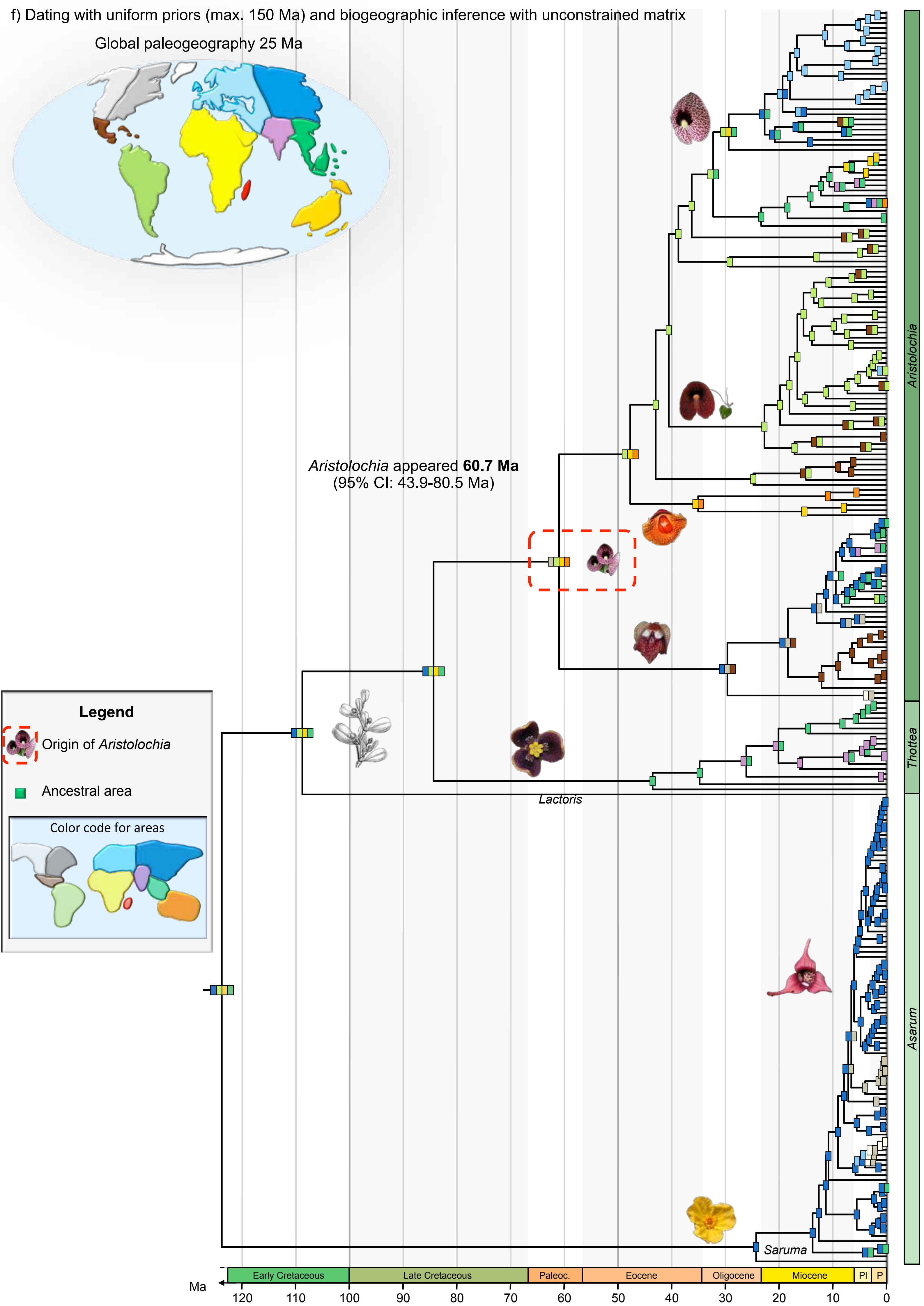
f) Dating with uniform priors (max. 150 Ma) and biogeographic inference with unconstrained matrix
Global paleogeography 25 Ma
Aristolochia appeared 60.7 Ma
(95% CI: 43.9-80.5 Ma)
Legend
Origin of Aristolochia
Ancestral area
Color code for areas
Lactoris
Saruma
Aristolochia
Thottea
Asarum
Early Cretaceous
Late Cretaceous
Paleoc.
Eocene
Oligocene
Miocene
Pl
P
Ma
120
110
100
90
80
70
60
50
40
30
20
10
0

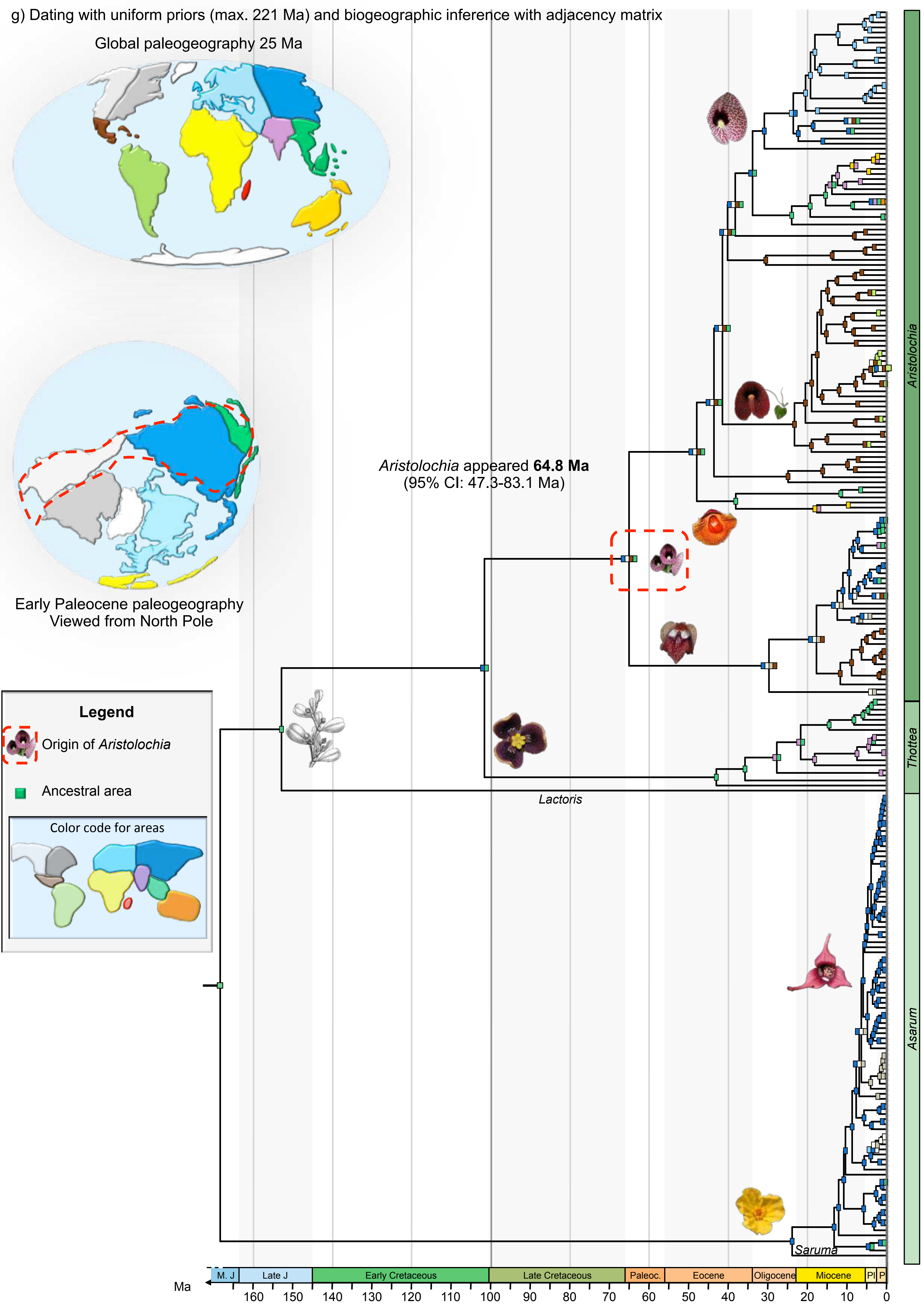

g) Dating with uniform priors (max. 221 Ma) and biogeographic inference with adjacency matrix
Global paleogeography 25 Ma
Early Paleocene paleogeography
Viewed from North Pole
Aristolochia appeared 64.8 Ma
(95% CI: 47.3-83.1 Ma)
Legend
Origin of Aristolochia
Ancestral area
Color code for areas
Lactoris
Saruma
Aristolochia
Thottea
Asarum
M. J
Late J
Early Cretaceous
Late Cretaceous
Paleoc.
Eocene
Oligocene
Miocene
Pl
P
Ma
160
150
140
130
120
110
100
90
80
70
60
50
40
30
20
10
0

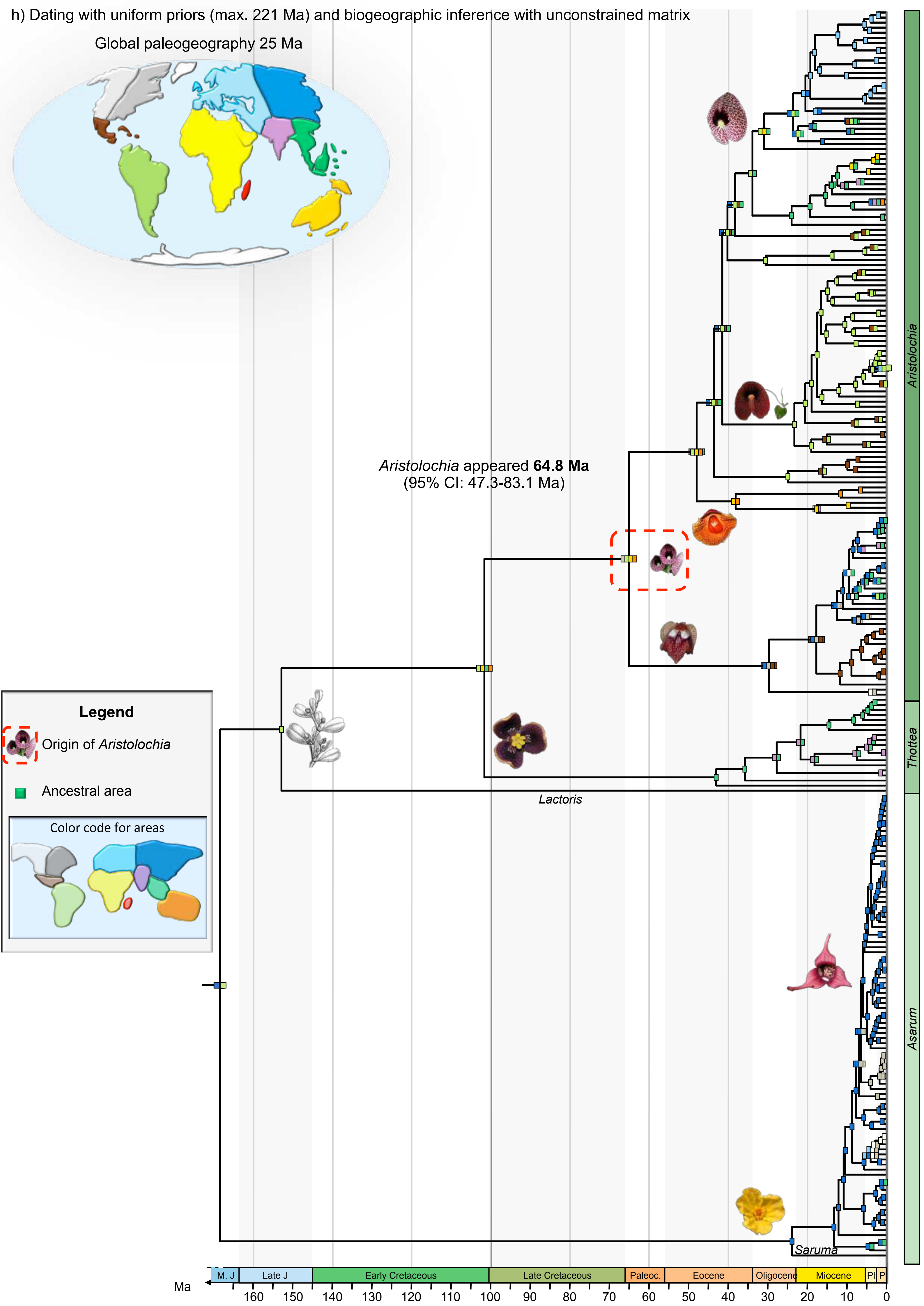

h) Dating with uniform priors (max. 221 Ma) and biogeographic inference with unconstrained matrix
Global paleogeography 25 Ma
Aristolochia appeared 64.8 Ma
(95% CI: 47.3-83.1 Ma)
Legend
Origin of Aristolochia
Ancestral area
Color code for areas
Lactoris
Saruma
Aristolochia
Thottea
Asarum
M. J
Late J
Early Cretaceous
Late Cretaceous
Paleoc.
Eocene
Oligocene
Miocene
Pl
P
Ma
160
150
140
130
120
110
100
90
80
70
60
50
40
30
20
10
0

**Supplementary figure 12.** Trait-dependent diversification of Papilionidae linked to their host plant. **a**, Bayesian inferences made with the full MuSSE model showed that speciation rates vary according to the host-plant trait. **b**, Boxplots showing the increase of diversification rates following host-plant shifts from the ancestral state (Aristolochiaceae). Only the species-poor swallowtail lineages feeding on Fabaceae, Zygophyllaceae and Magnoliaceae show decrease of diversification rates.

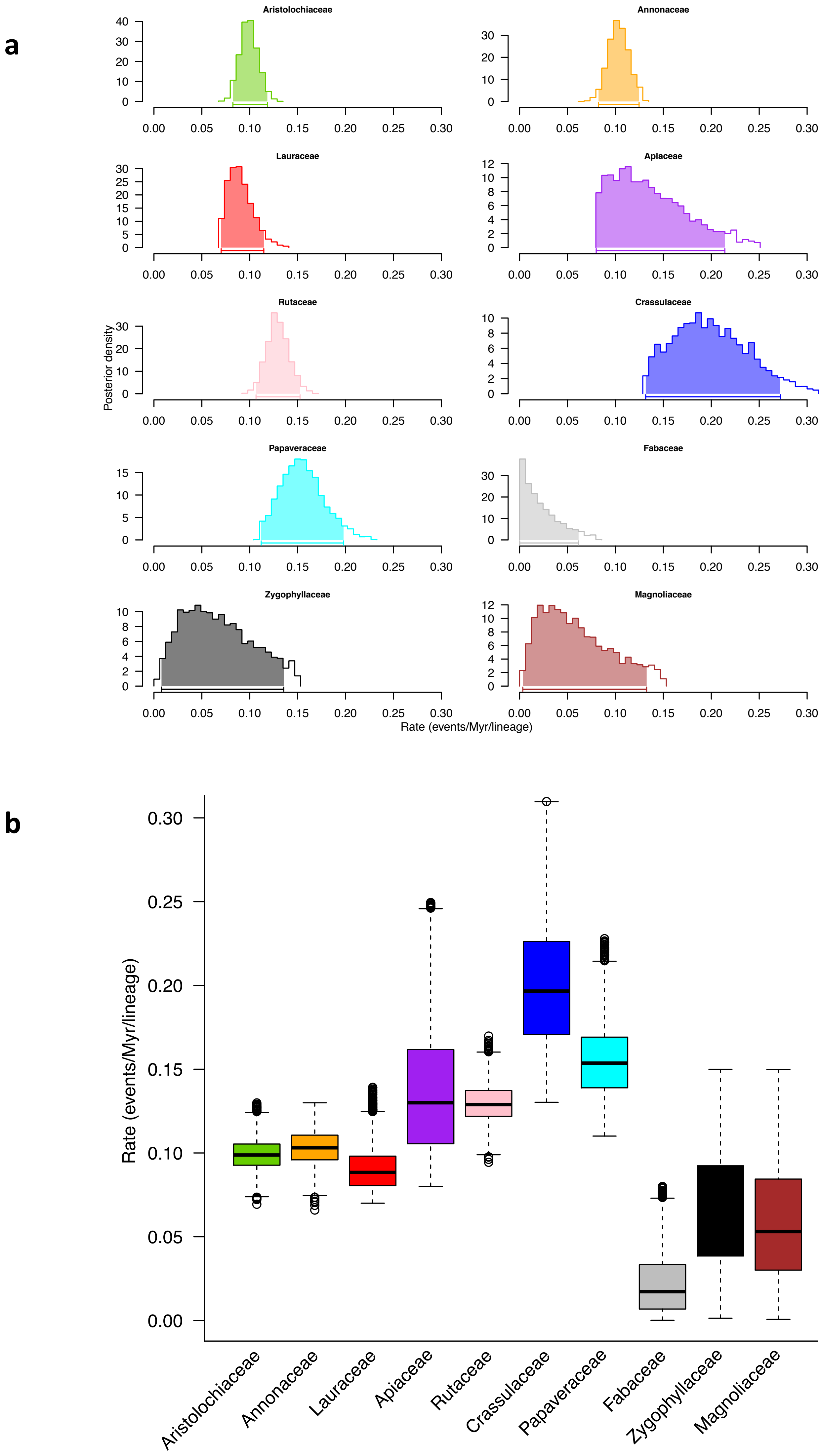
a
Aristolochiaceae
Annonaceae
Lauraceae
Apiaceae
Rutaceae
Crassulaceae
Papaveraceae
Fabaceae
Zygophyllaceae
Magnoliaceae
Posterior density
Rate (events/Myr/lineage)
b
Rate (events/Myr/lineage)
Aristolochiaceae
Annonaceae
Lauraceae
Apiaceae
Rutaceae
Crassulaceae
Papaveraceae
Fabaceae
Zygophyllaceae
Magnoliaceae

**Supplementary figure 13.** Time-dependent diversification of Papilionidae after shifting to new host plants. Diversification is inferred with the RPANDA models, and the best-fit model is plotted showing rates through time for each clade. A model with increasing diversification over time best fits the Aristolochiaceae feeders. A model with a slowdown of diversification through time explained the diversification of Annonaceae feeders, Lauraceae feeders, and Papaveraceae feeders. A model with constant rates through time best fits the diversification of Apiaceae feeders, Crassulaceae feeders, and Rutaceae feeders. K = Cretaceous, Pl = Pliocene, P = Pleistocene.

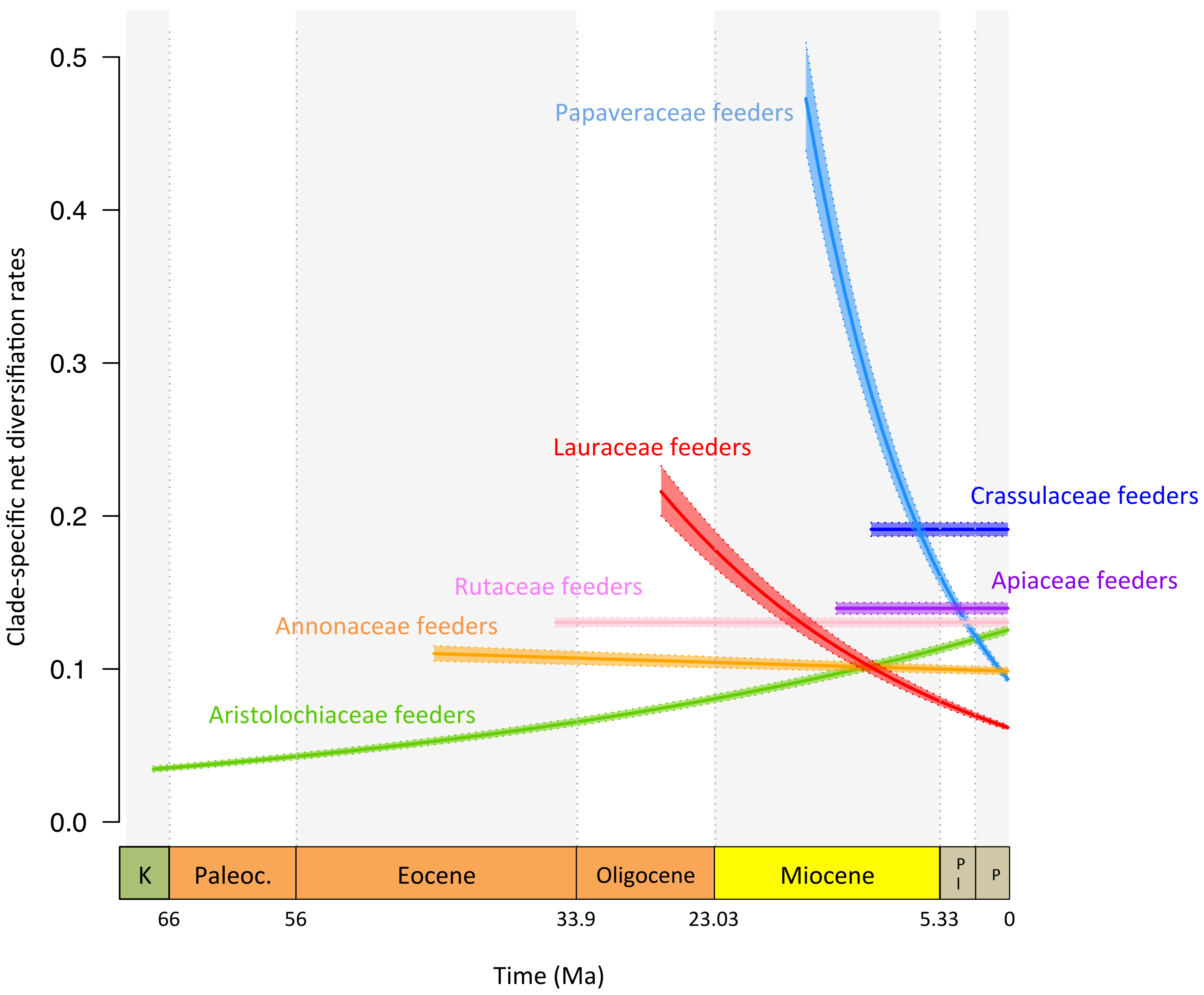

Clade-specific net diversifiation rates
0.5
0.4
0.3
0.2
0.1
0.0
Papaveraceae feeders
Lauraceae feeders
Crassulaceae feeders
Apiaceae feeders
Rutaceae feeders
Annonaceae feeders
Aristolochiaceae feeders
K
Paleoc.
Eocene
Oligocene
Miocene
P l
P
66
56
33.9
23.03
5.33
0
Time (Ma)

**Supplementary figure 14.** Bayesian analysis of clade-specific and time-dependent diversification of Papilionidae obtained with BAMM. a, Phylorate plot showing that global diversification rates increase through time in Papilionidae with no significant rate shifts detected by BAMM (the inset plot indicates the posterior probability for the estimated number of shifts). b, Rate-through-time plots for selected swallowtail lineages feeding on distinct host-plant families. The results also show an overall diversification increase through time for each group of swallowtails. P = Palaeocene, E = Eocene, O = Oligocene, M = Miocene.

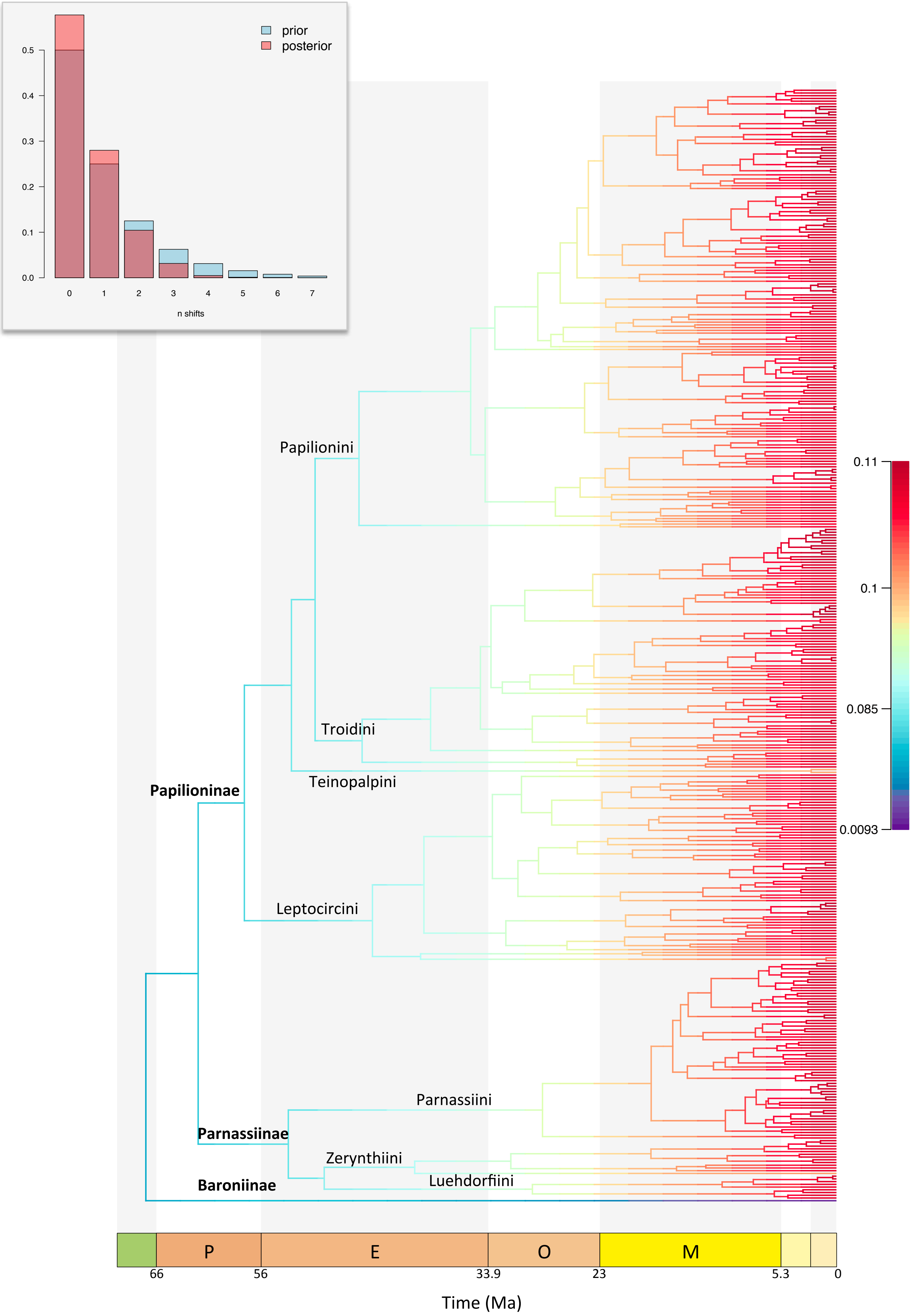

prior
posterior
0.5
0.4
0.3
0.2
0.1
0.0
0
1
2
3
4
5
6
7
n shifts
Papilionini
Troidini
Teinopalpini
Papilioninae
Leptocircini
Parnassiini
Parnassiinae
Zerynthiini
Luehdorfiini
Baroniinae
0.11
0.1
0.085
0.0093
P
E
O
M
66
56
33.9
23
5.3
0
Time (Ma)

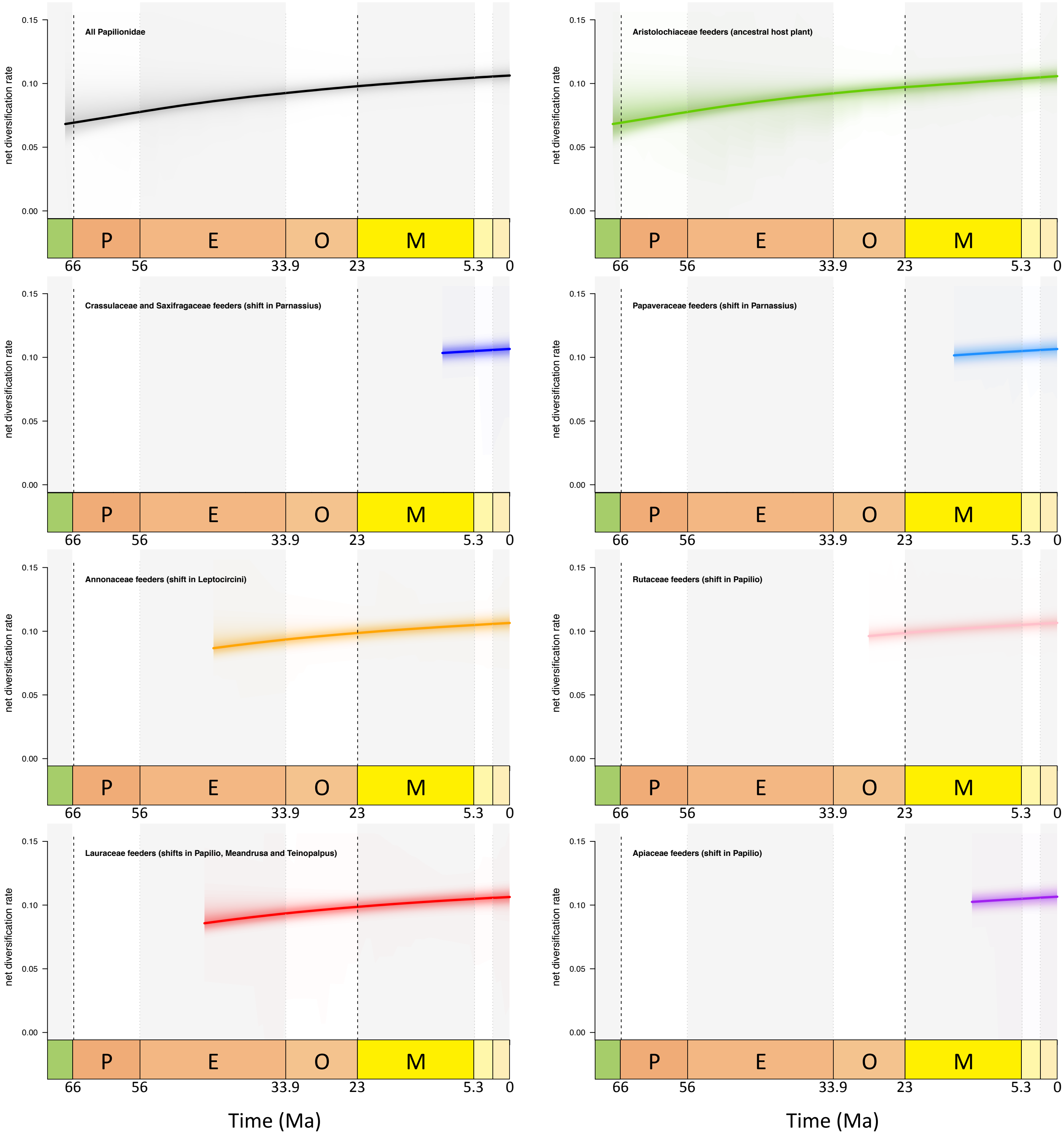

All Papilionidae
Aristolochiaceae feeders (ancestral host plant)
Crassulaceae and Saxifragaceae feeders (shift in Parnassius)
Papaveraceae feeders (shift in Parnassius)
Annonaceae feeders (shift in Leptocircini)
Rutaceae feeders (shift in Papilio)
Lauraceae feeders (shifts in Papilio, Meandrusa and Teinopalpus)
Apiaceae feeders (shift in Papilio)
net diversification rate
0.15
0.10
0.05
0.00
P
E
O
M
66
56
33.9
23
5.3
0
Time (Ma)

**Supplementary figure 15.** Bayesian analysis of branch-specific and time-dependent diversification of Papilionidae obtained with RevBayes. The median rates of diversification are plotted along each branch of the phylogeny, which shows a global increase of diversification rates through time in Papilionidae. Contrary to BAMM, this approach detected shifts in diversification rates in particular within the genera *Parnassius* and *Papilio* that have both shifted to new host-plant families. P = Palaeocene, E = Eocene, O = Oligocene, M = Miocene.

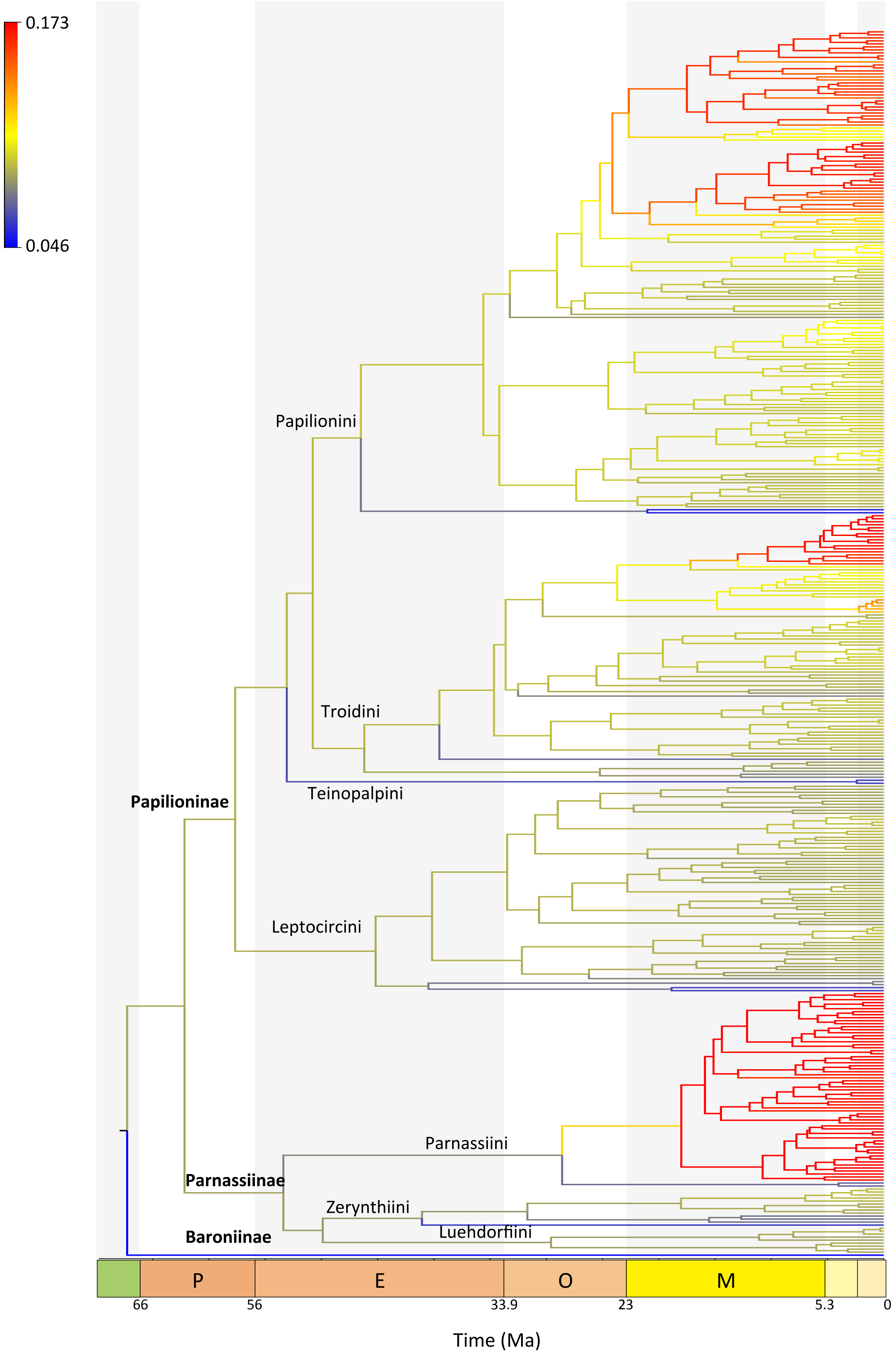

0.173
0.046
Papilionini
Troidini
Teinopalpini
Papilioninae
Leptocircini
Parnassiini
Parnassiinae
Zerynthiini
Luehdorfiini
Baroniinae
P
E
O
M
66
56
33.9
23
5.3
0
Time (Ma)

**Supplementary figure 16.** Bayesian analysis of episodic diversification of Papilionidae obtained with CoMET. The four plots represent speciation, extinction, net diversification, and relative extinction rates through time for the whole family. The result indicates a global increase of diversification rates over time, notably starting ~40 Ma. P = Palaeocene, E = Eocene, O = Oligocene, M = Miocene.

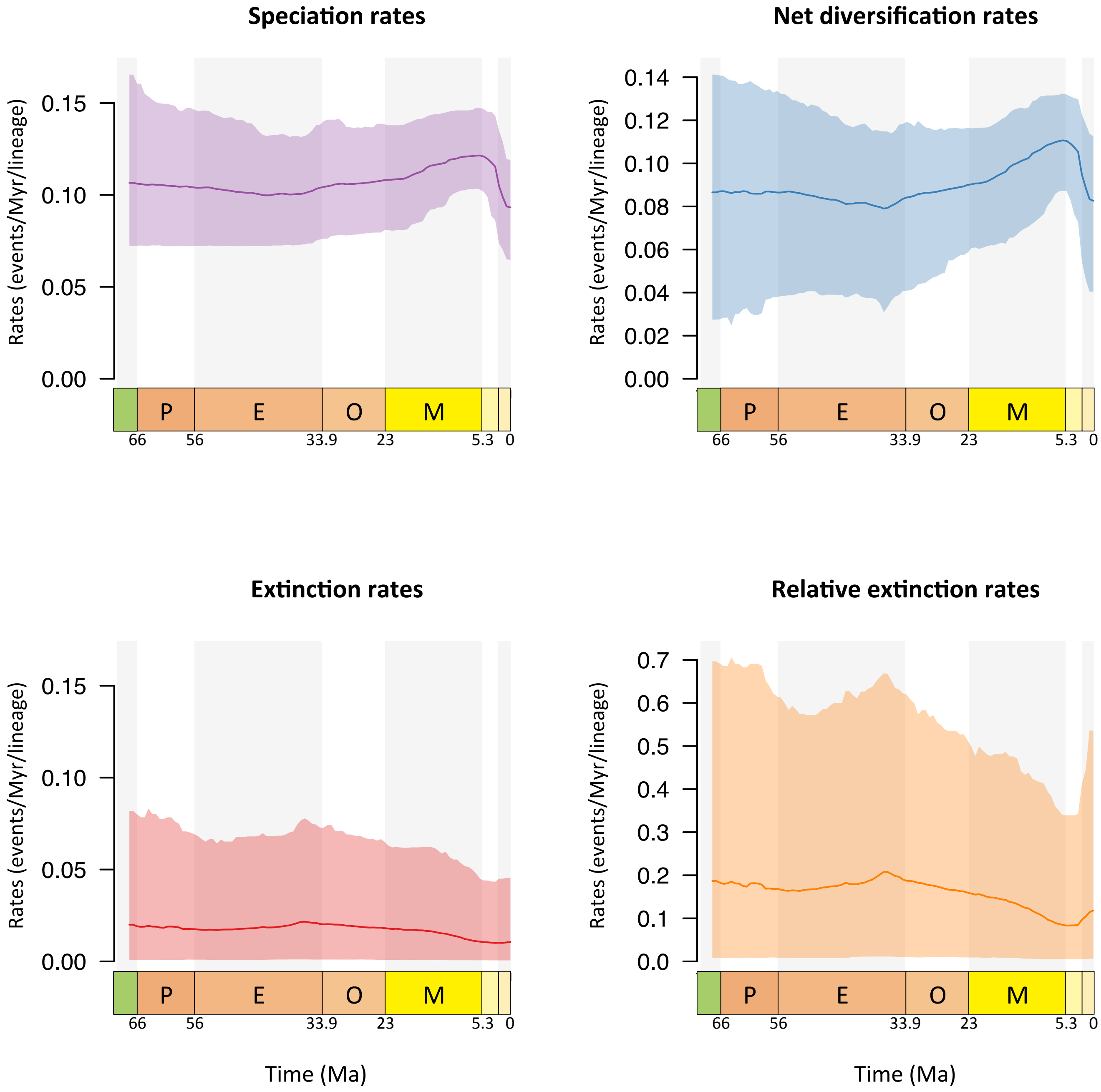

Speciation rates
Net diversification rates
Extinction rates
Relative extinction rates
Rates (events/Myr/lineage)
0.15
0.10
0.05
0.00
0.14
0.12
0.10
0.08
0.06
0.04
0.02
0.00
0.7
0.6
0.5
0.4
0.3
0.2
0.1
0.0
P
E
O
M
66
56
33.9
23
5.3
0
Time (Ma)

**Supplementary figure 17.** Number of host plants consumed through time by Papilionidae. Using the estimation of ancestral host-plant preferences (Fig. S4), we plotted the time at which a new host-plant family was colonised. This result shows that the swallowtail butterflies have a steady increase in the number of host families consumed over time. This ecological diversification can be paralleled with the global increase in diversification rates estimated by birth-death models (Figs. S13-16). K = Cretaceous, Pl = Pliocene, P = Pleistocene.

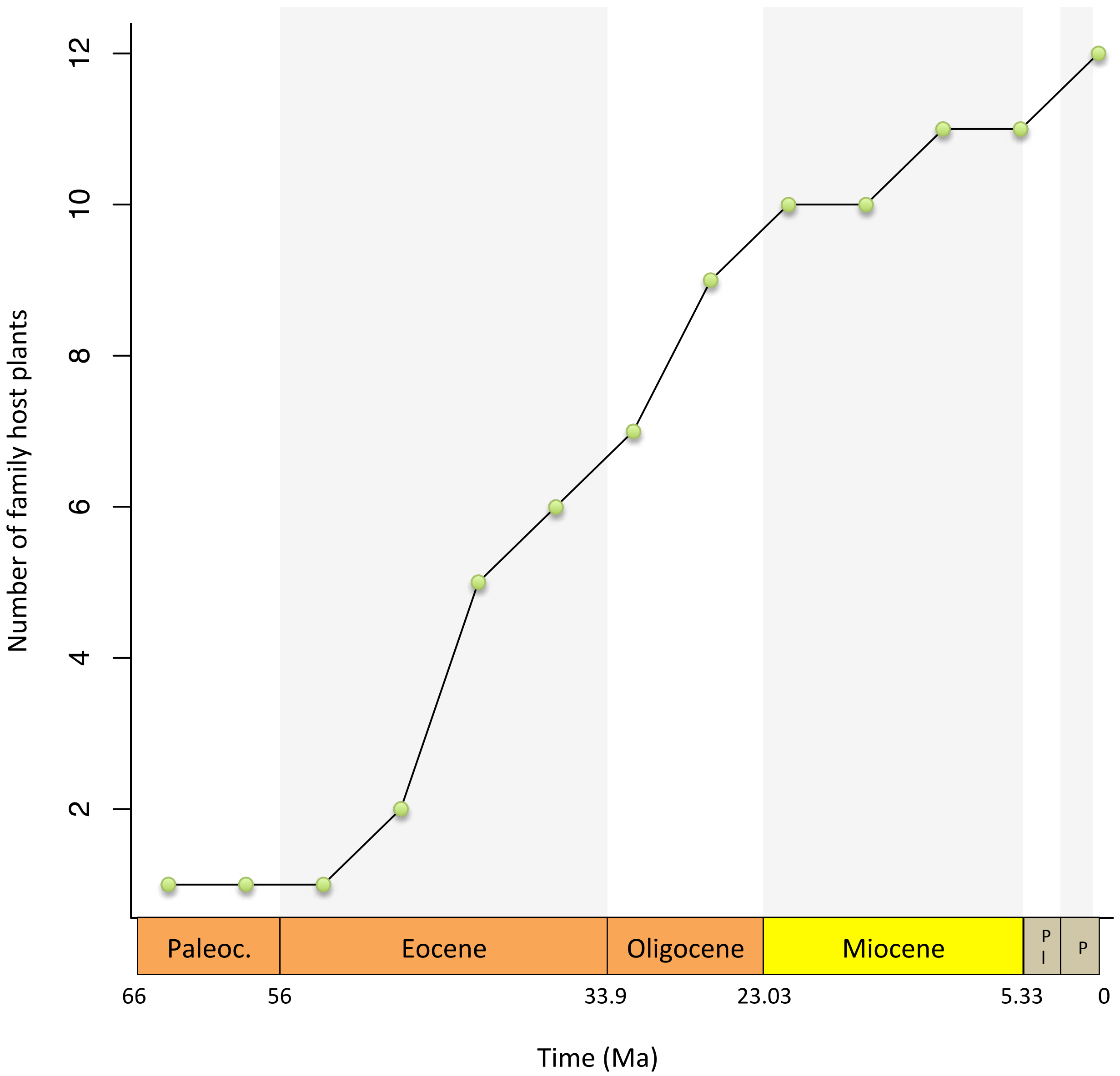

Number of family host plants
12
10
8
6
4
2
Paleoc.
Eocene
Oligocene
Miocene
P
l
P
66
56
33.9
23.03
5.33
0
Time (Ma)

**Supplementary figure 18.** Genus-level phylogenomic tree of Papilionidae showing the 14 selected branches with host-plant shifts and the 14 selected branches without host-plant shifts (control branches). The selection of these branches is based on the estimation of ancestral state models using the species-level phylogenies and current host-plant preferences (Figs. S4, S5).

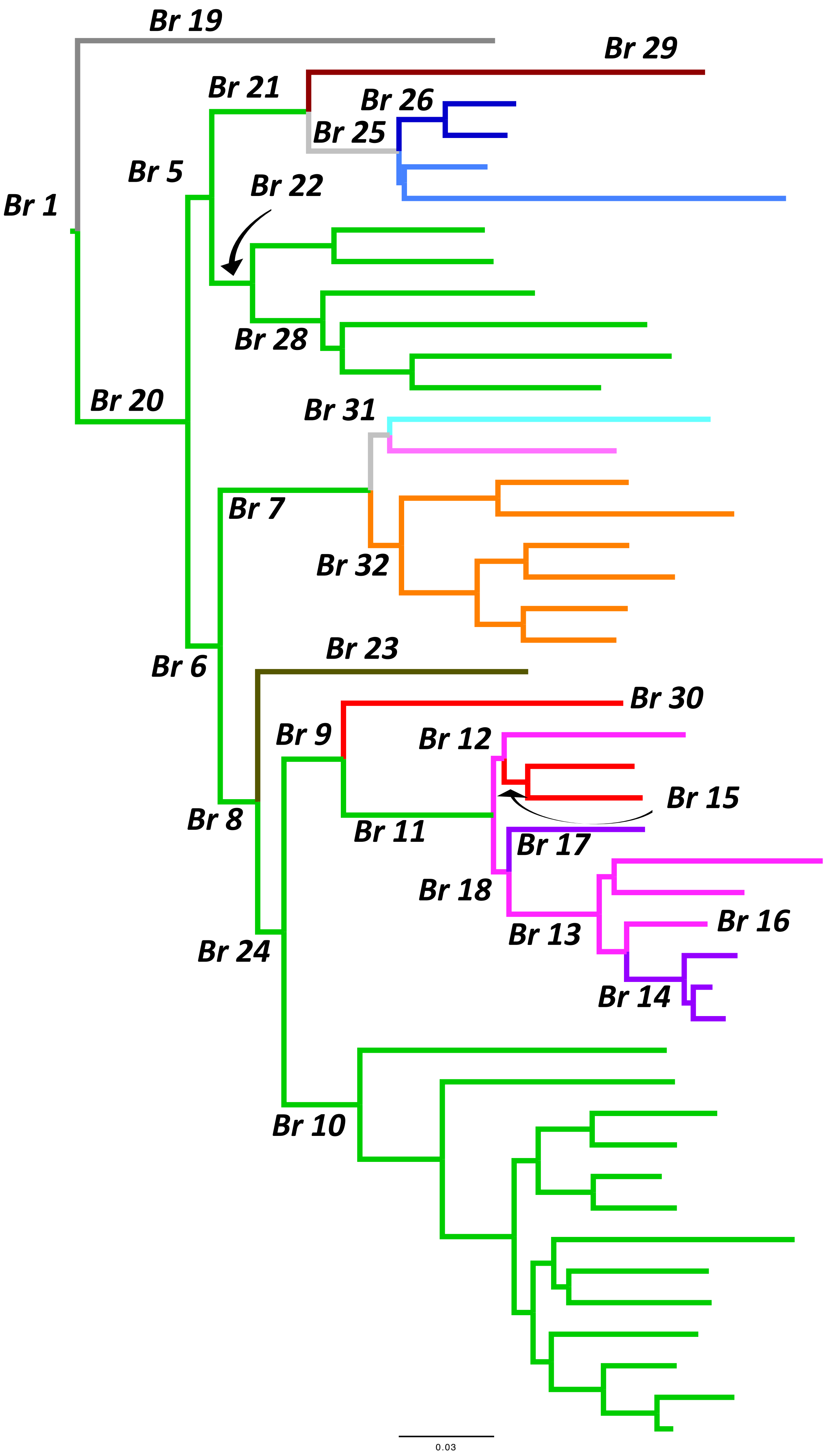

Br 19
Br 29
Br 21
Br 26
Br 25
Br 5
Br 1
Br 22
Br 28
Br 20
Br 31
Br 7
Br 32
Br 6
Br 23
Br 30
Br 9
Br 12
Br 15
Br 8
Br 11
Br 17
Br 18
Br 13
Br 16
Br 24
Br 14
Br 10
0.03

| ***Branch ID*** | ***Plant shift*** | ***Climate shift*** | ***Plant and climate shifts*** |
|---|---|---|---|
| Branch1 | Host-plant shift | No shift | Host shift only |
| Branch5 | No shift | No shift | No shift |
| Branch6 | No shift | No shift | No shift |
| Branch7 | Host-plant shift | No shift | Host shift only |
| Branch8 | No shift | No shift | No shift |
| Branch9 | Host-plant shift | No shift | Host shift only |
| Branch10 | No shift | No shift | No shift |
| Branch11 | Host-plant shift | No shift | Host shift only |
| Branch12 | No shift | No shift | No shift |
| Branch13 | No shift | No shift | No shift |
| Branch14 | Host-plant shift | Climate shift | Host and climate shifts |
| Branch15 | Host-plant shift | Climate shift | Host and climate shifts |
| Branch16 | No shift | No shift | No shift |
| Branch17 | Host-plant shift | Climate shift | Host and climate shifts |
| Branch18 | No shift | No shift | No shift |
| Branch19 | Host-plant shift | No shift | Host shift only |
| Branch20 | No shift | No shift | No shift |
| Branch21 | Host-plant shift | No shift | Host shift only |
| Branch22 | No shift | No shift | No shift |
| Branch23 | Host-plant shift | No shift | Host shift only |
| Branch24 | No shift | No shift | No shift |
| Branch25 | Host-plant shift | Climate shift | Host and climate shifts |
| Branch26 | No shift | No shift | No shift |
| Branch28 | No shift | No shift | No shift |
| Branch29 | Host-plant shift | No shift | Host shift only |
| Branch30 | Host-plant shift | No shift | Host shift only |
| Branch31 | Host-plant shift | Climate shift | Host and climate shifts |
| Branch32 | No shift | No shift | No shift |

**Supplementary figure 19.** Violin plots of the percentage of missing data ("N" or "-") and proportion of GC at third codon position (GC3) in alignment were positive selection have been detected ("Yes") and positive selection have not been detected ("No"). Panels a and b are dataset 1 with 520 genes, and panels c and d are dataset 2 with 1533 genes.

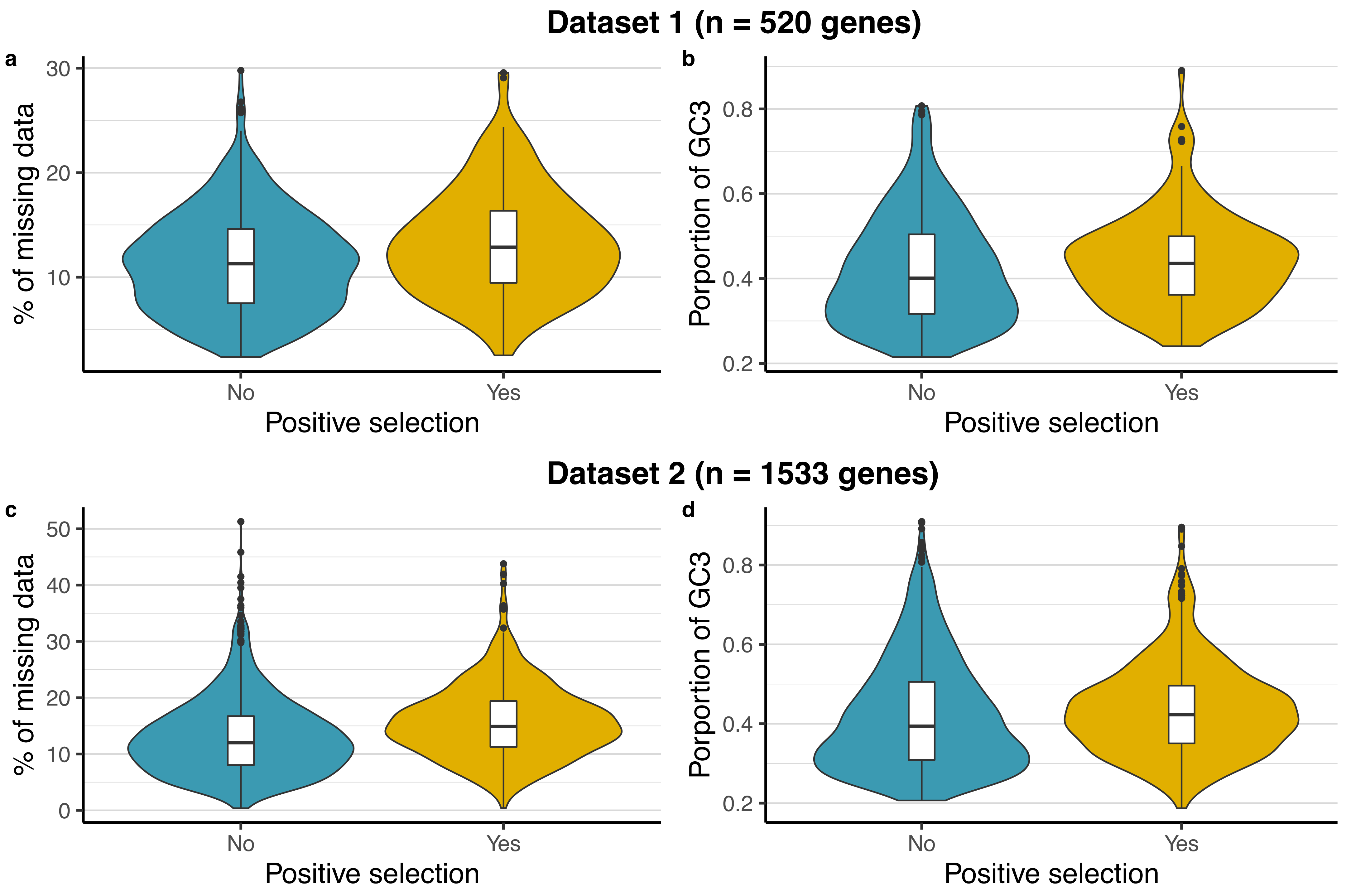
Dataset 1 (n = 520 genes)
a
% of missing data
30
20
10
No
Yes
Positive selection
b
Porportion of GC3
0.8
0.6
0.4
0.2
No
Yes
Positive selection
Dataset 2 (n = 1533 genes)
c
% of missing data
50
40
30
20
10
0
No
Yes
Positive selection
d
Porportion of GC3
0.8
0.6
0.4
0.2
No
Yes
Positive selection

**Supplementary figure 20.** The percentage of missing data ("N" or "-") per genes across species computed for dataset 1 and dataset 2.

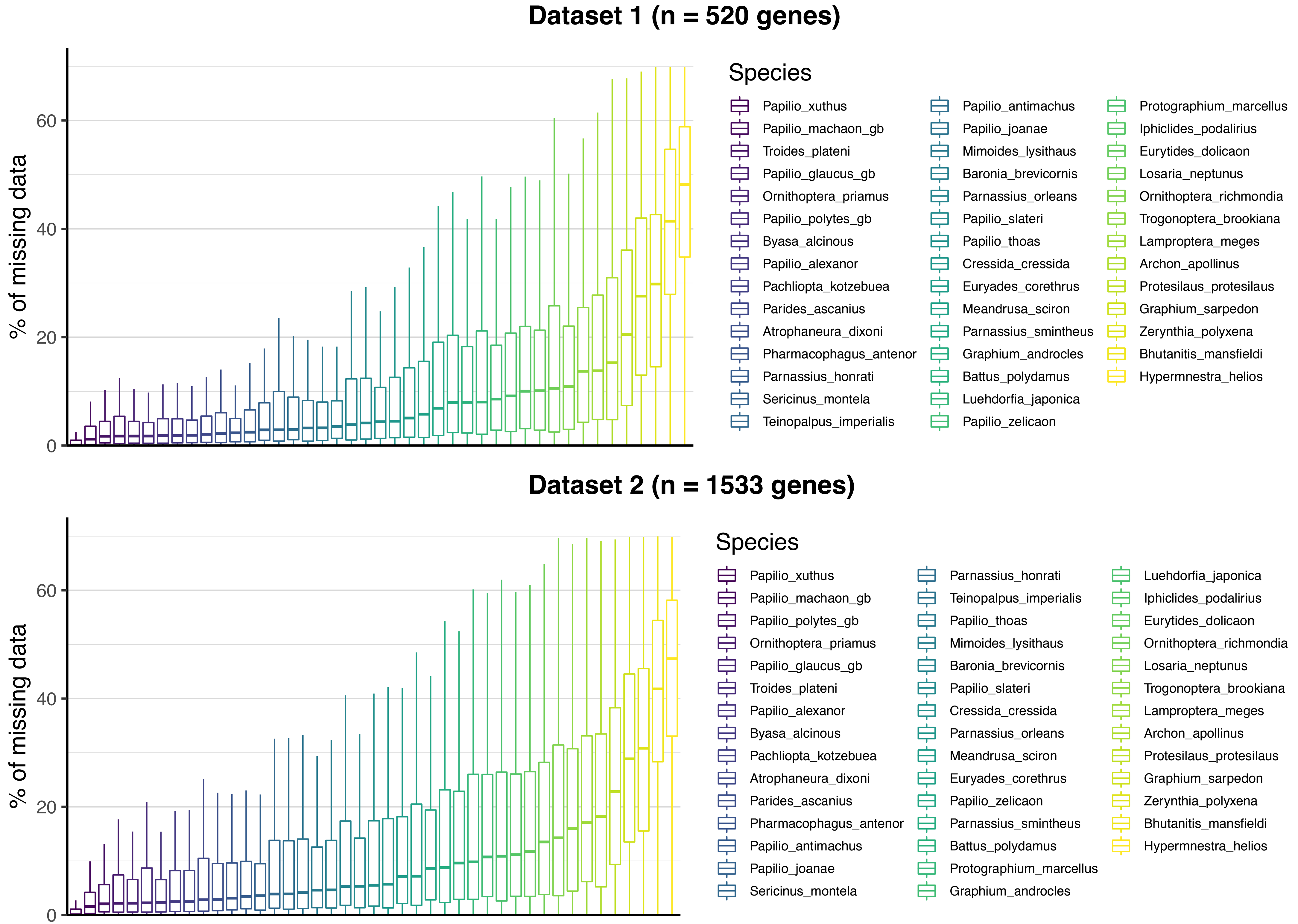
Dataset 1 (n = 520 genes)
% of missing data
0
20
40
60
Species
Papilio_xuthus
Papilio_machaon_gb
Troides_plateni
Papilio_glaucus_gb
Ornithoptera_priamus
Papilio_polytes_gb
Byasa_alcinous
Papilio_alexanor
Pachliopta_kotzebuea
Parides_ascanius
Atrophaneura_dixoni
Pharmacophagus_antenor
Parnassius_honrati
Sericinus_montela
Teinopalpus_imperialis
Papilio_antimachus
Papilio_joanae
Mimoides_lysithaus
Baronia_brevicornis
Parnassius_orleans
Papilio_slateri
Papilio_thoas
Cressida_cressida
Euryades_corethrus
Meandrusa_sciron
Parnassius_smintheus
Graphium_androcles
Battus_polydamus
Luehdorfia_japonica
Papilio_zelicaon
Protographium_marcellus
Iphiclides_podalirius
Eurytides_dolicaon
Losaria_neptunus
Ornithoptera_richmondia
Trogonoptera_brookiana
Lamproptera_meges
Archon_apollinus
Protesilaus_protesilaus
Graphium_sarpedon
Zerynthia_polyxena
Bhutanitis_mansfieldi
Hypermnestra_helios
Dataset 2 (n = 1533 genes)
% of missing data
0
20
40
60
Species
Papilio_xuthus
Papilio_machaon_gb
Papilio_polytes_gb
Ornithoptera_priamus
Papilio_glaucus_gb
Troides_plateni
Papilio_alexanor
Byasa_alcinous
Pachliopta_kotzebuea
Atrophaneura_dixoni
Parides_ascanius
Pharmacophagus_antenor
Papilio_antimachus
Papilio_joanae
Sericinus_montela
Parnassius_honrati
Teinopalpus_imperialis
Papilio_thoas
Mimoides_lysithaus
Baronia_brevicornis
Papilio_slateri
Cressida_cressida
Parnassius_orleans
Meandrusa_sciron
Euryades_corethrus
Papilio_zelicaon
Parnassius_smintheus
Battus_polydamus
Protographium_marcellus
Graphium_androcles
Luehdorfia_japonica
Iphiclides_podalirius
Eurytides_dolicaon
Ornithoptera_richmondia
Losaria_neptunus
Trogonoptera_brookiana
Lamproptera_meges
Archon_apollinus
Protesilaus_protesilaus
Graphium_sarpedon
Zerynthia_polyxena
Bhutanitis_mansfieldi
Hypermnestra_helios

**Supplementary figure 21.** The percentage of missing data ("N" or "-") per branch for the branches with ("Yes", $n$ = 14) and without ("No", $n$ = 14) host-plant shift. For a given branch, the percentage of missing data is the mean value of the species of a clade for which the branch is the root.

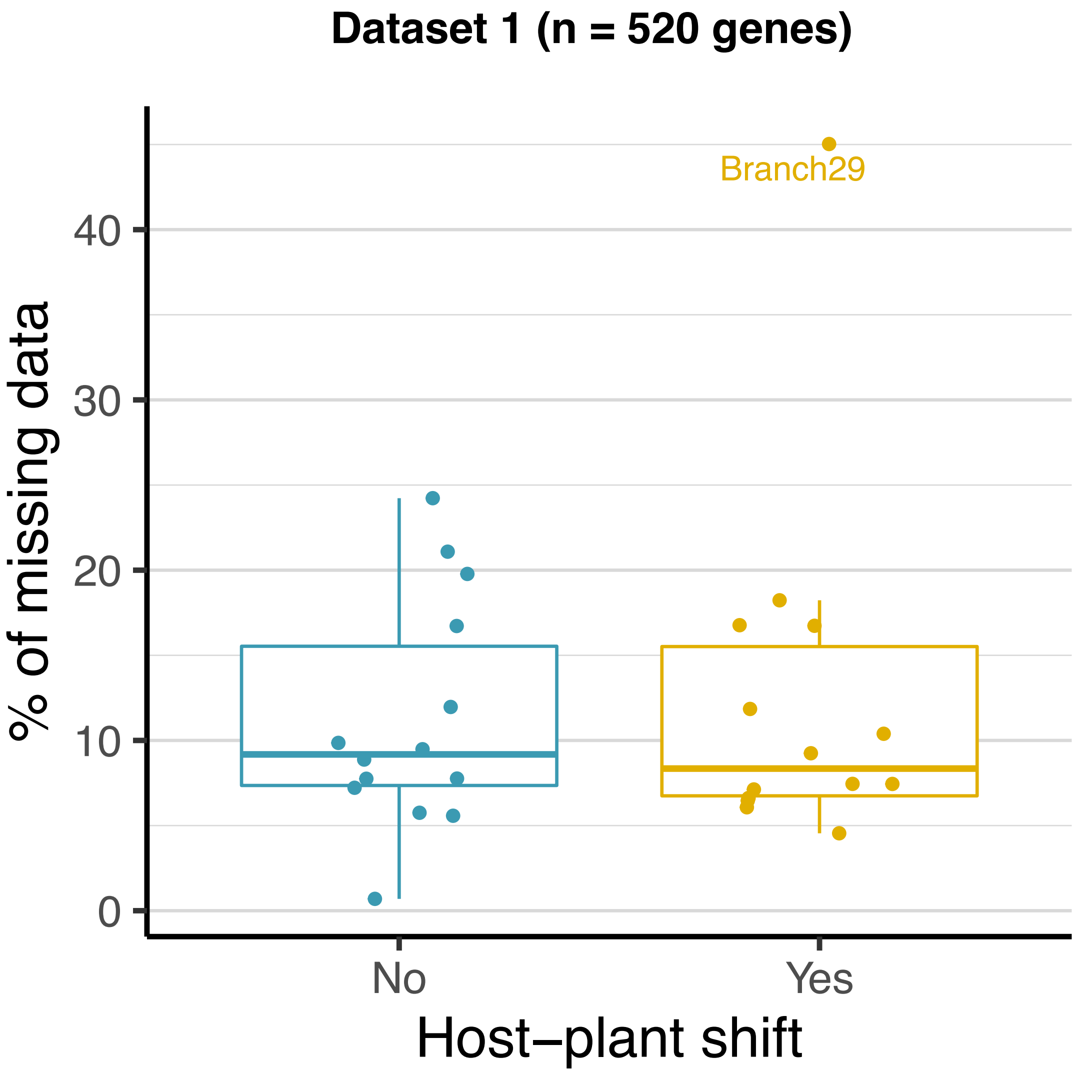
Dataset 1 (n = 520 genes)
Branch29
% of missing data
0
10
20
30
40
No
Yes
Host−plant shift

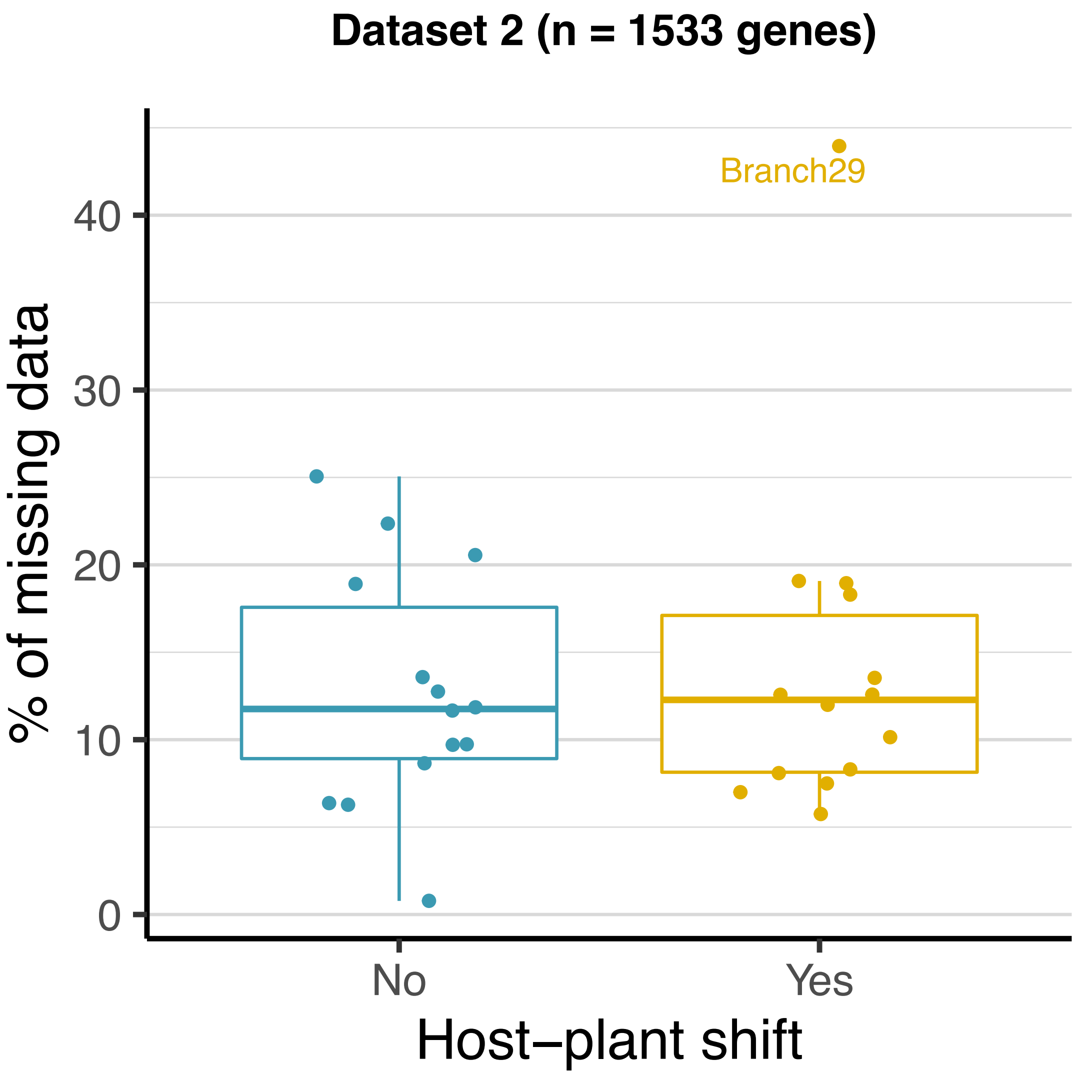
Dataset 2 (n = 1533 genes)
Branch29
% of missing data
0
10
20
30
40
No
Yes
Host−plant shift

**Supplementary figure 22.** Relationship between the percentage of missing data ("N" or "-") and the number of positively selected genes per branch. For a given branch, the percentage of missing data is the mean value of the species of a clade for which the branch is the root.

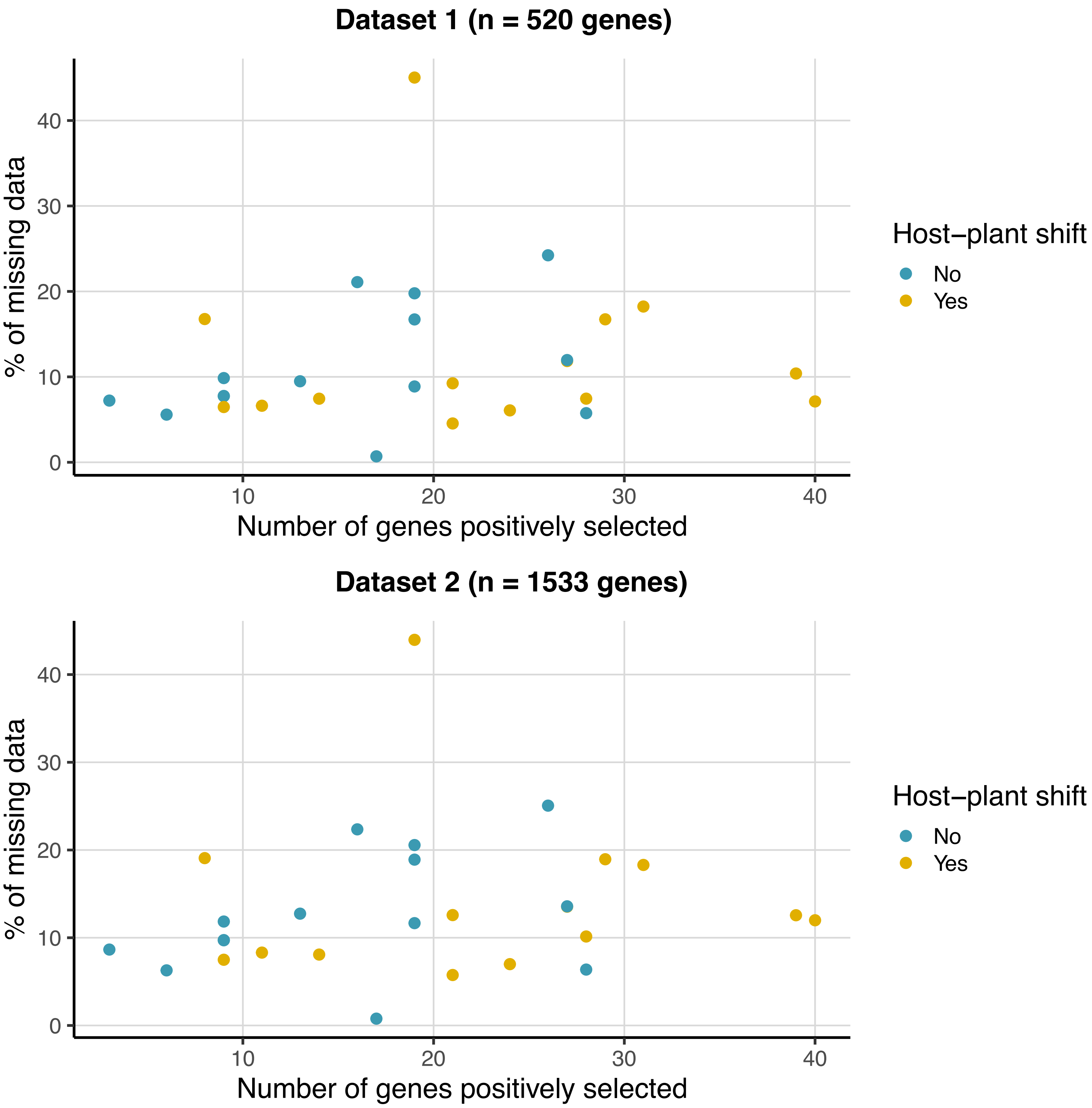
Dataset 1 (n = 520 genes)
% of missing data
0
10
20
30
40
10
20
30
40
Number of genes positively selected
Host−plant shift
No
Yes
Dataset 2 (n = 1533 genes)
% of missing data
0
10
20
30
40
10
20
30
40
Number of genes positively selected
Host−plant shift
No
Yes

**Supplementary figure 23.** The percentage of GC at third codon position (GC3) per gene across species computed for dataset 1 and dataset 2.

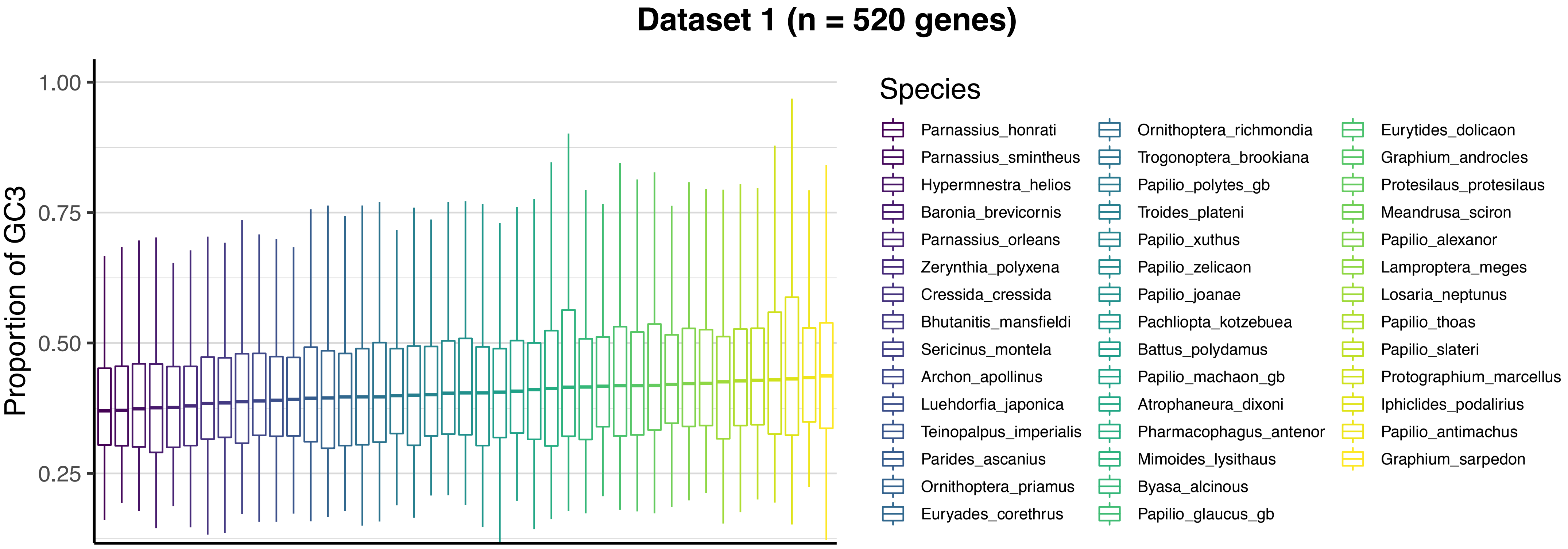
Dataset 1 (n = 520 genes)
Proportion of GC3
1.00
0.75
0.50
0.25
Species
Parnassius_honrati
Parnassius_smintheus
Hypermnestra_helios
Baronia_brevicornis
Parnassius_orleans
Zerynthia_polyxena
Cressida_cressida
Bhutanitis_mansfieldi
Sericinus_montela
Archon_apollinus
Luehdorfia_japonica
Teinopalpus_imperialis
Parides_ascanius
Ornithoptera_priamus
Euryades_corethrus
Ornithoptera_richmondia
Trogonoptera_brookiana
Papilio_polytes_gb
Troides_plateni
Papilio_xuthus
Papilio_zelicaon
Papilio_joanae
Pachliopta_kotzebuea
Battus_polydamus
Papilio_machaon_gb
Atrophaneura_dixoni
Pharmacophagus_antenor
Mimoides_lysithaus
Byasa_alcinous
Papilio_glaucus_gb
Eurytides_dolicaon
Graphium_androcles
Protesilaus_protesilaus
Meandrusa_sciron
Papilio_alexanor
Lamproptera_meges
Losaria_neptunus
Papilio_thoas
Papilio_slateri
Protographium_marcellus
Iphiclides_podalirius
Papilio_antimachus
Graphium_sarpedon

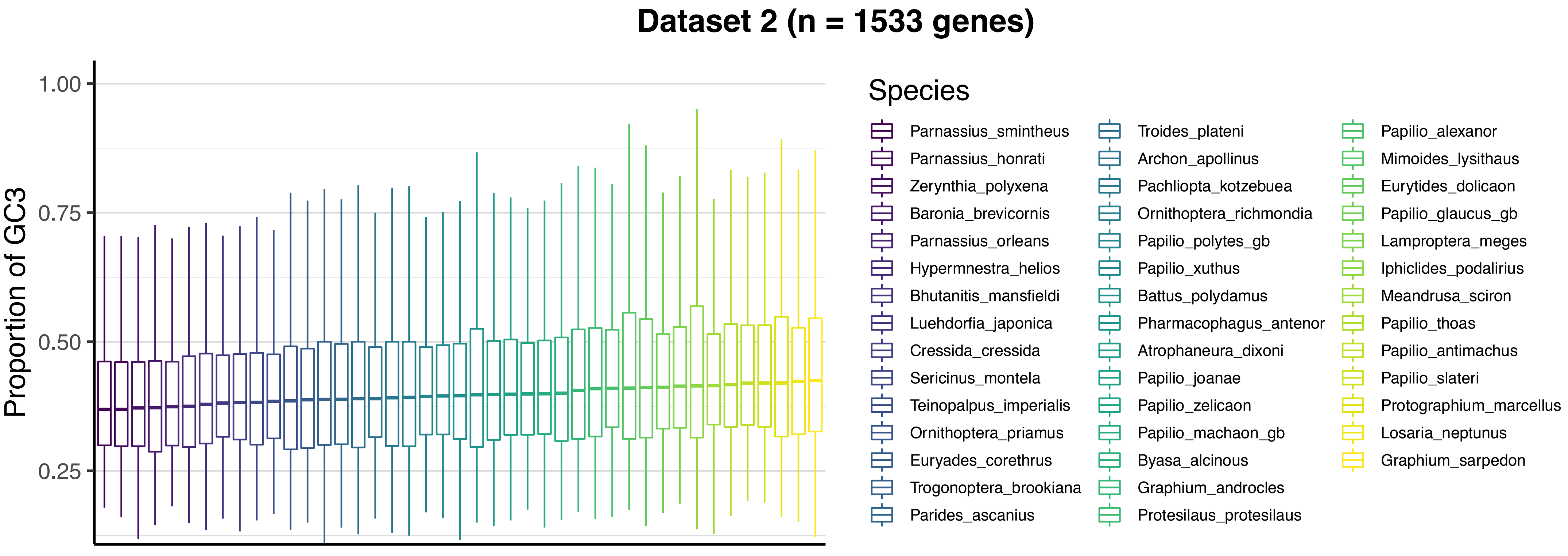
Dataset 2 (n = 1533 genes)
Proportion of GC3
1.00
0.75
0.50
0.25
Species
Parnassius_smintheus
Parnassius_honrati
Zerynthia_polyxena
Baronia_brevicornis
Parnassius_orleans
Hypermnestra_helios
Bhutanitis_mansfieldi
Luehdorfia_japonica
Cressida_cressida
Sericinus_montela
Teinopalpus_imperialis
Ornithoptera_priamus
Euryades_corethrus
Trogonoptera_brookiana
Parides_ascanius
Troides_plateni
Archon_apollinus
Pachliopta_kotzebuea
Ornithoptera_richmondia
Papilio_polytes_gb
Papilio_xuthus
Battus_polydamus
Pharmacophagus_antenor
Atrophaneura_dixoni
Papilio_joanae
Papilio_zelicaon
Papilio_machaon_gb
Byasa_alcinous
Graphium_androcles
Protesilaus_protesilaus
Papilio_alexanor
Mimoides_lysithaus
Eurytides_dolicaon
Papilio_glaucus_gb
Lamproptera_meges
Iphiclides_podalirius
Meandrusa_sciron
Papilio_thoas
Papilio_antimachus
Papilio_slateri
Protographium_marcellus
Losaria_neptunus
Graphium_sarpedon

**Supplementary figure 24.** The percentage of GC at third codon position (GC3) per branch for the branches with (“Yes”, $n$ = 14) and without (“No”, $n$ = 14) host-plant shift. For a given branch, the percentage of GC3 is the mean value of the species of a clade for which the branch is the root.

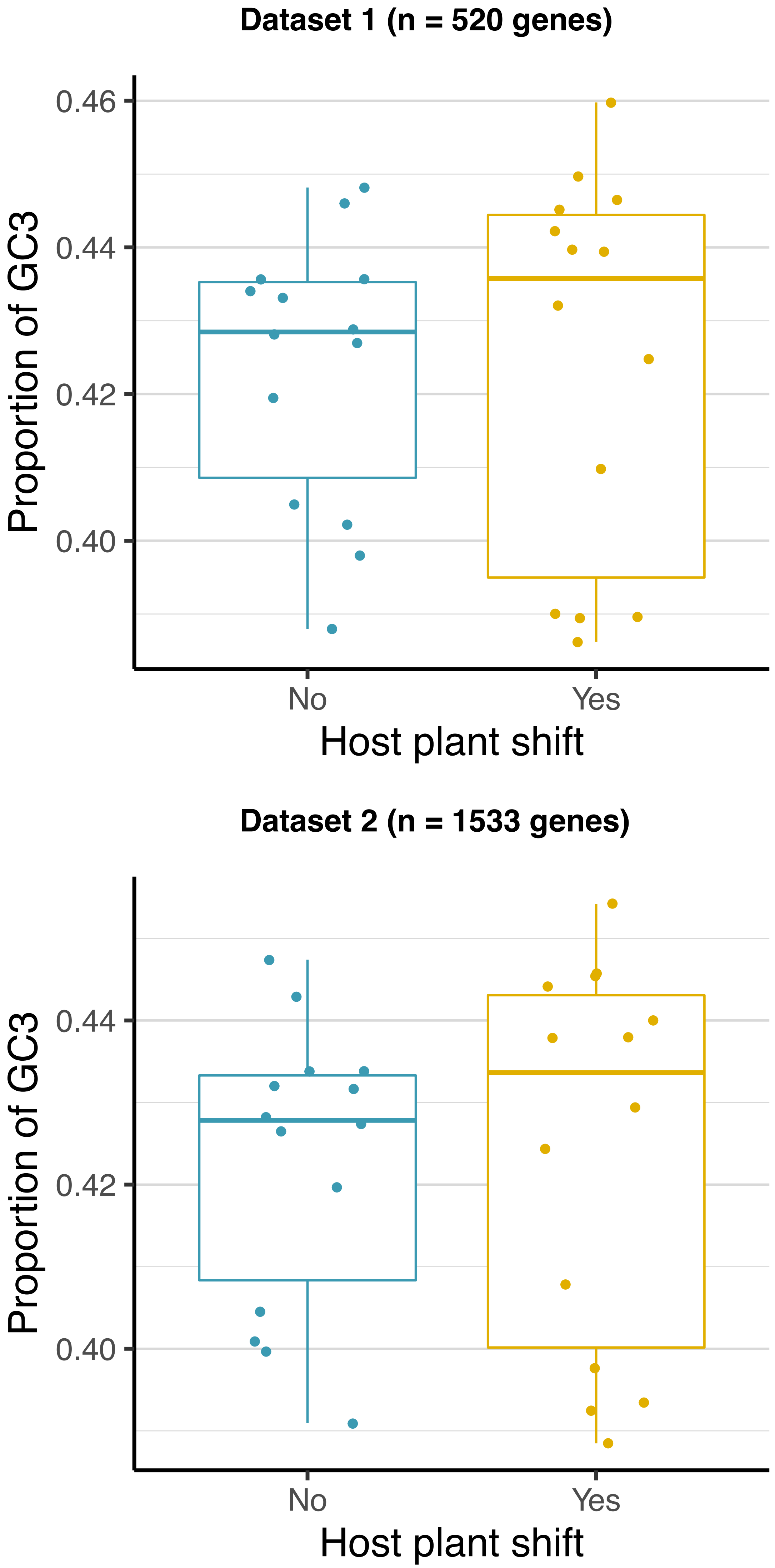

Dataset 1 (n = 520 genes)
0.46
0.44
0.42
0.40
Proportion of GC3
No
Yes
Host plant shift
Dataset 2 (n = 1533 genes)
0.44
0.42
0.40
Proportion of GC3
No
Yes
Host plant shift

**Supplementary figure 25.** Relationship between the percentage of GC at third codon position (GC3) and the number of positively selected genes per branch. For a given branch, the percentage of GC3 is the mean value of the species of a clade for which the branch is the root.

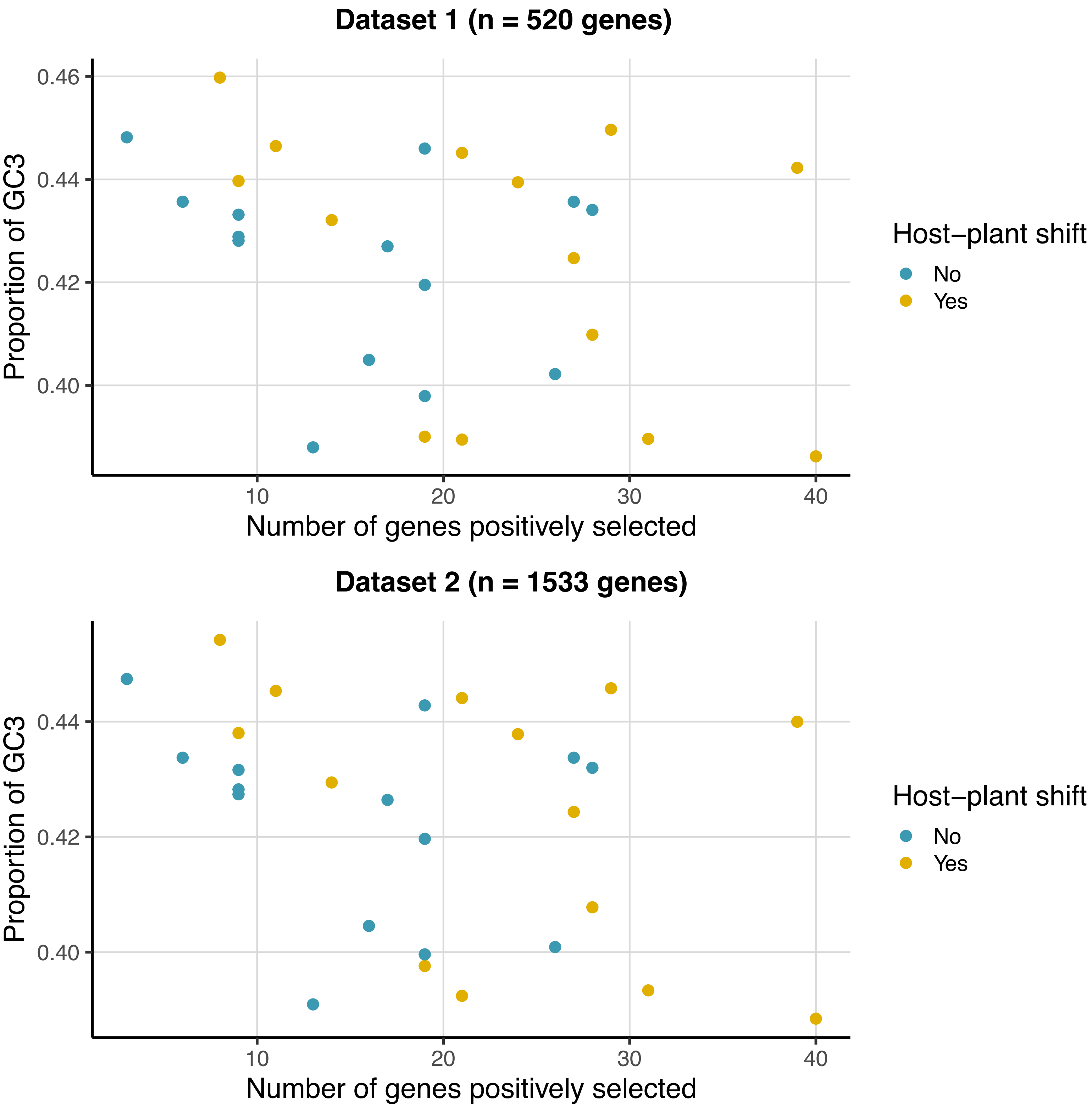
Dataset 1 (n = 520 genes)
Proportion of GC3
0.46
0.44
0.42
0.40
10
20
30
40
Number of genes positively selected
Host−plant shift
No
Yes
Dataset 2 (n = 1533 genes)
Proportion of GC3
0.44
0.42
0.40
10
20
30
40
Number of genes positively selected
Host−plant shift
No
Yes
